# The long-chain fatty acid-CoA ligase FadD1 monitors *Pseudomonas aeruginosa* quorum sensing as a receptor of cis-2-decenoic acid

**DOI:** 10.1101/2024.06.08.598097

**Authors:** Shihao Song, Jingyun Liu, Bing Wang, Yang Si, Hongguang Han, Xiuyun Sun, Mingfang Wang, Binbin Cui, Guangliang Wu, Yongliang Huo, Liangxiong Xu, Beile Gao, Liang Yang, Xiaoxue Wang, Lian-Hui Zhang, Yinyue Deng

**Affiliations:** Key Laboratory of Tropical Biological Resources of Ministry of Education, School of Pharmaceutical Sciences, Hainan University, Haikou 570228, China; Integrative Microbiology Research Center, South China Agricultural University, Guangzhou 510642, China; Zhengzhou Shuqing Medical College, Zhengzhou 450064, China; Department of Microbiology, College of Life Sciences, Nanjing Agricultural University, Nanjing 210095, China; School of Pharmaceutical Sciences (Shenzhen), Shenzhen Campus of Sun Yat-sen University, Sun Yat-sen University, Shenzhen 518107, China; Pharmacy Department, The Affiliated LiHuiLi Hospital of Ningbo University, Ningbo315046, China; Experimental Animal Center, Guangzhou Medical University, Guangzhou, Guangdong, 511436, China; School of Life Sciences, Huizhou University, Huizhou 516007, China; Guangdong Key Laboratory of Marine Materia Medica, South China Sea Institute of Oceanology, Chinese Academy of Sciences, Guangzhou 510301, China; School of Medicine, Southern University of Science and Technology, Shenzhen 518055, China

## Abstract

Diffusible signal factor (DSF)-family quorum sensing (QS) signals are widely utilized by many bacterial pathogens to modulate various biological functions and virulence. Previous studies showed that *cis*-2-decenoic acid (*cis*-DA) is involved in modulation of biofilm dispersion in *Pseudomonas aeruginosa*, but its signaling mechanism remains vague. Here, we report that *cis*-DA regulates the physiology and virulence of *P. aeruginosa* through the long-chain fatty acid-CoA ligase FadD1. *Cis*-DA specifically binds to FadD1 with high affinity and enhances the binding of FadD1 to the promoter DNA region of *lasR*. Further analysis showed that the FadD1 is a response regulator of *cis*-DA with a DNA-binding leucine zipper motif to control the transcription of various target genes. Moreover, FadD1 exhibited catalytic activity on *cis*-2-dodecenoic acid (BDSF) of *Burkholderia cenocepacia* and enhanced the competitiveness of *P. aeruginosa*. Together, our work presents a new DSF-type QS signaling system, which is highlighted by its receptor and response regulator evolved from a canonical enzyme of fatty acid metabolism.

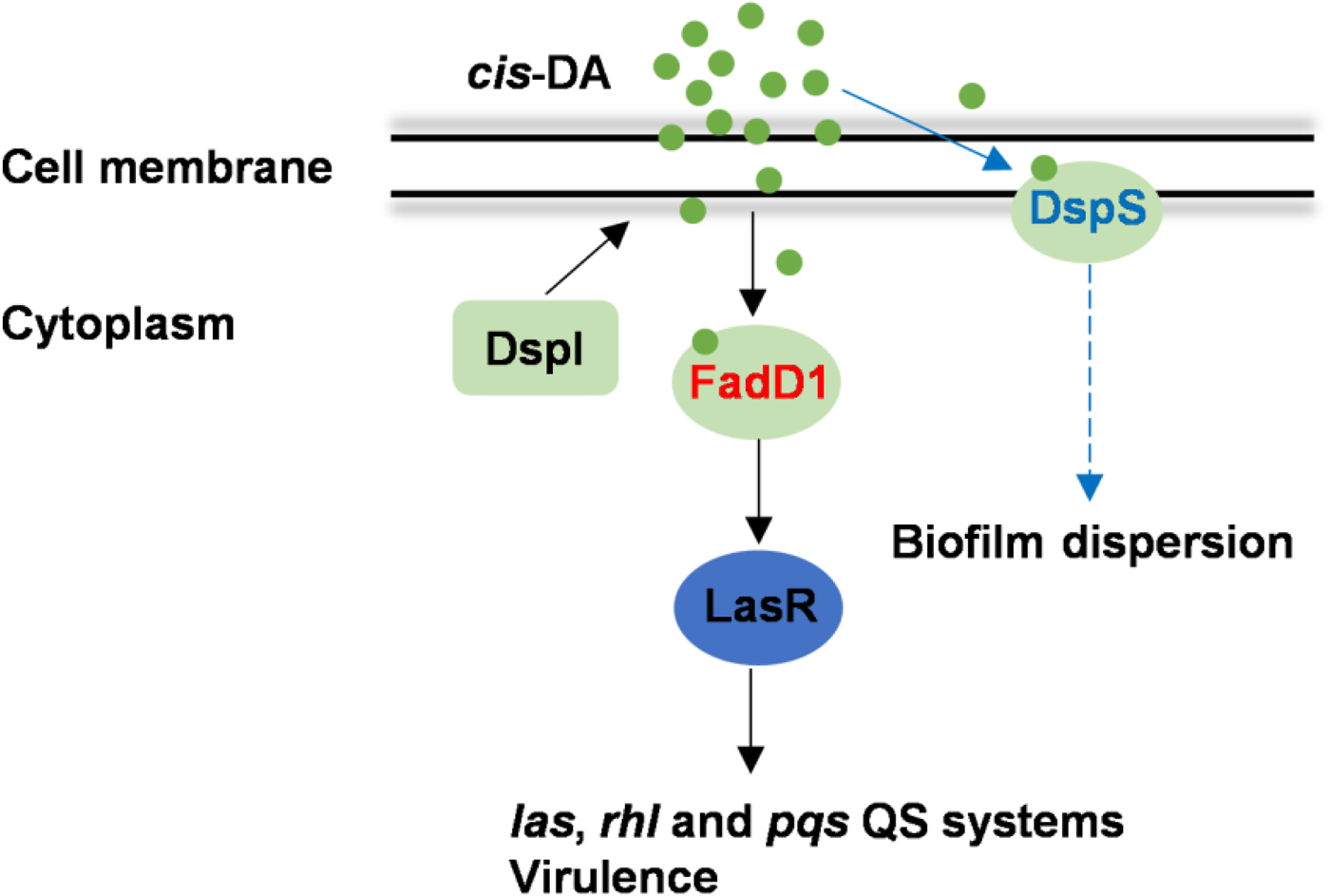

**Highlights:** The long-chain fatty acid-CoA ligase FadD1 acts as a global transcriptional regulator in *P. aeruginosa*. FadD1 is a receptor of QS signal *cis*-DA and functions at the top of the QS hierarchy in *P. aeruginosa*. The homolog of FadD1 in *P. fluorescens* could bind to the target gene promoter by responding to *cis*-DA.

**In brief:** *Pseudomonas aeruginosa* is a human pathogen with antibiotic resistance and a wide range of dynamic defenses makes it an extremely challenging organism to treat in modern-day medicine. It employs *cis*-2-decenoic acid (*cis*-DA) quorum sensing to regulate the physiology and pathogenesis. Song *et al.* demonstrate that the long-chain fatty acid-CoA ligase FadD1 controls the important biological functions and virulence in *P. aeruginosa* as a novel response regulator of *cis*-DA signal.

## INTRODUCTION

Quorum sensing (QS) is a cell-to-cell communication mechanism utilized by numerous species of bacteria. The first QS signal discovered in gram-negative bacteria is the acyl homoserine lactone (AHL) produced by *Vibrio fischeri*^1–5^. In addition to AHL-family signals, there are many other types of QS signals^6–16^, including the diffusible signal factor (DSF)-family signals. *Pseudomonas aeruginosa* is a major source of opportunistic infections in both immunocompromised individuals and cystic fibrosis patients and has evolved at least three types of QS systems, e.g., *las*, *pqs* and *rhl*^17–19^. The *las* and *rhl* AHL QS systems use N-3-oxo-dodecanoyl-l-homoserine lactone (3-oxo-C12-HSL) and N-butyryl-l-homoserine lactone (C4-HSL), respectively, to regulate biological functions, such as biofilm formation, motility and virulence factor production^20–24^. The *pqs* system employs 2-heptyl-3-hydroxy-4(1H)-quinolone (PQS), to regulate biological functions and virulence^25–29^. These QS systems of *P. aeruginosa* are hierarchically interrelated, and the *las* system was verified to monitor both the *rhl* and *pqs* systems^17, 30^. Moreover, it was also revealed that the fatty acid signaling molecule *cis*-2-decenoic acid (*cis*-DA) exhibited important functions in *P. aeruginosa*^31^. The production of *cis*-DA in *P. aeruginosa* needs an enoyl-CoA synthase encoded by DspI^32^. The inactivation of DspI leads to a significant reduction in biofilm dispersion^33,34^. However, it is not yet clear how *cis*-DA regulates biofilm dispersion, and its receptor and downstream signaling network remain to be investigated.

Fatty acyl coenzyme A (CoA) synthetases (FACSs; fatty acid-CoA ligases) are a class of enzymes that activate alkanoic acids to CoA esters^35^. FACSs are widely distributed in both prokaryotic and eukaryotic organisms and exhibit broad substrate specificity^36^. The long-chain fatty acid-CoA ligase (FadD) is responsible for the activation of endogenous long-chain fatty acids into acyl-CoAs^37^. RpfB, which has recently been reported as a long-chain fatty acid-CoA ligase of *Xanthomonas campestris* pv. campestris (*Xcc*), is involved in the activation of a wide range of fatty acids to their CoA esters *in vitro*^38^. Intriguingly, RpfB is required for the turnover of both *cis*-11-methyl-dodecenoic acid (DSF) and *cis-*2-dodecenoic acid (BDSF) QS signals *in vivo*^39^. The *rpfB* deletion mutant produced high levels of both DSF and BDSF signals and displayed increased virulence in *Xcc*^40,41^. Moreover, RpfB-dependent QS signal turnover was also detected in various bacterial species^40^.

Transcription factors (TFs) are proteins that bind to specific DNA sequences, thereby regulating the transcription of genetic information from DNA to messenger RNA^42^. Once bound to DNA, these proteins can promote or block the recruitment of RNA polymerase to specific genes to control transcriptional expression^43^. Although there is great diversity in the structure and function of DNA-binding proteins, some polypeptide motifs enable binding to the major groove of DNA, including helix-turn-helix (HTH), helix-loop-helix (HLH), zinc fingers, and leucine zippers^44–47^.

In this study, we demonstrated that FadD1 of *P. aeruginosa* could convert BDSF to *cis*-2-dodecenoic acid-CoA and reduce the competitiveness of *B. cenocepacia*. FadD1 also controls the physiology and virulence of *P. aeruginosa* by sensing *cis*-DA, then monitors the QS hierarchy and controls the expression of various target genes as a global transcriptional regulator. These findings indicate that FadD1 is a member of a new class of response regulators of QS signal that controls the important biological functions and virulence of bacterial pathogens, with an additional role in fatty acid oxidation to catalyze the formation of fatty acyl-CoA.

## RESULTS

### FadD1 of *P. aeruginosa* shows enzyme activity on the BDSF signal

BDSF was first found to be involved in the regulation of biofilm formation, motility and virulence of *B. cenocepacia*^9–12,48,49^. It was subsequently reported that BDSF from *B. cenocepacia* interferes with QS systems and the type III secretion system (T3SS) of *P. aeruginosa*^50^. Interestingly, we found that BDSF could also be degraded when it was added to the culture of *P. aeruginosa* PAO1. The amount of BDSF in the culture of *P. aeruginosa* decreased to 21.78% after 6 h of incubation (Figure 1A). Previous work demonstrated that RpfB is a long-chain fatty acid-CoA ligase responsible for BDSF and DSF signal turnover in *Xcc*^38,39^. To discover the enzyme that degrades BDSF, we then searched RpfB homologs in the genome of *P. aeruginosa* PAO1 by using the Basic Local Alignment Search Tool (BLAST) algorithm^51^. We found six RpfB homologs, FadD1-6^52^; among them, FadD1 showed the highest homology, with 55.35% identity with RpfB. Therefore, FadD1, which contains 562 aa with a calculated molecular weight of 61.67 kDa, was purified to homogeneity using affinity chromatography (Figure 1B). The *in vitro* enzymatic activity assays showed that when FadD1 was mixed with BDSF, the free BDSF level in the mixture decreased nearly undetectable after 1 h (Figure 1C). These results suggested that FadD1 exhibits enzyme activity on BDSF. Consistently, deletion of *fadD1* in *P. aeruginosa* caused a substantial reduction in BDSF degradation activity (Figure 1D) but did not affect the growth rate of the bacterial cells (Figure S1).

**Figure 1.**
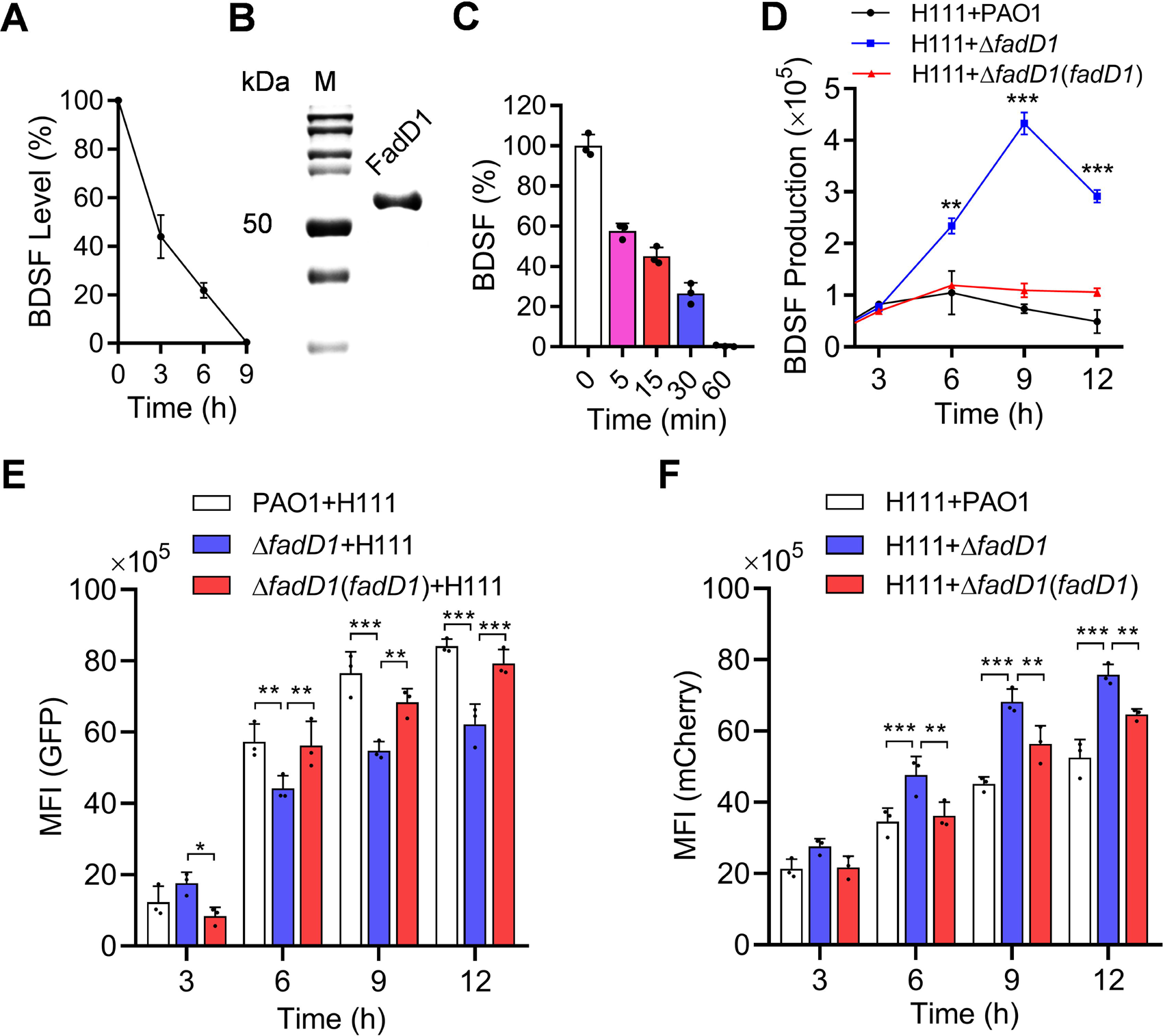
Effect of *fadD1* on the competitive capability of *P. aeruginosa* against *B. cenocepacia*. (A) Analysis of the BDSF signal quenching ability of *P. aeruginosa* PAO1 (n = 3 biological replicates). (B) SDS-PAGE of the purified FadD1 protein. (C) Analysis of the enzymatic quenching activity of FadD1 on BDSF signal (*n* = 3 biological replicates). (D) The BDSF signal production of the *B. cenocepacia* H111 strain cocultured with *P. aeruginosa* PAO1 wild-type, Δ*fadD1* and Δ*fadD1*(*fadD1*) strains (*n* = 3 biological replicates). (E) Green mean fluorescence intensity of the *P. aeruginosa* PAO1 wild-type, Δ*fadD1* and Δ*fadD1*(*fadD1*) strains carrying the green fluorescent protein expression vector (*n* = 3 biological replicates). (F) Red mean fluorescence intensity of the *B. cenocepacia* H111 strain carrying the mCherry fluorescent protein expression vector (*n* = 3 biological replicates). The data are the means ± standard deviations of three independent experiments. The *P. aeruginosa* PAO1 wild-type, Δ*fadD1* and Δ*fadD1*(*fadD1*) strains carried the green fluorescent protein expression vector. The mCherry fluorescent protein expression vector was carried by the *B. cenocepacia* H111 strain. The *P. aeruginosa* PAO1 wild-type, Δ*fadD1* and Δ*fadD1*(*fadD1*) strains were cocultured with *the B. cenocepacia* H111 strain at a ratio of 1:4 (v/v) at OD_600_ = 0.1. The statistical comparisons were performed using two-way ANOVA (\**p* < 0.05; \*\**p* < 0.01; \*\*\**p* < 0.001).

We then investigated the *in vivo* activity of FadD1 in *B. cenocepacia* H111 by overexpressing the *fadD1* gene. Overexpression of *fadD1* did not affect the growth rate but markedly reduced the BDSF production of the bacterial cells (Figure S2A, B). As expected, deletion of *rpfF_Bc_* resulted in complete abolishment of the production of BDSF in *B. cenocepacia* H111 (Figure S2B). Interestingly, overexpression of *fadD1* in the *B. cenocepacia* wild-type strain reduced biofilm formation, swarming motility and protease activity, which could be restored by the addition of exogenous BDSF at 5 µM (Figure S2C-E). Overexpression of *fadD1* in the wild-type strain H111 also decreased cytotoxicity (Figure S2F).

### FadD1 boosts the competitive capability of *P. aeruginosa* against *B. cenocepacia*

As BDSF mediates the cross-talk between *B. cenocepacia* and *P. aeruginosa* ^50^ and FadD1 exhibits catalysis activity on BDSF, we investigated whether FadD1 plays a role in the competitive interactions between the two bacterial species. To this end, we cocultured the green fluorescent protein (GFP) fluorescence-labeled *P. aeruginosa* PAO1 wild-type, Δ*fadD1* and complemented strains in the presence of the *B. cenocepacia* H111 strain labeled with mCherry and measured the BDSF production of *B. cenocepacia* (Figure 1D) and the mean fluorescence intensity (MFI) of both *B. cenocepacia* and *P. aeruginosa* at different time points as indicated (Figure 1E, F). The results showed that deletion of *fadD1* caused a drastic reduction in the degradation capability of *P. aeruginosa* toward the BDSF signal. The BDSF signal yield of *B. cenocepacia* cocultured with the *P. aeruginosa* deletion mutant Δ*fadD1* was more than 5 times higher than that when it was cocultured with wild-type PAO1 or the complemented strain Δ*fadD1*(*fadD1*) at 9 h postinoculation (Figure 1D). On the other hand, we found that the GFP mean fluorescence intensity (MFI) of the *P. aeruginosa* Δ*fadD1* strain was lower than that of the *P. aeruginosa* wild-type and complemented strains when they were cocultured with the *B. cenocepacia* strains (Figure 1E). The mCherry mean fluorescence intensity (MFI) of the *B. cenocepacia* H111 strain cocultured with the *P. aeruginosa* Δ*fadD1* strain was higher than that of the *B. cenocepacia* H111 strain cocultured with the *P. aeruginosa* wild-type and complemented strains (Figure 1F). These results indicated that FadD1 plays an important role in the competition between *P. aeruginosa* and *B. cenocepacia*.

### Deletion of FadD1 impairs biological functions and pathogenicity in *P. aeruginosa*

It was determined that FadD1 plays an important role in the interaction between *P. aeruginosa* and *B. cenocepacia*. We continued to study whether FadD1plays a role in the regulation of biological functions and pathogenesis in *P. aeruginosa*. We found that deletion of *fadD1* resulted in decreases in biofilm formation, swarming, and pyocyanin production by 30.47%, 36.72% and 62.15%, respectively (Figure 2A-C). Deletion of *fadD1* also attenuated cytotoxicity by 69.78% when A549 cells were incubated with the *P. aeruginosa* strains at 8 h postinoculation (Figure 2D). We tested whether there was a change in the expression level of upstream (*PA3298*) and downstream (*fadD2*) genes of the *fadD1* mutant gene. The results showed that there was no change in the expression of upstream and downstream genes in the *fadD1* deletion mutant (Figure S3). To further study the regulatory roles of FadD1 in controlling bacterial physiology, we analyzed and compared the transcriptomes of the wild-type strain and the Δ*fadD1* mutant strain by using RNA sequencing (RNA-Seq). Differential gene expression analysis showed that the expression levels of 23 genes were increased and those of 208 genes were decreased in the Δ*fadD1* mutant strain compared with the expression levels in the wild-type strain (|Log_2_ fold-change| ≥ 1.0) (Table S1 and Figure S4A). These differentially expressed genes are associated with a range of biological functions (Table S1 and Figure S4B). These genes include the *las*, *rhl* and *pqs* QS system genes *PA1432* (*lasI*), *PA1430* (*lasR*), *PA3476* (*rhlI*), *PA3477* (*rhlR*), *PA0996* (*pqsA*), and *PA1003* (*mvfR*), which were shown to be involved in pathogenicity in *P. aeruginosa*.

**Figure 2.**
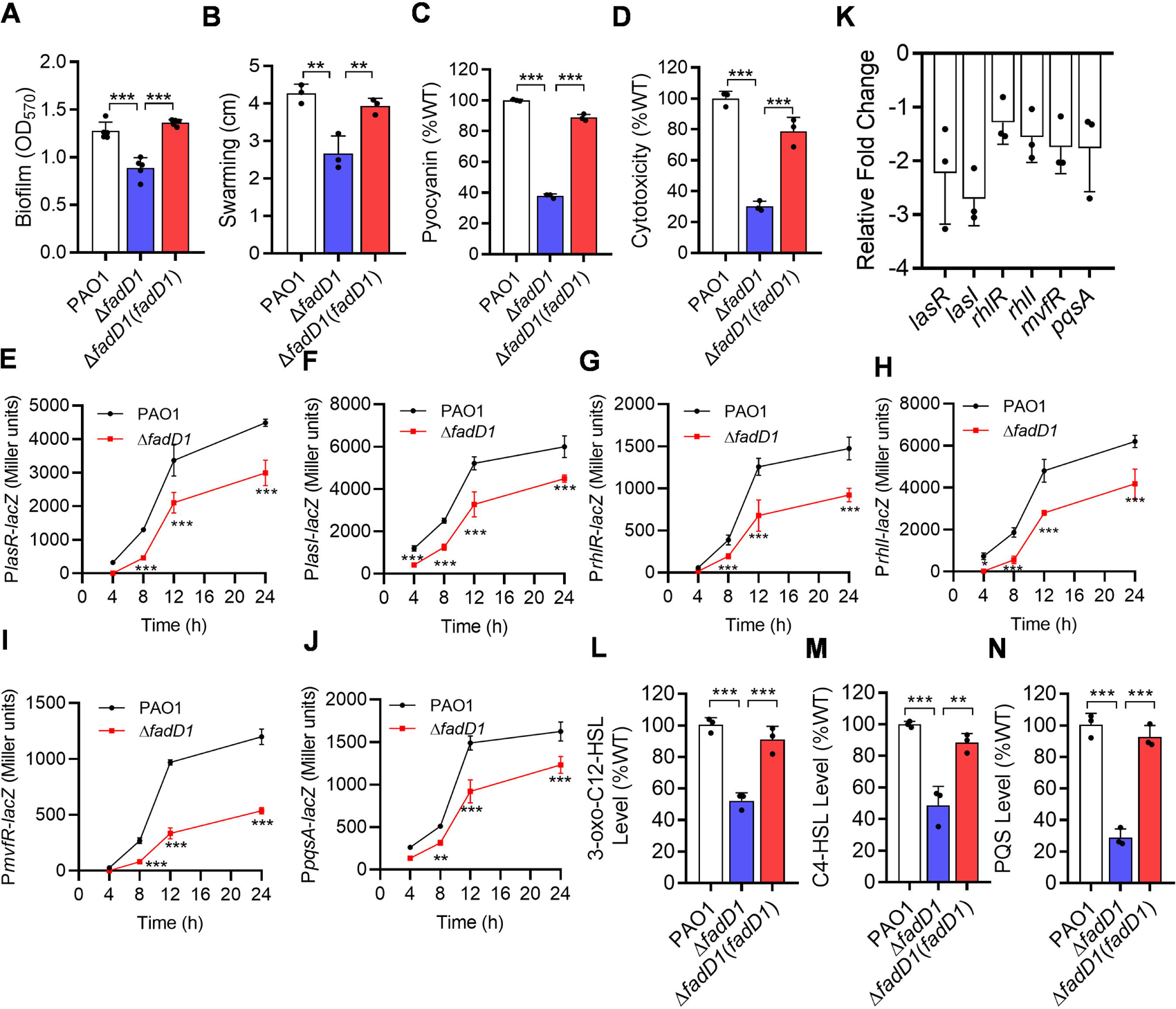
Effects of *fadD1* on the QS-regulated phenotypes and QS of *P. aeruginosa*. Analysis of biofilm formation (A), swarming motility (B), pyocyanin production (C), and virulence (D) in *P. aeruginosa* PAO1((A), *n* = 5 biological replicates; (B), (C), (D), *n* = 3 biological replicates). Effects of *fadD1* on the gene expression levels of *lasR* (E), *lasI* (F), *rhlR* (G), *rhlI* (H), *mvfR* (I) and *pqsA* (J) by assessing the β-galactosidase activity of the promoter-*lacZ* transcriptional fusions and by RT-qPCR (K), which are the signal molecule synthase-encoding genes and receptor genes of the *rhl*, *las* and *pqs* QS systems in *P. aeruginosa* PAO1, respectively ((E), (F), (G), (H), (I), (J), *n* = 6 biological replicates; (K), *n* = 3 biological replicates). The production of 3-oxo-C12-HSL (L), C4-HSL (M) and PQS (N) in the *P. aeruginosa* PAO1 wild-type, Δ*fadD1* and Δ*fadD1*(*fadD1*) strains (*n* = 3 biological replicates). The production of QS signals in the *P. aeruginosa* PAO1 wild-type strain was arbitrarily defined as 100%. The data are means ± standard deviations of three independent experiments. The statistical comparisons were performed using one-way ANOVA or two-way ANOVA ((A), (B), (C), (D), (L), (M), (M), one-way ANOVA; (E), (F), (G), (H), (I), (J), two-way ANOVA; \**p* < 0.05; \*\**p* < 0.01; \*\*\**p* < 0.001).

### FadD1 monitors the QS systems of *P. aeruginosa*

As FadD1 obviously affects the QS-regulated genes and phenotypes of *P. aeruginosa* Figure 2A-D, Figure S4 and Table S1), we inferred that FadD1 might affect the QS systems. Analysis of the expression profiles of *lacZ* under the control of the *lasR*, *lasI*, *rhlR*, *rhlI*, *mvfR* and *pqsA* promoters showed that deletion of *fadD1* resulted in reduced expression levels of *lasR*, *lasI*, *rhlR*, *rhlI*, *mvfR* and *pqsA* (Figure 2E-J), which are signal molecule receptor genes and synthase-encoding genes of the *las*, *rhl* and *pqs* QS systems, respectively. RT-qPCR analysis and RNA-seq also showed that the mutation of *fadD1* caused a decrease in the expression levels of *lasR*, *lasI*, *rhlR*, *rhlI*, *mvfR* and *pqsA* (Figure 2K). Therefore, we measured and compared the production of 3-oxo-C12-HSL, C4-HSL and PQS in the wild-type, *fadD1* mutant, and complemented strains. We found that the production of 3-oxo-C12-HSL, C4-HSL and PQS was reduced in the *fadD1* mutant strain (Figure 2L-N). These results demonstrated that FadD1 positively regulated the *las*, *rhl* and *pqs* QS systems in *P. aeruginosa*.

### FadD1 regulates target gene expression by directly binding to the promoter

FadD1 obviously affects the QS gene expression levels and QS signal production of *P. aeruginosa*. Therefore, we performed electrophoretic mobility shift assays (EMSAs) to test whether the transcriptional regulation of the signal molecule synthase-encoding genes and receptor genes of the *las*, *rhl* and *pqs* QS systems is achieved by direct binding of FadD1 to their promoters. The 339-bp, 321-bp, 270-bp, 330-bp, 306-bp and 324-bp DNA fragments from the *lasR*, *lasI*, *rhlR*, *rhlI*, *mvfR* and *pqsA* promoters were PCR-amplified and used as probes. Surprisingly, as shown in Figure. 3A, the *lasR* promoter DNA fragment formed stable DNA-protein complexes with FadD1 and migrated slower than unbound probes. The amount of labeled probe that bound to FadD1 increased with increasing amounts of FadD1 but decreased in the presence of a 50-fold greater concentration of unlabeled probe (Figure 3A). However, FadD1 did not bind to the promoters of the other tested genes, i.e., *rhlR*, *mvfR*, *lasI*, *rhlI* and *pqsA* (Figure S5 and Figure S6).

**Figure 3.**
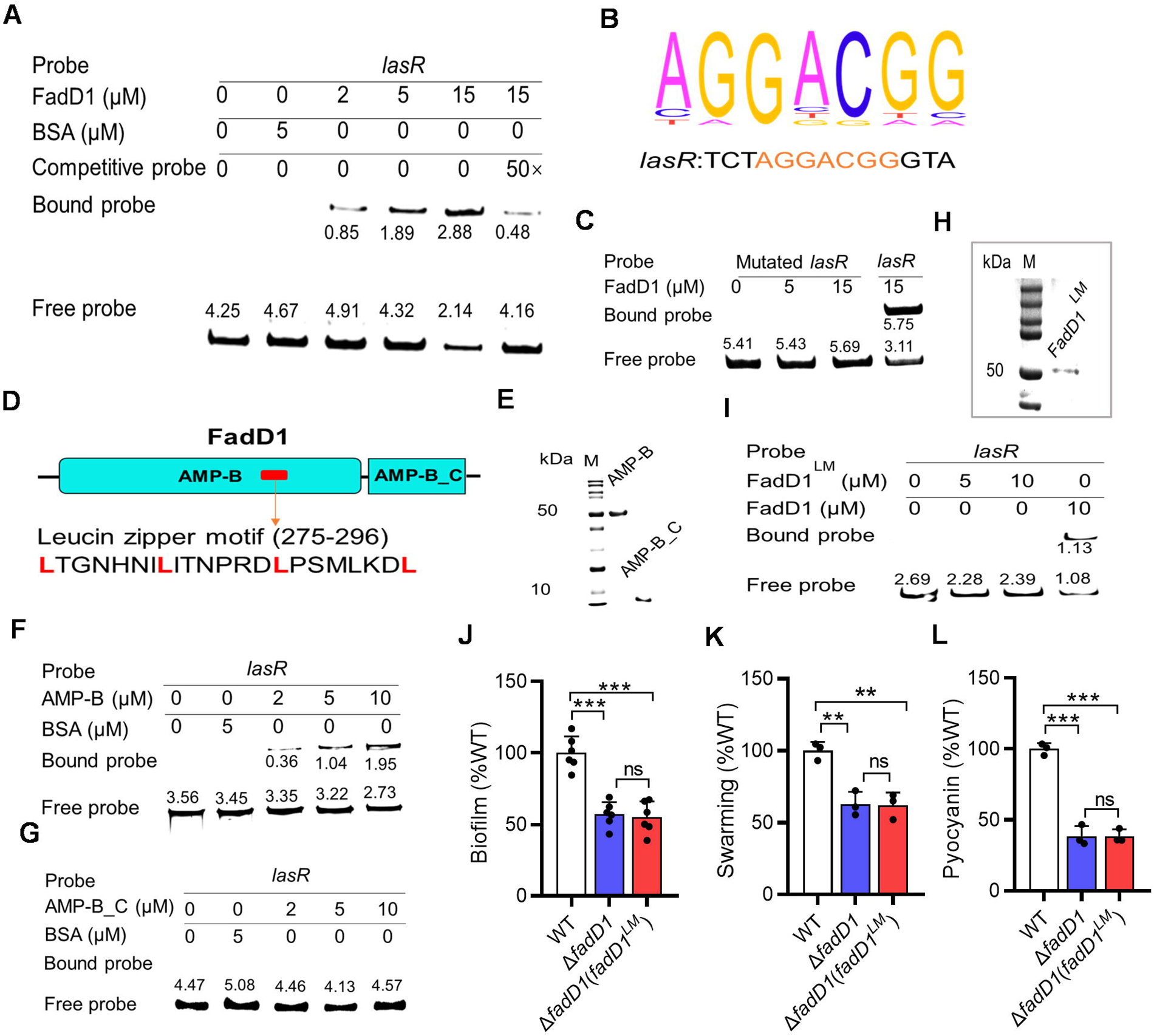
Analysis of the binding of FadD1 to target gene promoters. (A) EMSA analysis of *in vitro* binding of FadD1 to the promoter of *lasR*, in which the biotin-labeled 339-bp *lasR* promoter DNA probe was used for the protein binding assay. A protein-DNA complex, represented by a band shift, was formed when different concentrations of FadD1 protein were incubated with the probe at room temperature for 30 min. (B) A potential FadD1 binding region was identified by ChIP-seq. The conserved sequence is shown in orange. (C) Analysis of the binding between FadD1 and the mutated *lasR* promoter with deletion of the FadD1 binding sequence AGGACGG. EMSA analysis was performed *in vitro*. (D) Domain structure analysis of FadD1 (https://www.ebi.ac.uk/Tools/hmmer/). The red box represents the leucine zipper motif (https://www.novopro.cn/tools/motifscan.html) in AMP-B domain. (E) SDS-PAGE of the purified AMP-B domain and the AMP-B_C domain of the FadD1 protein. EMSA analysis of *in vitro* binding of the AMP-B domain (F) and the AMP-B_C domain (G) to the promoter of *lasR*. (H) SDS-PAGE of the purified mutated FadD1 protein (FadD1^LM^: L275A L282A L289A L296A). (I) EMSA analysis of the *in vitro* binding of the mutated FadD1 protein to the promoter of *lasR*. *In trans* expression of the mutated *fadD1* could not restore biofilm formation (J), swarming motility (K), and pyocyanin production (L) in the *P. aeruginosa* Δ*fadD1* strain ((J), *n* = 6 biological replicates; (K), (L), *n* = 3 biological replicates). The EMSA experiments were performed three times, and representative images from one experiment are shown. The data are the means ± standard deviations of three independent experiments. The statistical comparisons were performed using one-way ANOVA (\*\**p* < 0.01; \*\*\**p* < 0.001; ns = no significance).

To identify the DNA motif recognized by FadD1 and the genes directly regulated by FadD1, we performed chromatin immunoprecipitation sequencing (ChIP-Seq) analysis. ChIP-Seq data analysis revealed the potential FadD1 binding site as 5’-AGGACGG-3’ in the *lasR* gene promoter probe (Figure 3B). The results showed that FadD1 could directly bind to multiple target gene promoters to regulate their transcription and expression (Table S2). To further examine the binding sites, three target genes, *flgM*, *katA* and *PA0692,* were chosen for EMSA analysis. The amounts of labeled probes of *flgM*, *katA* and *PA0692* bound to FadD1 were also increased with increasing amounts of FadD1 (Figure S7). Then, the potential binding sites 5’-AGGACGG-3’, 5’-TGGCCGG-3’, 5’-AGGAGGG-3’, and 5’-AGGGCGG-3’ were deleted from the promoter regions of *lasR*, *flgM*, *katA* and *PA0692*, respectively. EMSA analysis showed that no DNA-protein complex was formed when the binding site was deleted from the probes (Figure 3C and Figure S8), indicating that this fragment is essential for the binding of FadD1 to the *lasR, flgM*, *katA* and *PA0692* promoters. Together, these results demonstrated that FadD1 regulates target gene expression by directly binding to a specific region in the gene promoter.

### The leucine zipper motif of FadD1 is responsible for binding with DNA

Since FadD1 can bind to *lasR* and regulate transcription, and the domain structure analysis of FadD1 by using the HMMER (https://www.ebi.ac.uk/Tools/hmmer/) shows that it has an AMP-binding enzyme domain (AMP-B) and an AMP-binding enzyme C-terminal domain (AMP-B_C) (Figure 3D). We then purified the AMP-B domain and the AMP-B_C domain for EMSA (Figure 3E). The results showed that only the AMP-B domain can bind to the promoter region of *lasR* (Figure 3F, G). To determine the location of the new DNA binding motif in the AMP-B domain, we used the protein motif analysis website (https://www.novopro.cn/tools/motifscan.html). A structure called “leucine zipper” was characterized at 275 to 296 amino acids **L**TGNHNI**L**ITNPRD**L**PSMLKD**L** in AMP-B domain (Figure 3D and Figure S9). The leucine zipper consists of a periodic repetition of leucine residues at every seventh position over a distance covering eight helical turns. This motif is found in many eukaryotic transcription factors^45^. However, in bacterial transcription factor, this motif is only found in the classical DNA-binding HTH domain of the LysR family regulator MetR^46^ and a BDSF response regulator DsfR identified recently^47^. So we speculated that the four leucine residues L275, L282, L289 and L296 might be important for the binding of FadD1 to target gene promoters. We then introduced four site-specific substitutions into FadD1 (named FadD1^LM^), and the results showed that the simultaneous mutation of L275A, L282A, L289A and L296A abolished the binding of FadD1 to the *lasR* promoter (Figure 3I). In addition, *in trans* expression of FadD1^LM^ did not rescue the defective biofilm, swarming motility and pyocyanin production phenotypes of the Δ*fadD1* strain (Figure 3J-L). We then used the HDOCK server to automatically predict the interaction between FadD1 and the DNA-binding site 5’-AGGACGG-3’. The binding model of DNA and FadD1 and the detailed contacts of the complex are shown in Figure S10 and Table S3, respectively. The binding sites of FadD1 are in the leucine zipper structure. The results are consistent with the conclusion that the DNA-binding motif of FadD1 is the leucine zipper.

### FadD1 is a receptor of *cis*-DA

Since *cis*-DA is a DSF-type QS signal in *P. aeruginosa*, and the FadD1 controls QS-regulated phenotypes as a global transcriptional regulator in *P. aeruginosa*, we then measured whether there is an interaction between *cis*-DA and FadD1. To test this hypothesis, we purified FadD1 to performed microscale thermophoresis (MST) analysis to test whether FadD1 binds *cis*-DA. As shown in Figure 4A, FadD1 exhibited binding activity to *cis*-DA, with an estimated dissociation constant (*K_D_*) of 1.06 ± 0.14 μM. To further confirm FadD1 is a receptor of *cis*-DA, we generated the *cis*-DA deficient mutant Δ*dspI*, the double deletion mutants Δ*dspI*Δ*fadD1* and *in trans* expressed *fadD1* in the Δ*dspI* and measured the motility activity and pyocyanin of these strains. The results showed that *in trans* expression of *fadD1* rescued the defective motility and pyocyanin phenotypes of the Δ*dspI* strain (Figure 4B, C). Interestingly, we found that the addition of exogenous *cis*-DA rescued the impaired motility and pyocyanin of Δ*dspI* (Figure 4B, C); however, addition of exogenous *cis*-DA showed no effect on these phenotypes of the mutant Δ*fadD1* and Δ*dspI*Δ*fadD1* (Figure 4B, C). We then measured the gene expression levels of *lasR*, *lasI*, *rhlR*, *rhlI*, *mvfR* and *pqsA* in the wild-type strain and the Δ*dspI* strain. The results indicated that deletion of *dspI* caused a significant decrease in the gene expression levels of *las*, *rhl* and *pqs* QS system genes (Figure 4D-I). Therefore, we measured and compared the production of 3-oxo-C12-HSL, C4-HSL and PQS in the wild-type, *dspI* mutant, and complemented strains. The result showed that the production of 3-oxo-C12-HSL, C4-HSL and PQS was reduced in the *dspI* mutant strain (Figure 4J-L). These results demonstrated that both *cis*-DA and FadD1 positively regulates the *las*, *rhl* and *pqs* QS systems in *P. aeruginosa*.

**Figure 4.**
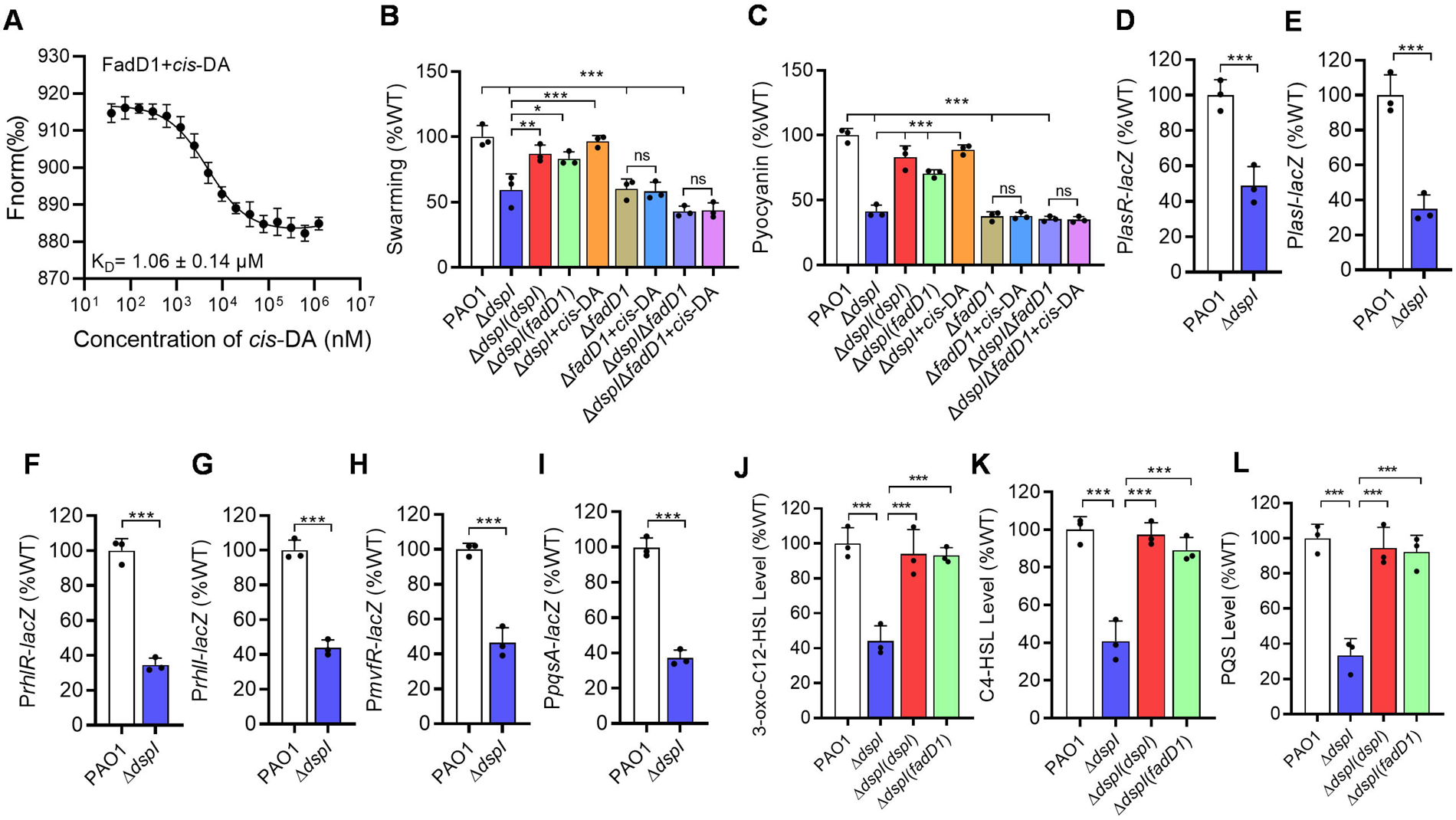
Effects of FadD1 on *cis*-DA-regulated phenotypes of *P. aeruginosa*. (A) MST analysis of the binding of *cis*-DA to the FadD1. “Fnorm (‰)” indicates the fluorescence time trace changes in the MST response. Complementation of the *dspI* mutant with *fadD1*. *In trans* expression of *fadD1* complemented swarming motility (B) and pyocyanin production (C) in the *dspI*-deficient mutant. Effects of *dspI* on the gene expression levels of *lasR* (D), *lasI* (E), *rhlR* (F), *rhlI* (G), *mvfR* (H), *pqsA* (I) and the production of 3-oxo-C12-HSL (J), C4-HSL (K) and PQS (L) of the *P. aeruginosa* PAO1 strains. The production of QS signals in the *P. aeruginosa* PAO1 wild-type strain was arbitrarily defined as 100%. *n* = 3 biological replicates. The data are means ± standard deviations of three independent experiments. The statistical comparisons were performed using one-way ANOVA (\**p* < 0.05; \*\**p* < 0.01; \*\*\**p* < 0.001).

### *cis*-DA enhances the binding of FadD1 to target gene promoters

To determine how the binding of *cis*-DA to FadD1 might affect the activity of FadD1, we examined the effects of *cis*-DA on the binding of FadD1 to the *lasR* promoter by EMSA. As shown in Figure 5A, the binding of FadD1 to the promoter probe of *lasR* was enhanced when *cis*-DA was present in the reaction mixtures, and the amount of probe bound to FadD1 increased as the amount of *cis*-DA increased (Figure 5A).

**Figure 5.**
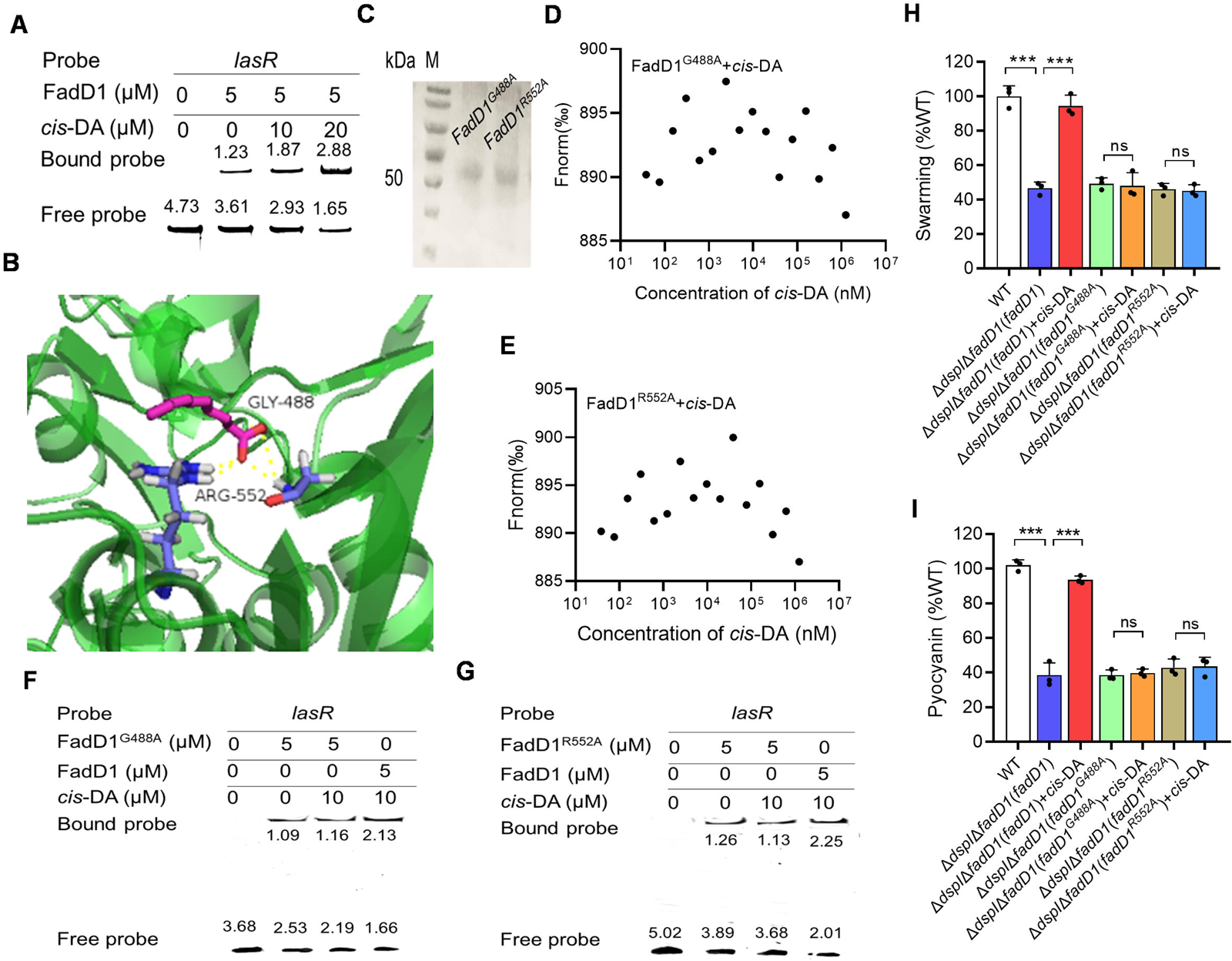
Effects of *cis*-DA on the binding of FadD1 to the promoters of the target genes. (A) EMSA analysis of the *in vitro* binding of FadD1 to the promoter of *lasR* with the addition of different amounts of *cis*-DA. (B) The potential residues (which are shown as stick models and are labeled) of FadD1 involved in *cis*-DA binding analyzed by AutoDocking software. The hydrogen bonds were represented by yellow dashed lines. (C) SDS‒PAGE gel electrophoresis of purified FadD1 variants. MST analysis of the binding of BDSF to the (D) FadD1^G488A^ and (E) FadD1^R552A^. EMSA analysis of the *in vitro* binding of (F) FadD1^G488A^ and (G) FadD1^R552A^ to the promoters of *lasR* with the addition of different amounts of *cis*-DA. The EMSA experiments were performed three times, and representative images from one experiment are shown. Analysis of the perception of *cis*-DA by FadD1^G488A^ and FadD1^R552A^, *cis*-DA-regulated phenotypes: (H) swarming motility and (I) pyocyanin production in *P. aeruginosa* (*n* = 3 biological replicates). The data are means ± standard deviations of three independent experiments. The statistical comparisons were performed using one-way ANOVA (\*\*\**p* < 0.001; ns = no significance).

Then, we attempted to identify the *cis*-DA-binding sites in FadD1. Autodocking analysis revealed 2 amino acid residues, Gly488 (G488) and Arg552 (R552), that might be critical for the interaction between FadD1and *cis*-DA (Figure 5B). To test whether the roles of these acid residues are related to the binding to *cis*-DA, we then generated two single point mutants (FadD1^G488A^ and FadD1^R552A^) (Figure 5C). MST analysis showed that the mutant FadD1^G488A^ and FadD1^R552A^ did not bind to *cis*-DA (Figure 5D, E). Consistent with this, EMSA analysis showed that mutation of the *cis*-DA-binding site (Gly488 and Arg552) completely eliminated the effect of *cis*-DA on FadD1 (Figure 5F, G). In addition, *in trans* expression of *fadD1^G488A^* and *fadD1^R552A^* only partially rescued motility and pyocyanin (Figure 5H, I), and the addition of exogenous 20 μM *cis*-DA showed no effect on these phenotypes of the mutant Δ*dspI*Δ*fadD1*(*fadD1^G488A^*) and Δ*dspI*Δ*fadD1*(*fadD1^R552A^*) (Figure 5H, I). Thus, we concluded that FadD1 is a specific response regulator of *cis*-DA in *P. aeruginosa*.

### Reconstruction of FadD2 with the leucine zipper motif

Our study found that the DNA binding motif of FadD1 is a leucine zipper motif. Therefore, we used the protein motif analysis website (https://www.novopro.cn/tools/motifscan.html) to analyze whether the other FadD proteins, FadD2-6, contain a leucine zipper structure. The results showed that the other FadD proteins did not contain a leucine zipper structure, and EMSA analysis showed that FadD2, which shares about 59.82% amino acid identity with FadD1, did not bind to the *lasR* gene promoter (Figure S11A, B). To further investigate the role of the leucine zipper motif of FadD1, we reconstructed FadD2 by adding the leucine zipper structure of FadD1 to the same part of FadD2 (named FadD2_LZ_) and tested whether the reconstructed FadD2 has regulatory activity similar to that of FadD1. Therefore, FadD2_LZ_ was purified to homogeneity using affinity chromatography (Figure S11C), and assayed whether it could bind to the *lasR* promoter. Intriguingly, we found that the *lasR* promoter DNA fragment formed stable DNA-protein complexes with FadD2_LZ_ and migrated more slowly than unbound probes. The amount of labeled probe that bound to FadD2_LZ_ increased with increasing amounts of FadD2_LZ_ but decreased in the presence of a 50-fold greater concentration of unlabeled probe (Figure S11D). Interestingly, addition of exogenous *cis*-DA also enhanced the binding of FadD2_LZ_ to the *lasR* promoter probe at a final concentration of 20 µM (Figure S11E). In addition, *in trans* expression of the *fadD2_LZ_* gene rescued the defective swarming motility and pyocyanin production phenotypes of the Δ*fadD1* strain (Figure 11F, G). And we also found that the addition of exogenous *cis*-DA rescued the impaired motility and pyocyanin of Δ*dspI*Δ*fadD1*(*fadD2_LZ_*) (Figure S11F, G). These results showed that the leucine zipper motif plays a critical role in the regulatory activity of FadD1.

### Analysis of the catalytic motif of FadD1

Previous studies reported that fatty acyl-CoA ligase plays a crucial role in catalyzing the formation of fatty acyl-CoA through hydrolysis of pyrophosphate^35,36^. To test whether FadD1 can function in the uptake and activation steps of fatty acid degradation, we evaluated its enzymatic activity *in vitro*. The results showed that FadD1 had high activity on lauric acid (C12:0), while it displayed moderate activity on BDSF but no activity on *cis*-DA (Figure S12A). In addition, the previous study found that mutation of Trp433, Thr436 and Arg453, the substrate binding sties, results in complete inactivation of fatty acyl-CoA synthetase in *E. coli* (37). We continued to generated three single point mutants (FadD1^W434A^, FadD1^T437A^ and FadD1^R454A^) to explore whether the mutation affects the enzyme activity of FadD1 (Figure S12B). Similarly, it was shown that the FadD1^W434A^, FadD1^T437A^ and FadD1^R454A^ exerted no enzyme activity on *cis*-DA, BDSF and lauric acid (C12:0) (Figure S12C-E). To further characterize the relationship between enzyme activity of FadD1 and perception of *cis*-DA, we chose to test the regulatory activity of the mutant FadD1^R454A^. We *in trans* expressed the *fadD1^R454A^* in the Δ*dspI*Δ*fadD1* mutant strain. Interestingly, it was found that addition of exogenous *cis*-DA rescued the impaired motility of Δ*dspI*Δ*fadD1*(*fadD1^R454A^*) (Figure S12F). Consistent with this, EMSA analysis showed that mutation of the substrate binding stie (Arg454) would not affect the perception of *cis*-DA (Figure S12G).

### Homologs of FadD1 are widespread in bacteria

To determine whether FadD1 is widely distributed, homologs of FadD1 were sought in the genome database by using the BLAST program. It was indicated that homologs of FadD1 are distributed across many different bacterial species, including in the genera *Pseudomonas*, *Azotobacter*, *Azomonas*, and *Halopseudomonas* (Table S4). Furthermore, the homologs of FadD1 in the genera *Pseudomonas, Azotobacter* and *Azomonas* contain a leucine zipper structure (Table S4). We then selected *P. fluorescens* Migula ATCC17518, which contains both a FadD1_Pf_ protein with a leucine zipper structure and a *lasR_Pf_* gene, and the FadD1_Pf_ protein was purified by affinity chromatography and tested for its interaction with the *lasR_Pf_* gene promoter. The EMSA results showed that the FadD1_Pf_ protein from *P. fluorescens* could also bind to the *lasR_Pf_* gene promoter (Figure S13), indicating that the transcriptional regulatory mechanism of FadD1 might be widely conserved in bacteria.

## DISCUSSION

Fatty acid-CoA ligase plays a vital role in fatty acid metabolism, as it can convert fatty acids to fatty acyl-CoA in bacteria^37^. In this study, we found that the fatty acid-CoA ligase FadD1 of *P. aeruginosa* not only exhibits high activity for the catalysis of lauric acid (C12:0) to lauric acid-CoA (Figure. S12A), and employs a key role in the bacterial competition with *B. cenocepacia* by degrading its QS signal BDSF (Figure 1), but also plays a critical role in the regulation of *P. aeruginosa* physiology and pathogenesis by acting as the receptor and response regulator of its QS signal *cis*-DA (Figure 2 and Table S1). We demonstrated that FadD1 can directly bind to the promoter of the target genes, thereby regulate the transcription levels of the target genes (Figure 3A and Figure S7). It was further identified that mutation of the leucine zipper structure of FadD1 abolished the binding of FadD1 to the target gene promoters (Figure 3I). Interestingly, we found that FadD2 does not contain a leucine zipper structure, and EMSA analysis showed that FadD2 did not bind to the *lasR* gene promoter (Figure S11B). However, the reconstructed FadD2 with addition of the leucine zipper structure of FadD1 to the same part of FadD2 (named FadD2_LZ_) obtained the regulatory activity similar to that of FadD1. It could bind to the *lasR* gene promoter and restore the *fadD1* mutant phenotypes (Figure S11). Furthermore, our findings indicate that FadD1 is widely conserved in many other bacterial species, including *Pseudomonas*, *Azotobacter*, and *Azomonas* species (Table S4), indicating that FadD1 with a leucine zipper structure might be a new family of transcriptional regulatory proteins widely conserved in bacteria.

The fatty acid molecule *cis*-DA was first identified in *P. aeruginosa* to act as the autoinducer of biofilm dispersion^31^. It has also been shown to induce biofilm dispersion in a range of Gram-negative and Gram-positive bacteria and in the fungal pathogen *Candida albicans*^31^. Although the previous study reported that the two-component sensor/response regulator hybrid DspS (PA4112, PA14_10770) is required and essential for native biofilm dispersion in response to the cell-to-cell signaling molecule *cis*-DA^34^, there was no exploration of other functions of *cis*-DA. In this study, we found that deletion of *dspI*, which is the synthase of *cis*-DA, caused an impairment in motility and pyocyanin production (Figure 4B, C). Interestingly, in *trans* expression of *fadD1* rescued the defective motility and pyocyanin phenotypes of the Δ*dspI* strain (Figure 4B, C). However, the addition of exogenous *cis*-DA showed no effect on these phenotypes of the mutant Δ*fadD1* and Δ*dspI*Δ*fadD1* (Figure 4B, C). Moreover, *cis*-DA is shown to bind to FadD1 with high affinity and enhanced the binding of FadD1 to the target gene promoter probes (Figure 4A and Figure 5A). We also identified that two amino acid residues at positions Gly488 (G488) and Arg552 (R552) are critical for the interaction between FadD1 and *cis*-DA (Figure 5). Mutations at Gly488 and Arg552 abolished the binding between FadD1 and *cis*-DA (Figure 5D, E). Another interesting observation in this study was that mutation at the substrate binding stie Arg454 (R454) completely abolished the enzyme activity of FadD1 on both BDSF and lauric acid (C12:0) (Figure S12C-E), but could not affect the effect of *cis*-DA on FadD1 (Figure S12G), suggesting that the *cis*-DA-binding site is different from the substrate-binding stie in FadD1. Together, FadD1 was identified to be a new response regulator of *cis*-DA QS signal.

BDSF has been identified to be a DSF-family QS signal in *B. cenocepacia*^11^. As *cis*-DA shares a similar chemical structure with that of BDSF, we then continued to explore the interaction between BDSF and FadD1. Although FadD1 showed a degradation activity on BDSF, we found that FadD1 tightly binds to BDSF with an estimated dissociation constant (*K_D_*) of 1.65 ± 0.21 μM (Figure S14A). Interestingly, the addition of exogenous BDSF also enhanced the binding of FadD1 to the *lasR* promoter probe at a final concentration of 10 µM (Figure S14B). To distinguish the role of BDSF to be a “substrate” or “ligand” of FadD1, we then used EMSA analysis and the result showed that mutation of the *cis*-DA-binding sites abolished the perception of *cis*-DA (Figure 5), but will not affect the effect of BDSF on FadD1 (Figure S14C). However, EMSA analysis showed that mutation of the substrate-binding stie (Arg454) completely abolished the effect of BDSF on the binding of FadD1 to target gene promoter (Figure S14D), while not affect the effect of *cis*-DA on FadD1 (Figure S12G). These results suggested that BDSF also interferes with the binding of FadD1 to target gene promoters *in vitro* as a substrate of the canonical enzyme FadD1.

In *P. aeruginosa,* there are at least three QS systems, namely, *las*, *rhl* and *pqs*, which are involved in the regulation of a broad range of genes important for metabolism and virulence^18,19^. These QS systems are hierarchically interrelated, and the *las* system was verified to monitor both the *rhl* and *pqs* systems^17,30^. In this study, we identified that FadD1 affects the QS-regulated phenotypes, including the biofilm formation, swarming, and pyocyanin in *P. aeruginosa* (Figure 2A-C). Surprisingly, we also identified that FadD1 is a *cis*-DA receptor protein that positively regulates the *las*, *rhl* and *pqs* QS systems in *P. aeruginosa* (Figure 2 and Figure 6). These results indicated that FadD1 is not only a response regulator of *cis*-DA but also an important upstream regulator of the QS signaling network in *P. aeruginosa*. FadD1 regulates numerous target genes and physiological functions by monitoring the QS signaling systems, which broadens our understanding of the QS signaling network of *P. aeruginosa*.

**Figure 6.**
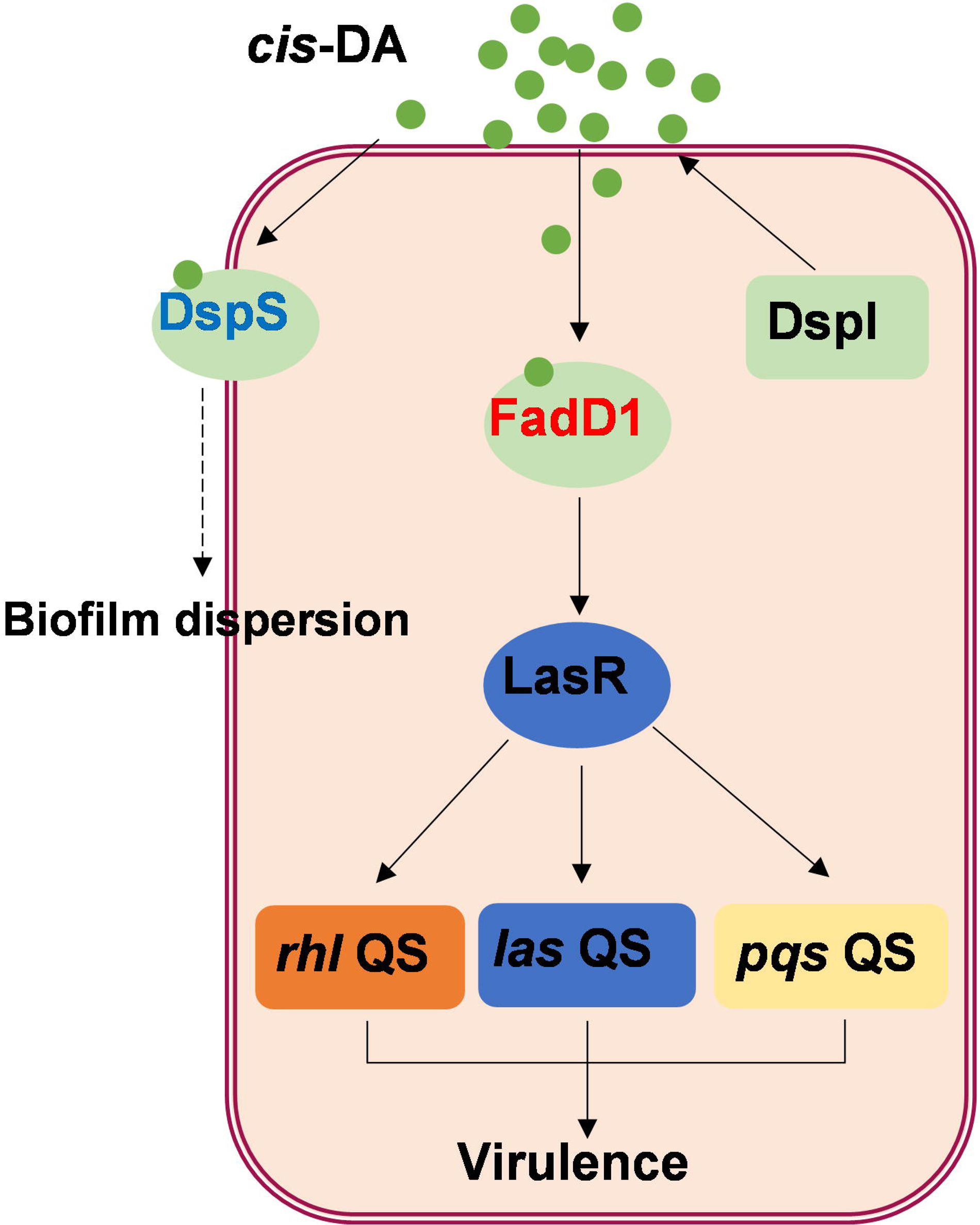
Schematic representation of *cis*-DA-regulated QS signaling in *P. aeruginosa*. DspI synthesized *cis*-DA signal molecule. FadD1, the signal receptor of *cis*-DA, perceives *cis*-DA signal and regulates the transcription of *lasR*, which positively regulates the *las*, *rhl* and *pqs* QS systems in *P. aeruginosa*. DspS is also the sensor of *cis*-DA involved in biofilm dispersion in response to *cis*-DA. The individual QS circuits are highly interconnected.

In summary, our findings suggest that FadD1, which contains a leucine zipper structure in the AMP-B domain, is not only an enzyme of fatty oxidation but also a receptor of *cis*-DA that directly controls target gene expression and controls the QS network in *P. aeruginosa*. In addition, the BLAST search revealed that homologs of FadD1 containing the leucine zipper structure are highly conserved in many bacteria (Table S4), suggesting that the regulatory mechanism of FadD1 might be present in various bacterial species. Consistently, our recent study revealed that the fatty acyl-CoA ligase DsfR (BCAM2136), which contains a leucine zipper motif, acts as a global transcriptional regulator to control *B*. *cenocepacia* virulence by sensing BDSF^47^. Different from FadD1, DsfR has a specific BDSF-sensing site and showed no enzyme activity on BDSF, while FadD1 only binds with BDSF on the substrate-binding sites. These results indicate that FadD1 and DsfR share some common characteristics and are distinguished from each other. In conclusion, our work presents a unique and widely conserved signal receptor of *cis*-DA, which might be an important new type of QS signal receptor in bacteria.

### Limitations of the study

* Limitations of the Study: As it reads right now, the section mostly focuses on future directions of the work to answer outstanding open questions. While future directions can belong to this section, it is mostly the place to openly discuss potential caveats of the work (technical, conceptual, or other limitations) with the intent of promoting clarity and transparency. I would like to suggest that you further develop and edit this section in this direction.

Our findings demonstrated that FadD1 is a specific receptor protein of *cis*-DA QS signal. It was revealed that *cis*-DA binds to FadD1 to enhance the binding of FadD1 to the promoter DNA of target genes. However, it remains unknown how does the binding to *cis*-DA induce the regulatory activity of FadD1 on the transcription of target genes.

## ACKNOWLEDGMENTS

This work was financially supported by the National Key Research and Development Program of China (2021YFA0717003 to Y.D.), the National Natural Science Foundation of China (32300033 to S.S.) and the Scientific Research Foundation of Hainan University (KYQD(ZR)-23006 to S.S.).

## AUTHOR CONTRIBUTIONS

Y.D. and S.S. conceived the project. S.S., J.L., and Y.D. designed the research. S.S., J.L., B.W., Y.S., H.H., X.S., M.W., and B.C. performed the research. S.S., J.L., G.W., Y.H., L.X., B.G., L.Y., X.W., L-H.Z., and Y.D. analyzed the data. S.S. and Y.D. wrote the paper.

## INCLUSION AND DIVERSITY

We support inclusive, diverse, and equitable conduct of research.

## STAR METHODS

### KEY RESOURCES TABLES

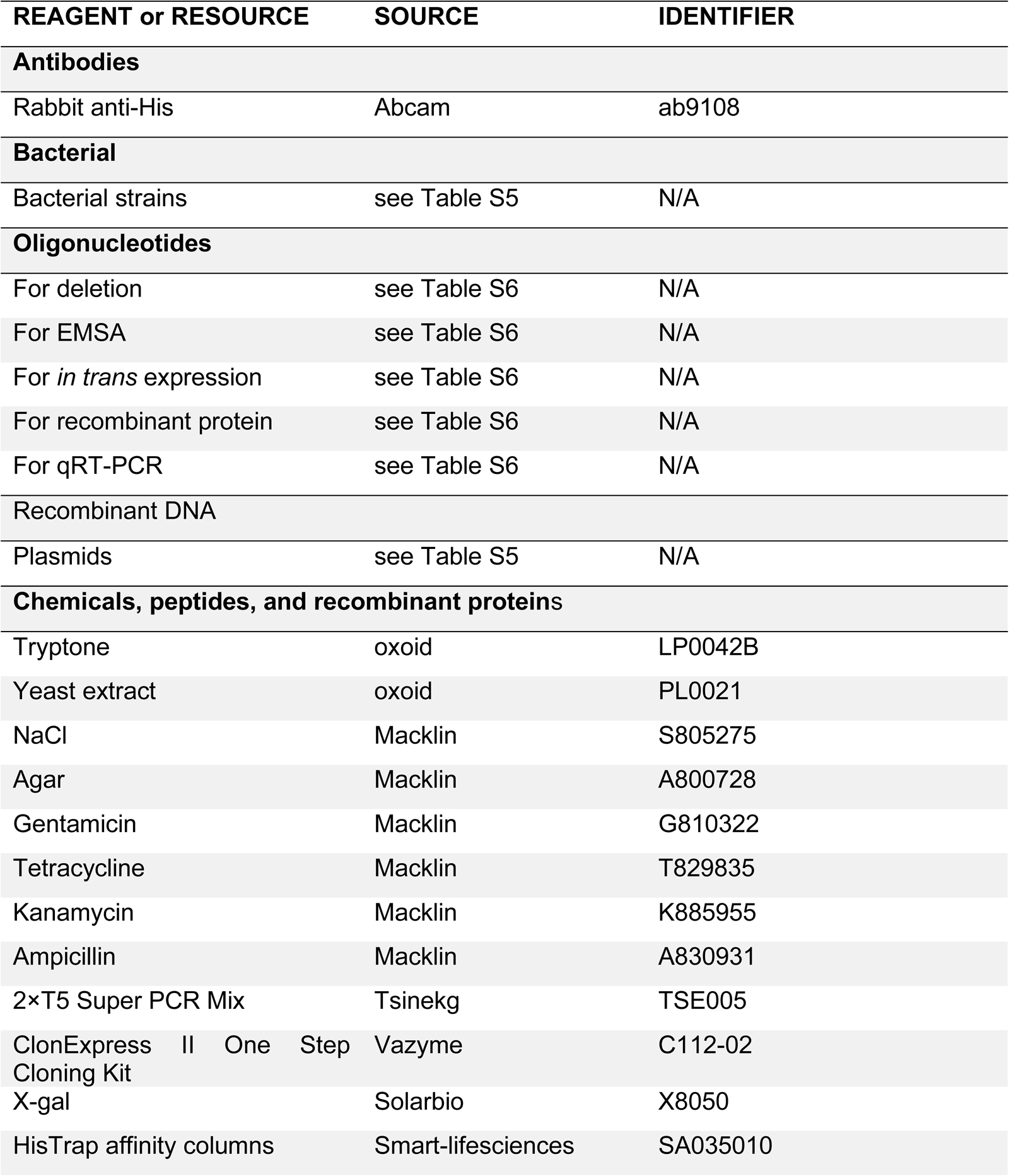
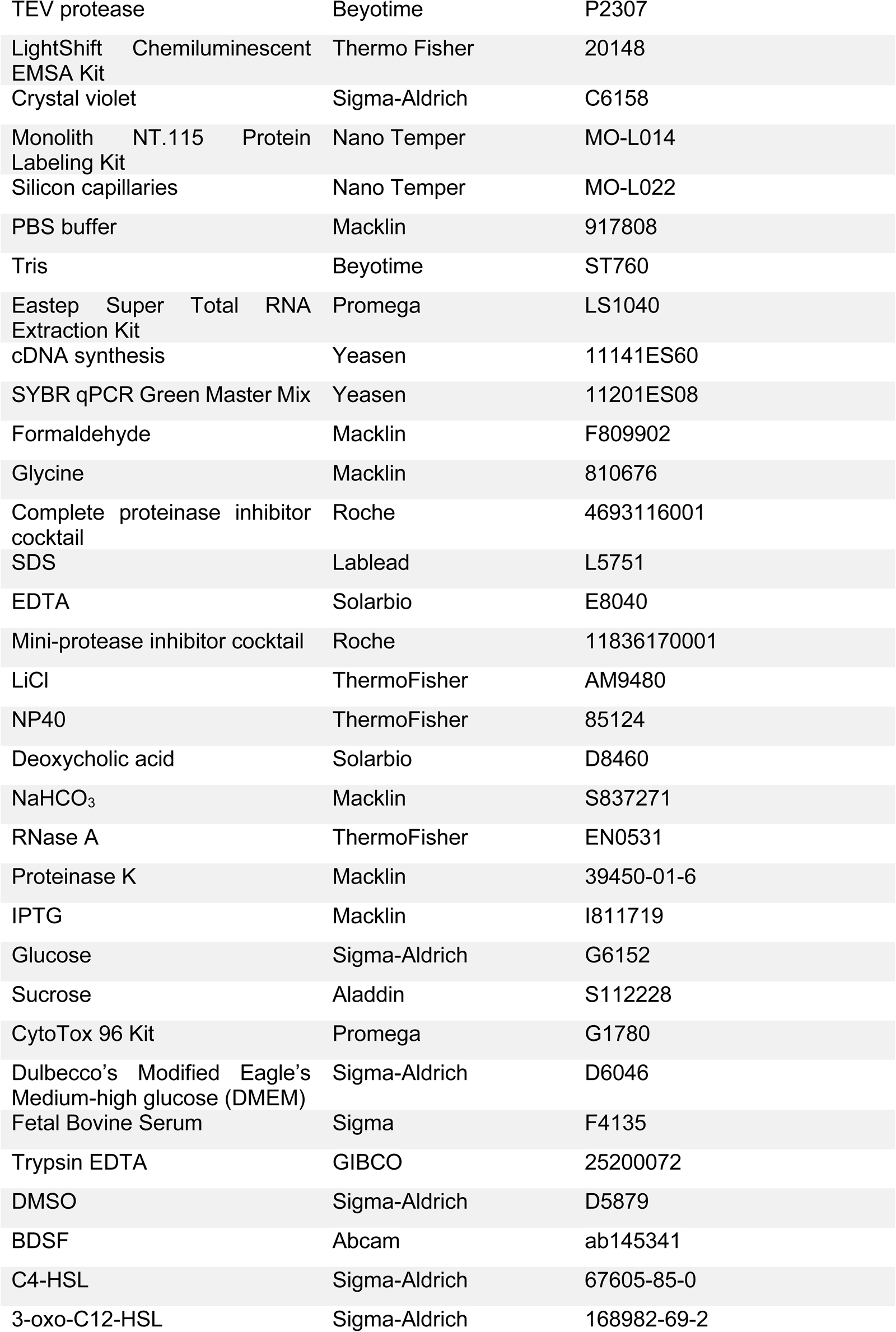
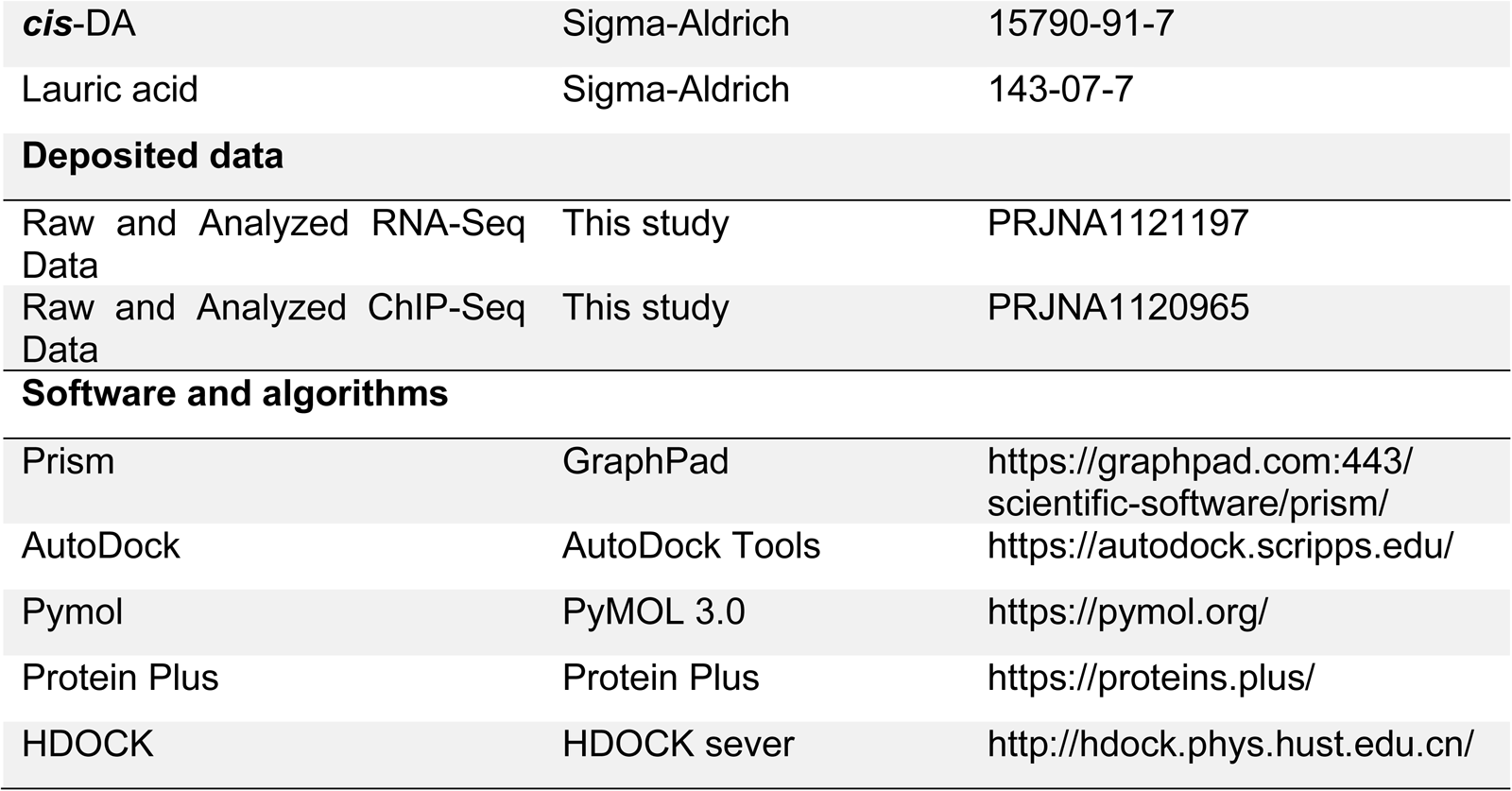

### RESOURCE AVAILABILITY

#### Lead contact

Further information and requests for resources and reagents should be directed to and will be fulfilled by the lead contact Yinyue Deng.

#### Materials availability

Constructs and reagents in this study will be made available upon request, but a completed Materials Transfer Agreement may be required if there is potential for commercial application.

#### Data and code availability

Unprocessed data related to figures in this manuscript are available from the lead contact upon request.

This paper does not report original code.

Any additional information required to reanalyze the data reported in this work paper is available from the lead contact upon request.

### EXPERIMENTAL MODEL AND STUDY PARTICIPANT DETAILS

#### Bacterial strains and culture conditions

The bacterial strains and plasmids used in this work are listed in Table S5. *P. aeruginosa* PAO1, *P. fluorescens* Migula ATCC17518 and *B. cenocepacia* H111 strains obtained from the American Type Culture Collection (ATCC) were maintained in Luria-Bertani broth (LB; 10 g/L tryptone, 5 g/L yeast extract, 5 g/L NaCl) at 37°C. The following antibiotics were added when necessary: ampicillin and kanamycin at 100 µg mL^-1^; tetracycline, 10 µg mL^-1^; and gentamicin, 10 µg mL^-1^. BDSF and *cis*-DA (HPLC ≥ 99%) were dissolved in methanol at a final concentration of 100 mM, and this solution was added to the medium in the experiments. Bacterial growth was determined by measuring the optical density at 600 nm.

#### Cell culture

A549 cell were maintained in Dulbecco’s modified Eagle’s medium (DMEM) supplemented with 10% fetal bovine serum (FBS) and 1% penicillin‒streptomycin, cells were cultivated in a humidified atmosphere containing 5% CO_2_ at 37 °C.

### METHOD DETAILS

#### BSDF extraction

The extraction of BDSF from culture supernatants was conducted following previously described methods^11,53^. To quantify BDSF production in the culture of *B. cenocepacia* strain or *P. aeruginosa* strain, 100 mL of the supernatant was collected. Its crude ethyl acetate extract was passed through a 0.22 µm Minisart filter unit and was then condensed to 1 mL for LC-MS analysis.

#### Quantification of BDSF and *cis*-DA

Quantification of BDSF and *cis*-DA was performed by using an LC-MS system, which consisted of an ACQUITY UPLC system and a Waters Q-Tof Premier high-resolution mass spectrometer^53^. An ACQUITY UPLC BEH C18 column 1.7 µm (2.1 × 50 mm) was used for the chromatography analysis of BDSF and *cis*-DA, which was eluted with a CH_3_OH gradient in water at 65-100% supplemented with 0.01% formic acid at a flow rate of 0.4 mL·min^-1^ for 10 min. Next, 65% CH_3_OH was used for 3 min. The entire elution column was introduced into the Q-TOF mass spectrometer according to the manufacturer’s instructions. The BDSF and *cis*-DA levels in the culture supernatant were measured using the peak area in the extracted ion chromatogram.

#### Protein expression and purification

The coding regions of proteins were amplified with the primers listed in Table S6, and were attached and fused to the expression vectors pET-21a, pET-28a and PDBHT2. The fusion gene constructs were transformed into *E. coli* strain BL21. Affinity purification of the 6×His fusion proteins was performed using Ni-NTA resin (Smart-lifesciences). The His-tag was removed using TEV protease (Beyotime), and the cleaved fusion protein was analyzed by SDS-PAGE^54,55^.

#### Enzyme activity assays

The activity of FadD1 was determined following the methods^56^. *cis*-DA, BDSF or lauric acid (C12:0) were dissolved in TME buffer (Tris-HCl, 60 mM; MgCl_2_, 10 mM; EDTA, 1 mM; coenzyme A (CoASH), 0.5 mM; and Triton X-100, 0.1%; pH 7.5) and added to the mixture at a final concentration of 200 µM, while the final concentrations of FadD1 and its derivatives were 20 µM. The reaction mixture was incubated at 37°C, and the reaction was stopped by placing sample tubes in boiling water for 5, 15, 30 and 60 min. *cis*-DA, BDSF or lauric acid (C12:0) levels were measured by LC-MS spectrometry.

#### Construction of in-frame deletion mutants and complementation

*P. aeruginosa* PAO1 was used as the parental strain for the generation of an in-frame deletion mutant of *fadD1* and *dspI* by the following method described previously^14^. The primers used to generate upstream and downstream regions flanking *fadD1* and *dspI* are listed in Table S7. For complementation analysis, the coding regions of *fadD1* and *dspI* were amplified by PCR and cloned into the plasmid pBBR1-MCS5, pBBR1-MCS2 and pLAFR3 using the primers listed in Table S6. The resulting constructs were conjugated into the *P. aeruginosa* PAO1 mutant, *B. cenocepacia* H111 strain by electroporation^14,54^.

#### Bacterial growth analysis

An overnight bacterial culture in LB medium was inoculated into fresh media to an OD600 of 0.01. A 10 mL cell suspension was grown in 50 mL conical centrifuge tubes at 37°C with shaking at 220 rpm. Bacterial growth was determined by measuring the optical density at 600 nm every 4 hours. LB medium was used as the negative control^47^.

#### Competition assays in the mixed culture

The green fluorescent protein expression vector was used and transformed into the *P. aeruginosa* PAO1 WT, Δ*fadD1* and Δ*fadD1*(*fadD1*) strains. The mCherry fluorescent protein expression vector was introduced into the *B. cenocepacia* H111 strain by electroporation. Then, the *P. aeruginosa* PAO1 WT, Δ*fadD1* and Δ*fadD1*(*fadD1*) strains were cocultured with the *B. cenocepacia* H111 strain at an initial ratio of 1:4 (vol/vol) at OD_600_ = 0.1 at 37°C with shaking at 220 rpm for 12 h. Then, the quantification of BDSF was performed by using an LC-MS system, and the mixed cultures were analyzed by a Spectra Max i3x multifunctional enzyme labeling instrument (Molecular Devices, CA, USA)^57^.

#### *P. aeruginosa* phenotype assays

For the analysis of biofilm formation^58^, bacterial cells were grown overnight at 37°C and inoculated into LB media (150 μL per well) in 96-well polypropylene microtiter plates. Microtiter plates with fitted lids were incubated at 37°C with shaking at 200 rpm for 12 h. The plates were then washed to remove planktonic cells and stained for 20 min with 1% (weight/vol) crystal violet (Sigma-Aldrich). After washing with water, 200 µL of 95% (vol/vol) ethanol was added to the wells to release the stain. Biofilm was determined by measuring the absorbance of the resulting solution at 570 nm.

Motility was determined on 0.3% semisolid agar (Becton, Dickinson and Company). Bacteria were inoculated into the center of plates containing 0.8% tryptone, 0.5% glucose, and 0.3% agar. The plates were incubated at 30°C for 18 h before the diameter of the colony was measured^54^.

The pyocyanin assay was performed with the method described previously^49^. Briefly, bacteria were cultured at 37°C for approximately 12 hours and centrifuged at 13,000 rpm for 1 min, and 1.5 mL supernatants were collected and extracted with double volume chloroform with vigorous shaking at room temperature for 30 min. The solvent phase was transferred to a new tube containing 1 mL of 1 N HCl (Sigma-Aldrich). The mixture was shaken gently to transfer pyocyanin to the aqueous phase. The quantity of pyocyanin was determined by measurement of absorbance at 520 nm and normalization against the cell density.

#### *B. cenocepacia* phenotype assays

Biofilm formation in 96-well polypropylene microtiter dishes was assayed essentially as described previously by *P. aeruginosa*. Swarming motility was determined on semisolid agar (0.3%). Bacteria were inoculated into the center of plates containing 0.8% tryptone, 0.5% glucose, and 0.3% agar. The plates were incubated at 30°C for 18 h before the diameter of the colony was measured.

Protease assays were performed following a previously described method^14^. Briefly, bacteria were cultured at 37°C for approximately 12 hours. Cultures were centrifuged at 13,000 rpm for 5 min, and the supernatants were removed and filtered through a 0.22 μm membrane. One hundred microliters of supernatant was incubated at 30°C with an equal volume of azocasein (Macklin) dissolved in proteolytic buffer (5 mg/mL) for 30 min. The reaction was stopped by the addition of 400 μL of 10% (weight/vol) TCA (Macklin) buffer. After incubation for 2 min at room temperature, the mixture was centrifuged at 13,000 rpm for 1 min to remove the remaining azocasein. Supernatants were removed and mixed with 700 μL of 525 mM NaOH (Macklin). The absorbance of the azopeptide supernatant was measured at a wavelength of 442 nm. Protease activity was obtained after normalization of the absorbance against the corresponding cell density.

#### Cytotoxicity assays

Cytotoxicity was assessed by measuring the release of LDH from human A549 cells^54^. The A549 cell line was cultured in Dulbecco’s modified Eagle’s medium (Sigma-Aldrich) at 37°C with 5% CO_2_. A549 cells were seeded in 96-well plates at a density of 5 × 10^4^ cells per well and grown until they reached ∼90% confluence. Culture supernatants were removed, and the cell monolayer was washed once with phosphate-buffered saline (PBS). Fresh bacterial cells cultured to an OD_600_ of 1.0 were washed and diluted in DMEM. The diluted bacterial cells were then added to the A549 cell monolayers at a multiplicity of infection (MOI) of approximately 1000. After incubation for 8 h, the cytotoxicity was determined by measuring the released lactate dehydrogenase (LDH) in the supernatants using a cytotoxicity detection kit (Roche).

#### Construction of reporter strains and measurement of β-galactosidase activity

The promoters of *lasR, lasI, rhlR*, *rhlI, mvfR* and *pqsA* were amplified using the primer pairs listed in Table S6. The resulting products were inserted upstream of the promoter less *lacZ* gene in the vector pME2-*lacZ*. Transconjugants were then selected on LB agar plates supplemented with tetracycline and X-gal (Solarbio). Measurement of β-galactosidase activities was performed following previously described methods^14,54,55^.

#### Quantitative reverse transcription PCR assays

Bacterial cells were cultured at 37°C in LB broth to the logarithmic-growth phase (OD_600_=1.0). Cells were collected by centrifugation, and total RNA was extracted using an Eastep Super Total RNA Extraction Kit (Promega). cDNA synthesis was performed using a HiScript III 1st Strand cDNA Synthesis Kit (Yeasen). Primers for RT-qPCR are listed in Table S6. The RT-qPCR experiments were performed on a QuantStudio™ 7 RT-qPCR System (Thermo Fisher) using SYBR qPCR Green Master Mix (Yeasen). The expression of the target genes was normalized to the expression level of 16S RNA. The relative transcript abundance was calculated by the 2^−ΔΔCt^ method^55^. At least three biological replicates were performed per sample.

#### Quantitative analysis of QS signal production in *P. aeruginosa*

Bacterial cells were grown in LB medium overnight with agitation at 37°C. One liter of culture supernatant was collected by centrifugation and extracted with an equal volume of ethyl acetate. The crude extract (organic phase) was dried using a rotary evaporator and dissolved in methanol. All of the above samples were kept at 4°C until analysis. Ultrahigh-performance liquid chromatography-electrospray ionization tandem mass spectrometry (UHPLC-ESI-MS/MS) was performed in a Shimadzu LC-30A UHPLC system with a Waters C18 column (1.8 µm, 150 × 2.1 mm) and a Shimadzu 8060 QQQ-MS mass spectrometer with an ESI source interface. The mass spectrometer was operated in positive-ion mode. The mobile phase was prepared as 0.1% formic acid/water and acetonitrile^57^.

#### RNA-Seq analysis

RNA was isolated from *P. aeruginosa* strains grown to OD_600_=1.0 by using the Eastep Super Total RNA Extraction Kit (Promega, Madison, USA). The concentration was measured using a Qubit® RNA Assay Kit in a Qubit® 2.0 Fluorometer (Life Technologies, CA, USA). The purity and integrity were checked using the NanoPhotometer® spectrophotometer (IMPLEN, CA, USA) and the RNA Nano 6000 Assay Kit of the Bioanalyzer 2100 system (Agilent Technologies, CA, USA), respectively. Sequencing libraries were generated using the NEBNext® Ultra™ Directional RNA Library Prep Kit for Illumina® (NEB, USA) and then purified by the AMPure XP system. Finally, library quality was assessed by using the Agilent Bioanalyzer 2100 system. The clustering of the index-coded samples was performed on a cBot Cluster Generation System using TruSeq PE Cluster Kit v3-cBot-HS (Illumina). After cluster generation, the library preparations were sequenced on an Illumina HiSeq platform, and paired-end reads were generated. Trimmed sequence reads were aligned to the *P. aeruginosa* PAO1 genome sequence using Bowtie2-2.2.3, and normalized read counts were compared using HTSeq v0.6.1 as described previously^59,60^. Then, the FPKM of each gene was calculated based on the 26 lengths of this gene, and the read count was mapped to the gene.

#### Electrophoretic gel mobility shift assay

Electrophoretic mobility shift assay (EMSA) experiments were performed as described previously^53,54,61^. The DNA probes used for EMSA were prepared by PCR amplification using the primer pairs listed in Table S6. The purified PCR products were 3-end-labeled with biotin following the manufacturer’s instructions (Thermo Fisher). The DNA-protein binding reactions were performed according to the manufacturer’s instructions (Thermo Fisher). A 5% polyacrylamide gel was used to separate the DNA-protein complexes. After UV cross-linking, the biotin-labeled probes were detected in the membrane using a Biotin luminescent detection kit (Thermo Fisher).

#### Chromatin immunoprecipitation sequencing analyses

Chromatin immunoprecipitation sequencing (ChIP-Seq) was performed according to previously published methods with some modifications^62^. ChIP-Seq analysis was conducted by the Wuhan IGENEBOOK Biotechnology Co. Ltd. (Wuhan, China). The anti-HIS antibody (ab9108, 1:1000, Abcam) was used for immunoprecipitation, and DNA fragments enriched by ChIP were sequenced on an Illumina HiSeq 2000. After removing sequencing adaptors and low-quality bases, the clean reads were mapped to the *P. aeruginosa* PAO1 genome using BWA software (version 0.7.15-r1140). The peak caller MACS (version, 2.1.1.20160309; *q* value, 0.05) was used to localize the potential binding sites of FadD1^63^.

#### Molecular docking FadD1

The 3D structure of the double-stranded DNA was built in the MOE1 DNA/RNA Builder. Protein-DNA docking in the HDOCK2 server was used for molecular docking simulation of FadD1 and DNA^64,65^. The 3D structure of FadD1 (Q9HYU4) predicted by AlphaFold2 was set as the receptor, and the 3D structure of the DNA was set as the ligand^66^. The HDOCK server automatically predicts their interaction through a hybrid algorithm of template-based and template-free docking. First, the server performs template-based modeling to identify possible homologous DNA templates. Second, the intrinsic scoring function for protein-RNA interactions is integrated in template-free docking. Finally, the top 100 predicted complex structures are provided to users for download, of which the top 10 models will be visualized on the result web page. The intermolecular contacts from the most likely poses were further evaluated. The models of the complex and interface residues were analyzed using MOE. Molecular graphics were generated by PyMOL.

AutoDock and PyMOL were used for docking and binding site analysis. The crystal structure of FadD1 from the AlphaFold2 was used for docking with *cis*-DA by AutoDock software. To screen the potential binding sites, a grid box was prepared to wrap the entire macromolecule. The receptor grid was centered at center_x = -1.038, center_y = 1.49, center_z = 11.31, size_x = 126, size_y = 126, and size_z = 126. The exhaustiveness was set at 50. The top scoring ligand conformation was chosen, and the potential binding sites were visualized in 3D by PyMOL. The potential binding sites were visualized in 2D by Protein Plus (https://proteins.plus/).

#### Microscale thermophoresis assay

Protein-binding experiments were carried out with a Nano Temper 16 Monolith NT.115 instrument (NanoTemper Technologies; www.nanotemper-technologies.com)^55^. In brief, the protein was labeled with the L014 Monolith NT.115 Protein Labelling Kit (Nano Temper). The labeled protein and different titers of unlabeled *cis*-DA were mixed and loaded onto standard treated silicon capillaries (Nano Temper), and fluorescence was measured. The measurements were carried out at 20% LED power and 40% MST power.

### STATISTICAL ANALYSIS

No statistical method was used to predetermine the sample size. No data were excluded from the analyses. The data are presented as the means ± standard deviations. Statistical analyses were performed with Prism 8 software (GraphPad). Statistical significance is indicated as follows: \**p* < 0.05; \*\**p* < 0.01; \*\*\**p* < 0.001; ns = no significance (one-way ANOVA or two-way ANOVA). Biological replicates and numbers of independent experiments were stated in the legends. All experiments presented as representative LC-MS graphs or gels were repeated at least 3 times with similar results.

