## Supporting Information for "The long-chain fatty acid-CoA ligase FadD1 monitors *Pseudomonas aeruginosa* quorum sensing as a receptor of cis-2-decenoic acid"

### Supplementary Figures and Tables

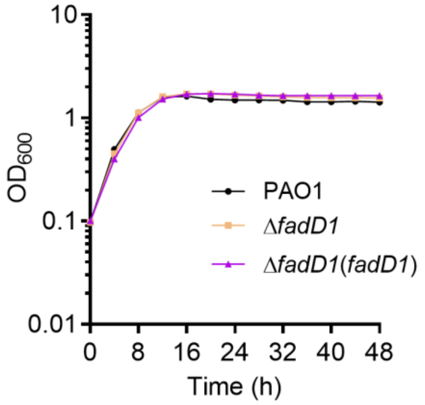


**Figure S1. Effect of *fadD1* on the growth of the *P. aeruginosa* PAO1 strains in LB medium.**

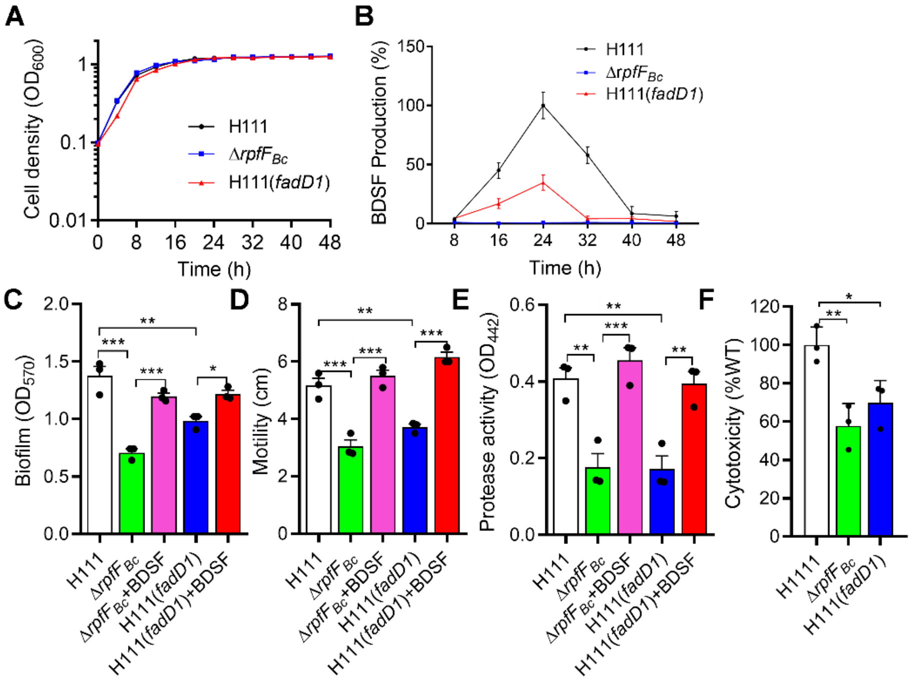


**Figure S2. Effects of *fadD1* on the BDSF production and BDSF-regulated phenotypes of *B. cenocepacia* H111 strain.** (A) Effects of *fadD1* on the growth curve of *B. cenocepacia* H111 ($n$ = 3 biological replicates). (B) The production of BDSF in the *B. cenocepacia* H111 strains ($n$ = 3 biological replicates). (C) Biofilm formation, (D) motility, (E) protease activity and (F) cytotoxicity in the *B. cenocepacia* H111 strains were measured ($n$ = 3 biological replicates). The data are the means ± standard deviations of three independent experiments. The statistical comparisons were performed using one-way ANOVA (*$p < 0.05$; **$p < 0.01$; ***$p < 0.001$).

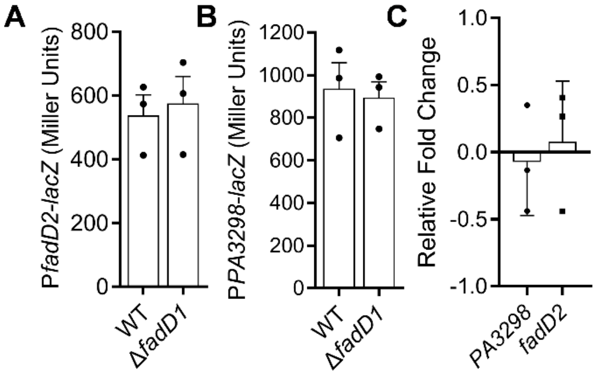


**Figure S3. Influence of mutation of *fadD1* on the expression levels of upstream and downstream genes of *fadD1*.** Effects of *fadD1* on the gene expression levels of *fadD2* (A) and *PA3298* (B) by assessing the β-galactosidase activity of the promoter-*lacZ* transcriptional fusions and by RT-qPCR (C) (*n* = 3 biological replicates). The data are the means ± standard deviations of three independent experiments.

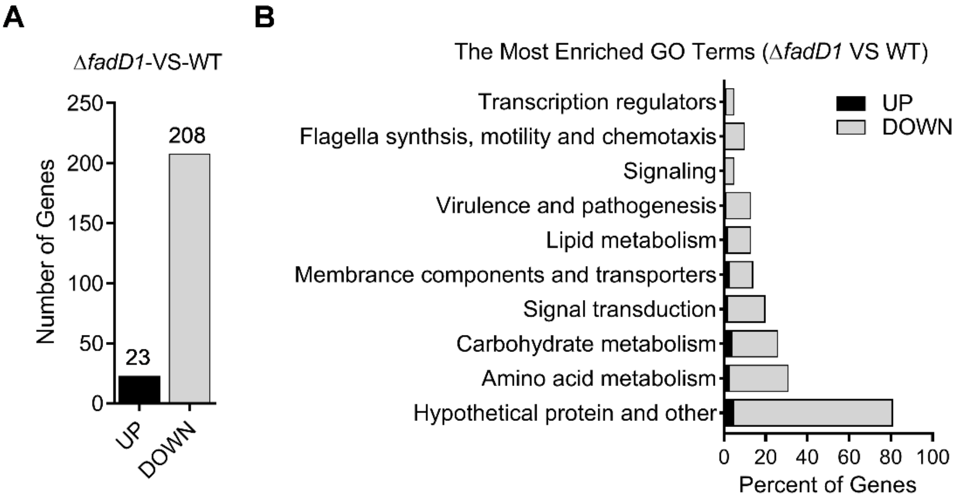


**Figure S4. Differential gene expression profiles between the Δ*fadD1* mutant strain and the wild-type strain as measured by RNA-Seq (|$Log_2$-fold fold change| ≥1.0).** (A) A number of genes were upregulated and downregulated in Δ*fadD1* compared with the wild-type strain. (B) KEGG classification analysis of differentially expressed genes between the Δ*fadD1* and wild-type strains.

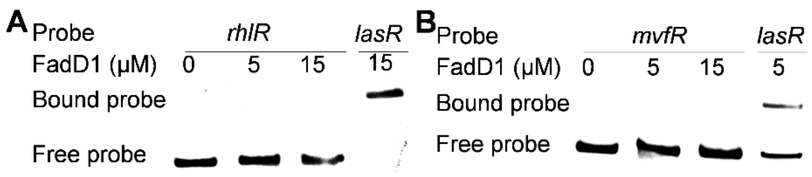


**Figure S5. EMSA analysis of the binding between FadD1 protein and the promoters of the QS receptor genes in *P. aeruginosa*.** EMSA analysis of *in vitro* binding of FadD1 to the promoters of *rhlR* (A) and *mvfR* (B), in which the biotin-labeled 270-bp *rhlR* promoter and 306-bp *mvfR* promoter DNA probes were used for the protein binding assay. The EMSA experiments were performed three times, and representative images from one experiment are shown.

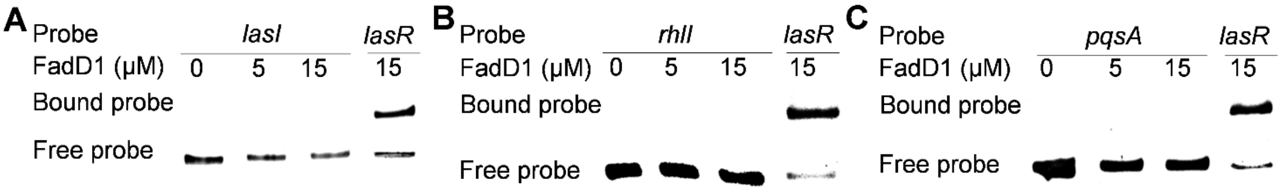


**Figure S6. EMSA analysis of the binding between FadD1 protein and the promoters of the QS synthase-encoding genes in *P. aeruginosa*.** EMSA analysis of the *in vitro* binding of FadD1 to the promoters of *lasI* (A), *rhlI* (B) and *pqsA* (C), in which biotin-labeled 321-bp *lasI* promoter, 330-bp *rhlI* promoter, and 324-bp *pqsA* promoter DNA probes were used for the protein binding assay. The EMSA experiments were performed three times, and representative images from one experiment are shown.

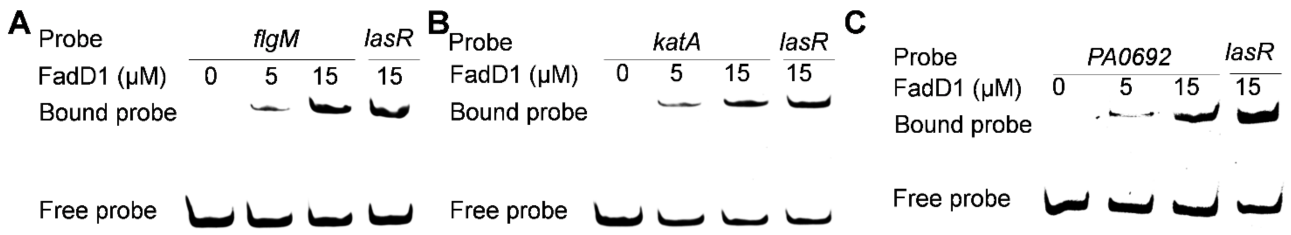


**Figure S7. EMSA analysis of the binding between FadD1 protein and the promoters of the target genes, *flgM*, *katA* and *PA0692*.** EMSA analysis of the *in vitro* binding of FadD1 to the promoters of *flgM* (A), *katA* (B) and *PA0692* (C), in which biotin-labeled 420-bp *flgM* promoter, 330-bp *katA* promoter and 285-bp *PA0692* promoter DNA probes were used for the protein binding assay. The EMSA experiments were performed three times, and representative images from one experiment are shown.

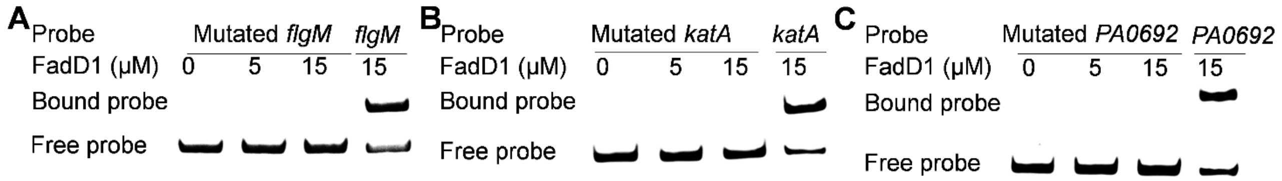


**Figure S8. EMSA analysis of the FadD1 binding site in the promoters of the target genes, *flgM*, *katA* and *PA0692*.** Analysis of the binding between FadD1 and the mutated *flgM* (A), *katA* (B) and *PA0692* (C) promoters with deletion of the FadD1 binding sequence 5’-TGGCCGG-3’, 5’-AGGAGGG-3’, and 5’-AGGGCGG-3’. EMSA analysis was performed *in vitro*. The EMSA experiments were performed three times, and representative images from one experiment are shown.

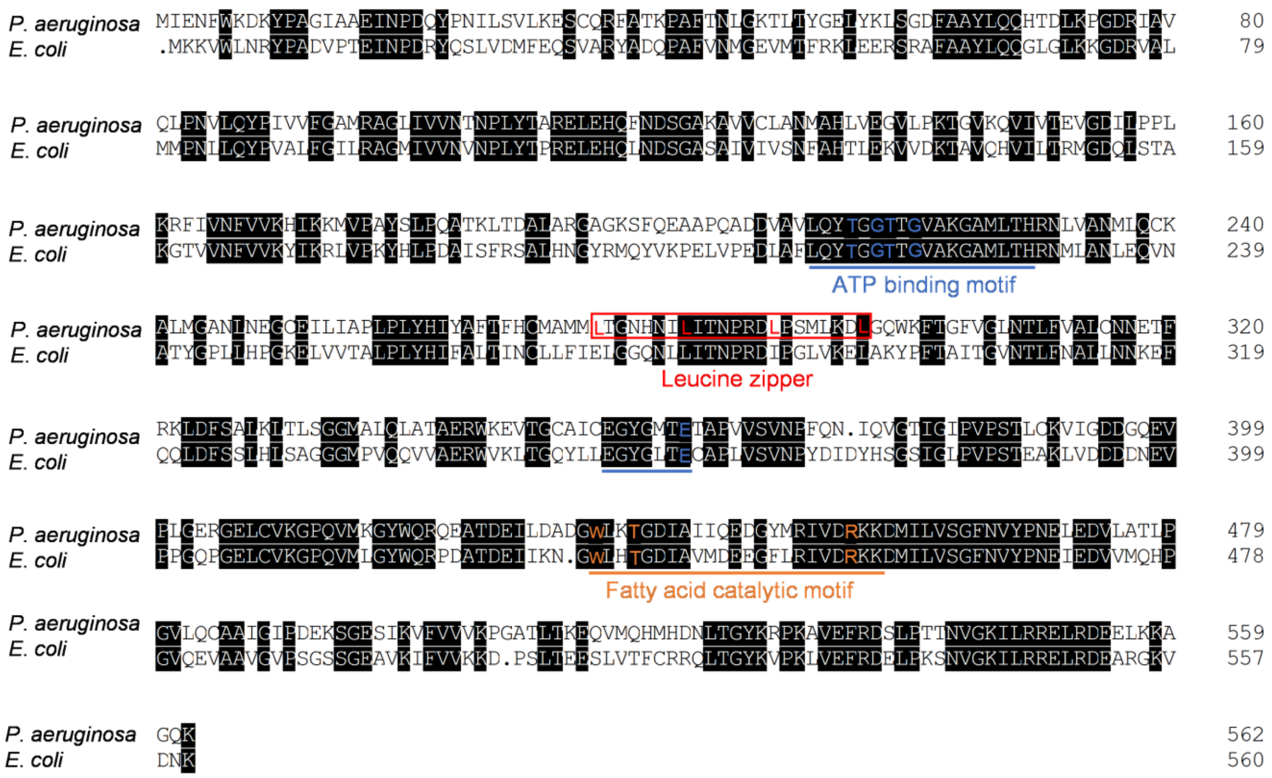


**Figure S9. Multiple sequence alignment of FadD1 of *P. aeruginosa* PAO1 and FadD of *E. coli* was performed and shaded using the DNAMAN 8.** The key amino acid for the leucine zipper motif is colored red. The key amino acid for fatty acid catalysis motif is colored orange.

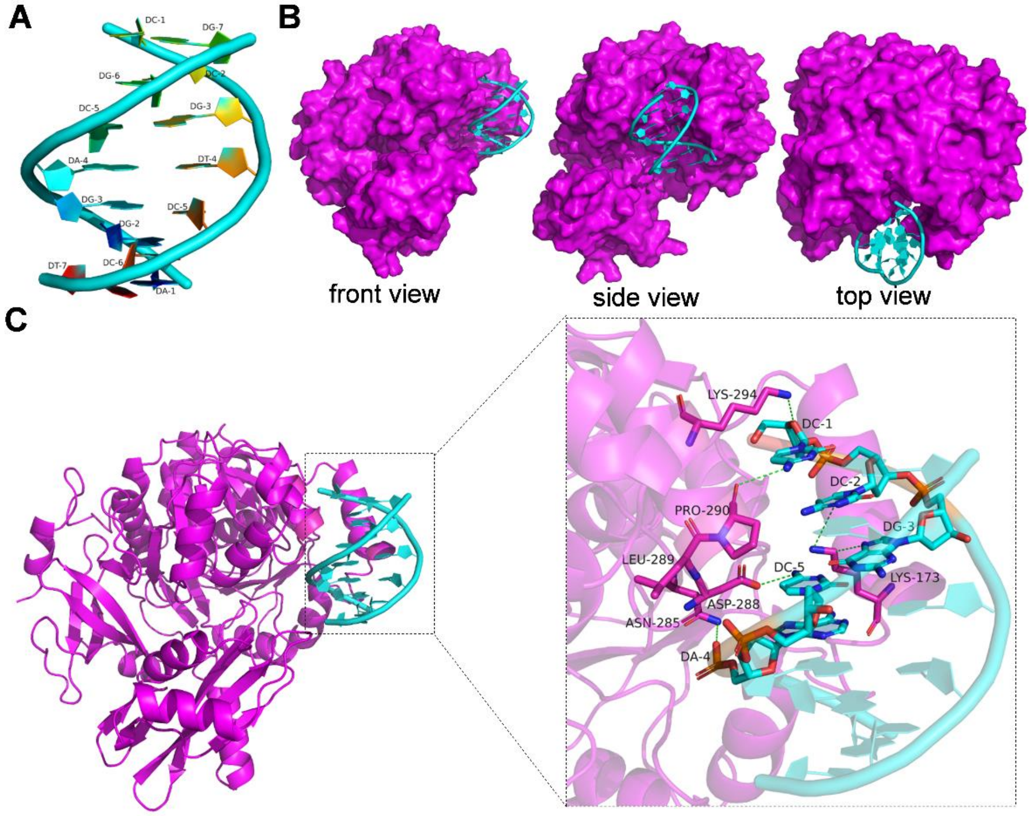


**Figure S10. Molecular docking predicts the binding model of the FadD1 protein and the binding sequence 5'-AGGACGG-3' DNA.** (A) The predicted 3D structure of DNA. The surface (B) and 3D (C) binding model of DNA and FadD1. FadD1 is colored magenta, while DNA is colored cyan. The residues in FadD1 are depicted as magenta sticks, while bases in DNA are depicted as cyan sticks. The hydrogen bond and Pi-cation interactions are depicted as dashed lines.

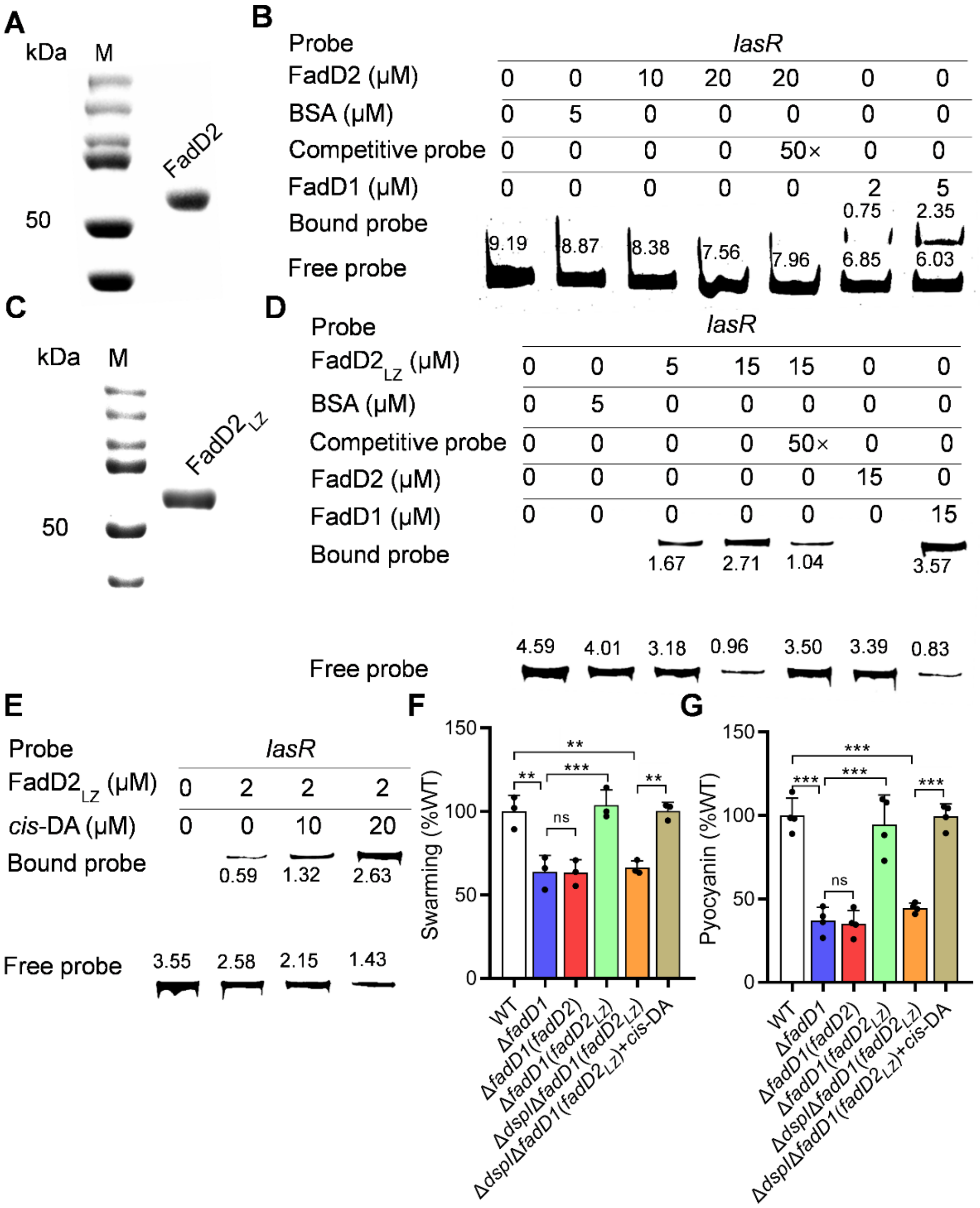


**Figure S11. Analysis of the regulatory activity of $FadD2_{LZ}$ (FadD2 containing the leucine zipper structure of FadD1) in *P. aeruginosa* PAO1.** (A) SDS-PAGE of the purified FadD2 protein. (B) EMSA analysis of *in vitro* binding of FadD2 to the promoters of *lasR*, in which the biotin-labeled 339-bp *lasR* promoter DNA probe was used for the protein binding assay. (C) SDS-PAGE of the purified $FadD2_{LZ}$ protein containing a leucine zipper structure. (D) EMSA analysis of *in vitro* binding of the $FadD2_{LZ}$ protein to the promoter of *lasR*. (E) EMSA analysis of the *in vitro* binding of $FadD2_{LZ}$ to the promoter of *lasR* with the addition of different amounts of *cis*-DA. The EMSA experiment was performed three times, and representative images from one experiment are shown. Effects of *$fadD2_{LZ}$* on the QS-regulated phenotypes: swarming motility (F) and pyocyanin production (G) in the *P. aeruginosa* (*n* = 3 biological replicates). The data are the means ± standard deviations of three independent experiments. The statistical comparisons were performed using one-way ANOVA (**$p$ < 0.01; ***$p$ < 0.001; ns = no significance).

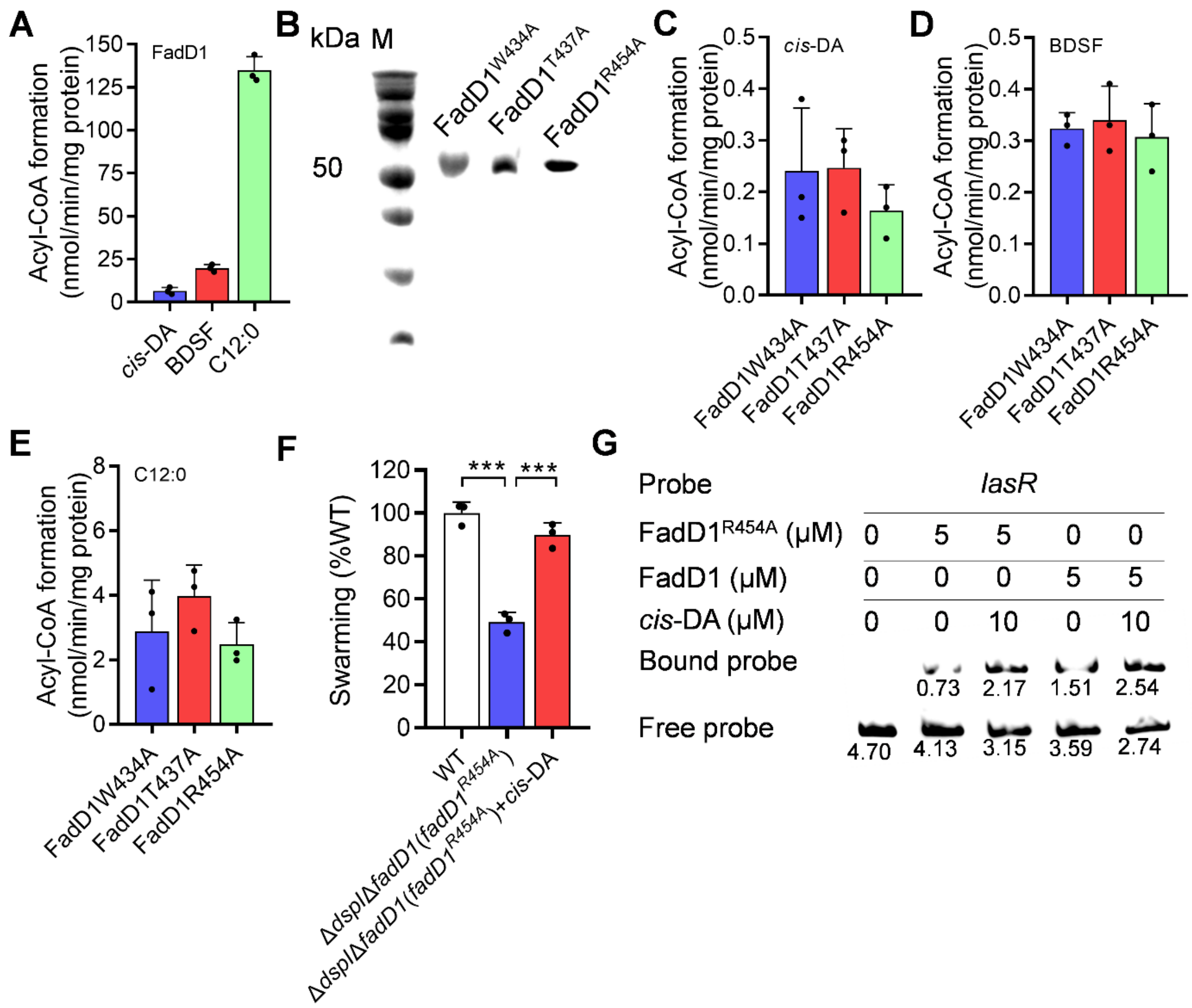


**Figure S12. Analysis of the catalytic sites of FadD1 as a fatty acyl-CoA synthetase.** (A) Analysis of fatty acyl-CoA synthetase activities of FadD1. Activities were defined using *cis*-DA (C10:1), BDSF (C12:1) and lauric acid (C12:0) as substrates ($n$ = 3 biological replicates). (B) SDS–PAGE gel electrophoresis of purified FadD1 variants. Analysis of fatty acyl-CoA synthetase activities of FadD1 variants. Activities were defined using *cis*-DA (C), BDSF (D) and lauric acid (E) as substrates ($n$ = 3 biological replicates). (F) Effects of *fadD1*$^{R454A}$ on QS-regulated swarming motility ($n$ = 3 biological replicates). (G) EMSA analysis of the *in vitro* binding of FadD1$^{R454A}$ to the promoter of *lasR* with the addition of different amounts of *cis*-DA. The EMSA experiment was performed three times, and representative images from one experiment are shown. The data are the means ± standard deviations of three independent experiments. The statistical comparisons were performed using one-way ANOVA (***$p$ < 0.001).

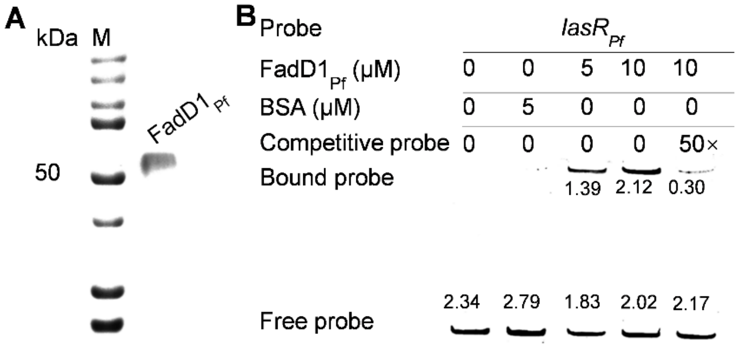


**Figure S13. EMSA analysis of the binding between the promoter of *lasR*$_{Pf}$ and FadD1$_{Pf}$ protein from *P. fluorescens* Migula ATCC17518.** (A) SDS-PAGE of the purified FadD1$_{Pf}$ protein. EMSA analysis of *in vitro* binding of FadD1$_{Pf}$ protein (B) to the promoter of *lasR*$_{Pf}$. The EMSA experiment was performed three times, and representative images from one experiment are shown.

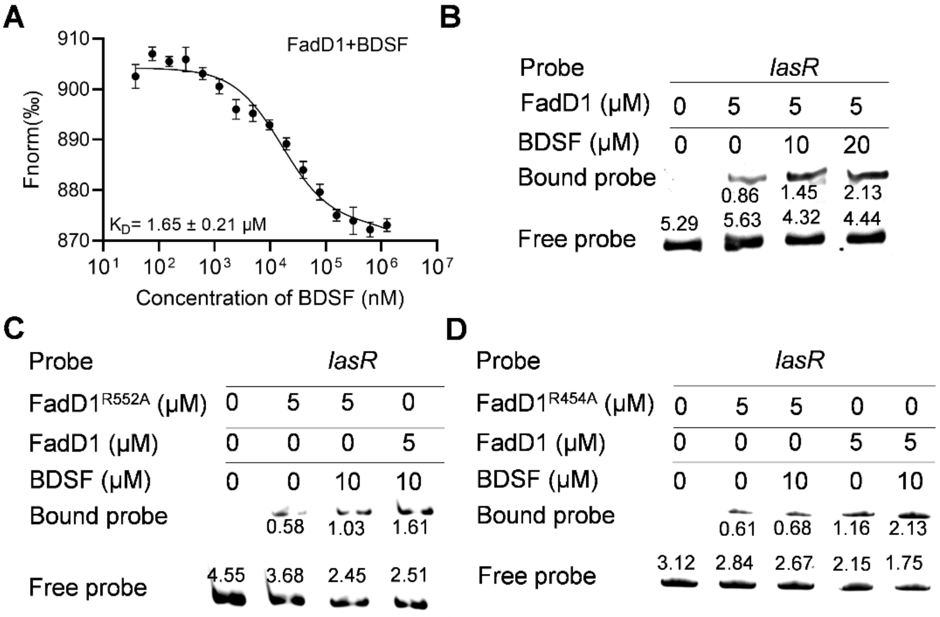


**Figure A14. Effects of BDSF on the binding of FadD1 to the promoters of the target genes.** (A) MST analysis of the binding of BDSF to the FadD1. "Fnorm (‰)" indicates the fluorescence time trace changes in the MST response. EMSA analysis of the *in vitro* binding of FadD1 (B), FadD1$^{R552A}$ (C) and FadD1$^{R454A}$ (D) to the promoter of *lasR* with the addition of different amounts of BDSF. The EMSA experiments were performed three times, and representative images from one experiment are shown.

**Table S1.** List of genes differentially expressed in the *fadD1* mutant compared to the wild-type strain ($|Log_2$-fold change$| \geq 1.0$). Significantly differentially expressed genes were determined by Cufflinks after Benjamini-Hochberg correction. The fold-change is the ratio of the mutant FPKM to the wild-type FPKM

| Gene ID[a] | Fold change | Description |
|---|---|---|
| PA1202 | -3.8878 | probable hydrolase |
| PA2432 | -3.7973 | *bexR*, bistable expression regulator, BexR |
| PA5421 | -3.2006 | *fdhA*, glutathione-independent formaldehyde dehydrogenase |
| PA1887 | -3.0699 | hypothetical protein |
| PA2696 | -3.0467 | probable transcriptional regulator |
| PA3417 | -2.9992 | probable pyruvate dehydrogenase E1 component, alpha subunit |
| PA5420 | -2.9658 | *purU2*, formyltetrahydrofolate deformylase |
| PA2698 | -2.9335 | probable hydrolase |
| PA0572 | -2.9105 | hypothetical protein |
| PA3361 | -2.8829 | *lecB*, fucose-binding lectin PA-IIL |
| PA1888 | -2.8731 | hypothetical protein |
| PA5418 | -2.8384 | *soxA*, sarcosine oxidase alpha subunit |
| PA4986 | -2.8383 | probable oxidoreductase |
| PA5400 | -2.809 | probable electron transfer flavoprotein alpha subunit |
| PA1421 | -2.7865 | *gbuA*, guanidinobutyrase |
| PA5419 | -2.6934 | *soxG*, sarcosine oxidase gamma subunit |

| | | |
|---|---|---|
| PA3416 | -2.6535 | probable pyruvate dehydrogenase E1 component, beta chain |
| PA5416 | -2.622 | *soxB*, sarcosine oxidase beta subunit |
| PA2762 | -2.6047 | hypothetical protein |
| PA2937 | -2.511 | hypothetical protein |
| PA5399 | -2.4972 | *dgcB*, DgcB, Dimethylglycine catabolism |
| PA5401 | -2.4746 | hypothetical protein |
| PA3431 | -2.4124 | conserved hypothetical protein |
| PA1419 | -2.4067 | probable transporter |
| PA5398 | -2.3686 | *dgcA*, DgcA, Dimethylglycine catabolism |
| PA3432 | -2.3565 | hypothetical protein |
| PA3415 | -2.3541 | probable dihydrolipoamide acetyltransferase |
| PA2570 | -2.3248 | *lecA*, LecA |
| PA5411 | -2.3186 | *gbcB*, GbcB |
| PA5397 | -2.2613 | hypothetical protein |
| PA5396 | -2.2522 | hypothetical protein |
| PA2431 | -2.1683 | hypothetical protein |
| PA0106 | -2.1425 | *coxA*, cytochrome c oxidase, subunit I |
| PA5410 | -2.1265 | *gbcA*, GbcA |
| PA1418 | -2.1174 | probable sodium:solute symport protein |
| PA1432 | -2.1072 | autoinducer synthesis protein LasI |

| | | |
|---|---|---|
| PA4985 | -2.0785 | Uncharacterized protein |
| PA5482 | -2.0674 | hypothetical protein |
| PA0451 | -2.0273 | conserved hypothetical protein |
| PA4305 | -2.0194 | *rcpC*, RcpC |
| PA0452 | -2.0178 | probable stomatin-like protein |
| PA2430 | -1.9979 | conserved hypothetical protein |
| PA3126 | -1.9799 | *ibpA*, heat-shock protein IbpA |
| PA5181 | -1.934 | probable oxidoreductase |
| PA1430 | -1.9267 | transcriptional regulator LasR |
| PA3418 | -1.8871 | *ldh*, leucine dehydrogenase |
| PA0108 | -1.8643 | *coIII*, cytochrome c oxidase, subunit III |
| PA4306 | -1.8619 | *flp*, Type IVb pilin, Flp |
| PA1731 | -1.8605 | conserved hypothetical protein |
| PA4377 | -1.8371 | hypothetical protein |
| PA3236 | -1.8325 | *betX*, BetX |
| PA0105 | -1.8316 | *coxB*, cytochrome c oxidase, subunit II |
| PA4303 | -1.8304 | *tadZ*, TadZ |
| PA4596 | -1.8156 | *esrC*, EsrC |
| PA1041 | -1.8101 | probable outer membrane protein precursor |
| PA5381 | -1.7887 | hypothetical protein |

| | | |
|---|---|---|
| PA1931 | -1.7697 | probable ferredoxin |
| PA4738 | -1.7295 | conserved hypothetical protein |
| PA5379 | -1.7285 | *sdaB*, L-serine dehydratase |
| PA4304 | -1.7261 | *rcpA*, RcpA |
| PA5376 | -1.7128 | *cbcV*, CbcV |
| PA3628 | -1.6901 | putative esterase |
| PA1289 | -1.6648 | hypothetical protein |
| PA3677 | -1.6589 | *mexJ*, MexJ |
| PA5348 | -1.6464 | probable DNA-binding protein |
| PA0788 | -1.6258 | hypothetical protein |
| PA5481 | -1.6138 | hypothetical protein |
| PA2433 | -1.6058 | hypothetical protein |
| PA4294 | -1.6042 | hypothetical protein |
| PA1176 | -1.5993 | *napF*, ferredoxin protein NapF |
| PA0177 | -1.5869 | probable purine-binding chemotaxis protein |
| PA0587 | -1.5778 | conserved hypothetical protein |
| PA4293 | -1.5743 | *pprA*, two-component sensor PprA |
| PA3006 | -1.5702 | *psrA*, transcriptional regulator PsrA |
| PA0195 | -1.5609 | *pntAB*, putative NAD(P) transhydrogenase, subunit alpha part 2 |
| PA1137 | -1.5525 | probable oxidoreductase |

| | | |
|---|---|---|
| PA4393 | -1.5489 | AmpG protein |
| PA3678 | -1.545 | *mexL*, MexL |
| PA0110 | -1.5398 | hypothetical protein |
| PA2024 | -1.5236 | probable ring-cleaving dioxygenase |
| PA3568 | -1.5094 | probable acetyl-coa synthetase |
| PA0588 | -1.5046 | conserved hypothetical protein |
| PA4296 | -1.5018 | *pprB*, two-component response regulator, PprB |
| PA2190 | -1.4967 | conserved hypothetical protein |
| PA1730 | -1.4952 | conserved hypothetical protein |
| PA2566 | -1.4943 | Uncharacterized protein |
| PA0905 | -1.4917 | *rsmA*, RsmA |
| PA0178 | -1.4711 | probable two-component sensor |
| PA1223 | -1.4696 | probable transcriptional regulator |
| PA4877 | -1.4686 | hypothetical protein |
| PA2072 | -1.4558 | conserved hypothetical protein |
| PA5378 | -1.4473 | *cbcX*, CbcX |
| PA3692 | -1.4393 | *lptF*, Lipotoxon F, LptF |
| PA3629 | -1.4382 | *adhC*, alcohol dehydrogenase class III |
| PA2166 | -1.4215 | hypothetical protein |
| PA0743 | -1.4173 | probable 3-hydroxyisobutyrate dehydrogenase |

| | | |
|---|---|---|
| PA2249 | -1.4138 | *bkdB*, branched-chain alpha-keto acid dehydrogenase (lipoamide component) |
| PA3973 | -1.4014 | probable transcriptional regulator |
| PA5377 | -1.3982 | *cbcW*, CbcW |
| PA2573 | -1.397 | probable chemotaxis transducer |
| PA0038 | -1.3958 | hypothetical protein |
| PA0059 | -1.393 | *osmC*, osmotically inducible protein OsmC |
| PA5213 | -1.3815 | *gcvP1*, glycine cleavage system protein P1 |
| PA1174 | -1.3782 | *napA*, periplasmic nitrate reductase protein NapA |
| PA3311 | -1.3782 | *nbdA*, NbdA |
| PA0866 | -1.3745 | *aroP2*, aromatic amino acid transport protein AroP2 |
| PA1930 | -1.3645 | probable chemotaxis transducer |
| PA4915 | -1.3641 | probable chemotaxis transducer |
| PA3478 | -1.3491 | *rhlB*, rhamnosyltransferase chain B |
| PA2564 | -1.3272 | hypothetical protein |
| PA4387 | -1.3229 | conserved hypothetical protein |
| PA1860 | -1.3102 | hypothetical protein |
| PA2248 | -1.305 | *bkdA2*, 2-oxoisovalerate dehydrogenase (beta subunit) |
| PA1324 | -1.3047 | hypothetical protein |
| PA0535 | -1.2971 | probable transcriptional regulator |
| PA3477 | -1.2931 | transcriptional regulator RhlR |

| | | |
|---|---|---|
| PA1999 | -1.293 | *dchA*, dehydrocarnitine CoA transferase, DchA |
| PA0355 | -1.2919 | *pfpI*, protease PfpI |
| PA0534 | -1.2904 | *pauB1*, FAD-dependent oxidoreductase |
| PA0179 | -1.2882 | probable two-component response regulator |
| PA3691 | -1.2872 | hypothetical protein |
| PA0586 | -1.2864 | conserved hypothetical protein |
| PA2699 | -1.2778 | hypothetical protein |
| PA2504 | -1.2571 | hypothetical protein |
| PA4739 | -1.251 | conserved hypothetical protein |
| PA2779 | -1.2472 | hypothetical protein |
| PA5546 | -1.2443 | conserved hypothetical protein |
| PA1327 | -1.2403 | probable protease |
| PA4573 | -1.2387 | hypothetical protein |
| PA5446 | -1.2387 | hypothetical protein |
| PA3476 | -1.2297 | autoinducer synthesis protein RhlI |
| PA1208 | -1.2269 | conserved hypothetical protein |
| PA0208 | -1.225 | *mdcA*, malonate decarboxylase alpha subunit |
| PA2362 | -1.2197 | *dotU3*, DotU3 |
| PA0996 | -1.2053 | *pqsA*, anthranilate-CoA ligase PqsA |
| PA4341 | -1.204 | probable transcriptional regulator |

| | | |
|---|---|---|
| PA3986 | -1.2005 | hypothetical protein |
| PA2247 | -1.1969 | *bkdA1*, 2-oxoisovalerate dehydrogenase (alpha subunit) |
| PA4236 | -1.1952 | catalase KatA |
| PA2746 | -1.1938 | hypothetical protein |
| PA1728 | -1.1911 | hypothetical protein |
| PA0196 | -1.1904 | *pntB*, pyridine nucleotide transhydrogenase, beta subunit |
| PA0052 | -1.1872 | hypothetical protein |
| PA4311 | -1.1752 | conserved hypothetical protein |
| PA3676 | -1.174 | *mexK*, MexK |
| PA4925 | -1.1727 | conserved hypothetical protein |
| PA3275 | -1.171 | conserved hypothetical protein |
| PA1415 | -1.1702 | hypothetical protein |
| PA1523 | -1.1681 | *xdhB*, xanthine dehydrogenase |
| PA2759 | -1.1513 | hypothetical protein |
| PA0457 | -1.1489 | hypothetical protein |
| PA4614 | -1.1487 | large-conductance mechanosensitive channel |
| PA2000 | -1.1485 | *dchB*, dehydrocarnitine CoA transferase, DchB |
| PA4312 | -1.1418 | conserved hypothetical protein |
| PA4914 | -1.1412 | probable transcriptional regulator |
| PA3089 | -1.141 | hypothetical protein |

| | | |
|---|---|---|
| PA3952 | -1.1389 | hypothetical protein |
| PA0256 | -1.1361 | hypothetical protein |
| PA0180 | -1.1356 | *cttP*, chemotactic transducer for trichloroethylene [positive chemotaxis], CttP |
| PA2226 | -1.1304 | *qsrO*, QsrO |
| PA4657 | -1.1297 | hypothetical protein |
| PA3723 | -1.128 | probable FMN oxidoreductase |
| PA3459 | -1.1218 | probable glutamine amidotransferase |
| PA4881 | -1.1211 | hypothetical protein |
| PA3569 | -1.1191 | *mmsB*, 3-hydroxyisobutyrate dehydrogenase |
| PA1323 | -1.1176 | hypothetical protein |
| PA3424 | -1.1169 | hypothetical protein |
| PA4913 | -1.111 | probable binding protein component of ABC transporter |
| PA4781 | -1.1033 | cyclic di-GMP phosphodiesterase |
| PA3160 | -1.1002 | *wzz*, O-antigen chain length regulator |
| PA3351 | -1.0963 | protein FlgM |
| PA5477 | -1.0888 | hypothetical protein |
| PA4027 | -1.0844 | hypothetical protein |
| PA3972 | -1.0777 | probable acyl-CoA dehydrogenase |
| PA2534 | -1.0696 | probable transcriptional regulator |
| PA3787 | -1.0569 | conserved hypothetical protein |

| | | |
|---|---|---|
| PA2572 | -1.056 | probable two-component response regulator |
| PA0798 | -1.0547 | *pmtA*, phospholipid methyltransferase |
| PA3957 | -1.054 | probable short-chain dehydrogenase |
| PA4209 | -1.0514 | *phzM*, probable phenazine-specific methyltransferase |
| PA4876 | -1.0484 | *osmE*, osmotically inducible lipoprotein OsmE |
| PA2363 | -1.0472 | *hsiJ3*, HsiJ3 |
| PA4112 | -1.0468 | probable sensor/response regulator hybrid |
| PA2571 | -1.0439 | probable two-component sensor |
| PA1249 | -1.0427 | *aprA*, alkaline metalloproteinase precursor |
| PA3570 | -1.0413 | *mmsA*, methylmalonate-semialdehyde dehydrogenase |
| PA5359 | -1.0334 | hypothetical protein |
| PA2250 | -1.0333 | *lpdV*, lipoamide dehydrogenase-Val |
| PA2002 | -1.0304 | conserved hypothetical protein |
| PA3709 | -1.0304 | probable major facilitator superfamily (MFS) transporter |
| PA2227 | -1.0285 | *vqsM*, AraC-type transcriptional regulator VqsM |
| PA1003 | -1.0262 | transcriptional regulator MvfR |
| PA0997 | -1.0259 | 2-heptyl-4(1H)-quinolone synthase subunit, PqsB |
| PA0746 | -1.0246 | probable acyl-CoA dehydrogenase |
| PA1761 | -1.0229 | hypothetical protein |
| PA0543 | -1.0215 | hypothetical protein |

| | | |
|---|---|---|
| PA5409 | -1.0211 | hypothetical protein |
| PA4211 | -1.0184 | probable phenazine biosynthesis protein |
| PA2001 | -1.0162 | *atoB*, acetyl-CoA acetyltransferase |
| PA4880 | -1.0141 | probable bacterioferritin |
| PA0122 | -1.0105 | *rahU*, rahU |
| PA4702 | -1.0036 | hypothetical protein |
| PA2290 | -1.0031 | *gcd*, glucose dehydrogenase |
| PA0547 | -1.0028 | probable transcriptional regulator |
| PA0692 | -1.0025 | hypothetical protein |
| PA3865 | -1.0019 | probable amino acid binding protein |
| PA5212 | -1.0013 | hypothetical protein |
| PA0714 | -1.0007 | hypothetical protein |
| PA1588 | 1.0129 | *sucC*, succinyl-CoA synthetase beta chain |
| PA1480 | 1.0194 | *ccmF*, cytochrome C-type biogenesis protein CcmF |
| PA4258 | 1.022 | *rplV*, 50S ribosomal protein L22 |
| PA1245 | 1.0374 | hypothetical protein |
| PA2840 | 1.0443 | probable ATP-dependent RNA helicase |
| PA2116 | 1.047 | conserved hypothetical protein |
| PA1155 | 1.0763 | *nrdB*, NrdB, tyrosyl radical-harboring component of class Ia ribonucleotide reductase |
| PA4740 | 1.0824 | *pnp*, polyribonucleotide nucleotidyltransferase |

| | | |
|---|---|---|
| PA3651 | 1.0852 | *cdsA*, phosphatidate cytidylyltransferase |
| PA1585 | 1.1098 | *sucA*, 2-oxoglutarate dehydrogenase (E1 subunit) |
| PA1123 | 1.1374 | hypothetical protein |
| PA4257 | 1.1647 | *rpsC*, 30S ribosomal protein S3 |
| PA5374 | 1.1669 | *betI*, transcriptional regulator BetI |
| PA5547 | 1.1835 | conserved hypothetical protein |
| PA3190 | 1.2115 | probable binding protein component of ABC sugar transporter |
| PA4442 | 1.2543 | *cysN*, ATP sulfurylase GTP-binding subunit/APS kinase |
| PA4855 | 1.2773 | *purD*, phosphoribosylamine--glycine ligase |
| PA3183 | 1.3552 | *zwf*, glucose-6-phosphate 1-dehydrogenase |
| PA0280 | 1.7592 | *cysA*, sulfate transport protein CysA |
| PA3187 | 1.813 | probable ATP-binding component of ABC transporter |
| PA5373 | 1.8779 | *betB*, betaine aldehyde dehydrogenase |
| PA5372 | 2.0769 | *betA*, choline dehydrogenase |
| PA3446 | 2.8022 | conserved hypothetical protein |

**Table S2.** Potential target genes regulated by FadD1 identified via ChIP-Seq analysis

| Gene ID | Gene name | Sumit | Binding sequence | Description |
|---|---|---|---|---|
| PA0122 | *PA0122* | 141586 | AGGACGG | hypothetical protein |
| PA0128 | *PA0128* | 145705 | AGGACGG | conserved hypothetical protein |
| PA0457 | *PA0457* | 515340 | AGGACGG | hypothetical protein |
| PA0547 | *PA0547* | 606123 | AGGAGTG | probable transcriptional regulator |
| PA0572 | *PA0572* | 628172 | CGGACGA | hypothetical protein |
| PA0642 | *PA0642* | 698536 | AGGACGG | hypothetical protein |
| PA0692 | *PA0692* | 763530 | AGGGCGG | hypothetical protein |
| PA0714 | *PA0714* | 784688 | AGGACGC | hypothetical protein |
| PA0973 | *oprL* | 1056935 | AGGACGG | peptidoglycan associated lipoprotein OprL |
| PA1245 | *PA1245* | 1349093 | AGGTCGG | hypothetical protein |
| PA1430 | *lasR* | 1557923 | AGGACGG | transcriptional regulator LasR |
| PA2840 | *PA2840* | 3195997 | AGGTCGA | probable ATP-dependent RNA helicase |
| PA3112 | *accD* | 3493362 | AGGACGG | acetyl-CoA carboxylase beta subunit |
| PA3351 | *flgM* | 3762671 | TGGCCGG | protein FlgM |
| PA3865 | *PA3865* | 4325467 | AGGAGGG | probable amino acid binding protein |
| PA4209 | *phzM* | 4713529 | AGGTCGC | phenazine-specific methyltransferase |
| PA4211 | *phzB1* | 4714444 | AGGACGG | probable phenazine biosynthesis protein |
| PA4236 | *katA* | 4752079 | AGGAGGG | catalase |

| | | | | |
|---|---|---|---|---|
| PA4393 | *ampG* | 4922173 | AGGACGC | AmpG protein |
| PA4614 | *mscL* | 5171963 | AGGTCGG | large-conductance mechanosensitive channel |
| PA4914 | *PA4914* | 5513446 | AGGACAG | transcriptional regulator |
| PA5045 | *ponA* | 5681055 | AGGACGG | penicillin-binding protein 1A |
| PA5411 | *gbcB* | 6088177 | AGGACAG | GbcB protein |
| PA5547 | *PA5547* | 6240651 | AAGACGG | hypothetical protein |

**Table S3.** The contact list between DNA and FadD1

| Chain 1 | Residue | Chain 2 | Residue | Distance (Å) | Interaction type |
|---|---|---|---|---|---|
| FadD1 | Asn285: ND2 | DNA | DA4: OP2 | 3.2 | Hydrogen bond |
| FadD1 | Asp288: OD2 | DNA | DC5: N4 | 3.0 | Hydrogen bond |
| FadD1 | Lys173: NZ | DNA | DC2: Pyrimidine | 4.3 | Pi-cation |
| FadD1 | Lys173: CE | DNA | DG3: N7 | 3.2 | Hydrogen bond |
| FadD1 | Pro290: O | DNA | DC1: N4 | 3.4 | Hydrogen bond |
| FadD1 | Lys294: NZ | DNA | DC1: O4’ | 3.3 | Hydrogen bond |

**Table S4.** Analysis of the homologs of FadD1 in various bacteria

| Bacteria | FadD1 homologue Accession No. | FadD1 homologue Identity (%) | Leucine zipper |
|---|---|---|---|
| ***Pseudomonas*** | | | |
| *P. aeruginosa* PAO1 | NP_251989.1 | 100 | LTGNHNILITNPRDLPSMLKDL |
| *P. nitroreducens* PSA00705 | MBG6289323.1 | 90.14 | LTGNHNILVTNPRDLASMVKEL |
| *P. nitritireducens* S00179 | MBB4863819.1 | 89.96 | LTGNHNILVTNPRDLASMVKEL |
| *P. multiresinivorans* populi | QJP08166.1 | 89.78 | LTGNHNILVTNPRDLASMVKEL |
| *P. nicosulfuronedens* LAM1902 | TLX74267.1 | 89.61 | LTGNHNILVTNPRDLASMVKEL |
| *P. panipatensis* CCM 7469 | SDH37009.1 | 87.84 | LTGNHNILVTNPRDLNALAKEL |
| *P. citronellolis* LA18T | GBL53074.1 | 87.3 | LTGNHNILVTNPRDLNAFSKEL |
| *P. knackmussii* B13 | CDF82887.1 | 86.94 | LTGNHNILVTNPRDLNALAKEL |
| *P. schmalbachii* Milli4 | MBO3276413.1 | 86.65 | LTGNHNVLVTNPRDLSAMVKEL |
| *P. oleovorans* PO_271 | RRW39041.1 | 85.94 | LIGGHNILLTNPRDLPAVVKDL |
| *P. indoloxydans* JCM 14246 | PTU80596.1 | 85.77 | LIGGHNILLTNPRDLPAVVKDL |
| *P. mendocina* S5.2 | ALN20083.1 | 85.59 | LIGGHNILLTNPRDLPSVIKDL |
| *P. toyotomiensis* | WP_059391876.1 | 85.59 | LIGGHNILLTNPRDLPAVVKDL |
| *P. guguanensis* JCM 18416 | SDO56856.1 | 85.77 | LIGGHNILLTNPRDLPAVVKDL |
| *P. delhiensis* CCM 7361 | SDH92421.1 | 86.40 | LTGNHNVLVTNPRDLNAFSKEL |
| *P. sediminis* B10D7D | QNG99969.1 | 85.94 | LIGGHNILLTNPRDLPAVVKDL |
| *P. alcaliphila* JCM 10630 | SDD42921.1 | 85.77 | LIGGHNILLTNPRDLPAVVKDL |

| | | | |
|---|---|---|---|
| *P. jinjuensis* JCM 21621 | SDO77105.1 | 85.77 | LTGNHNILVTNPRDLSAMVKEL |
| *P. composti* CCUG 5923 | SFP37007.1 | 85.41 | LIGGHNILLTNPRDLSAVVKDL |
| *P. chengduensis* | WP_247723315.1 | 85.23 | LIGGHNVLLTNPRDLPAVVKDL |
| *P. yangonensis* | WP_161866568.1 | 85.41 | LIGGHNILLTNPRDLPAVVKDL |
| *P. hydrolytica* | MCF2123116.1 | 85.59 | LIGGHNILLTNPRDLPAVVKDL |
| *P. wenzhouensis* A20 | UFQ96462.1 | 85.05 | LIGGHNILLTNPRDLPAVVKDL |
| *P. carbonaria* CIP 111764 | CAD5107370.1 | 85.23 | LIGGHNILITNPRDLPGTVKEL |
| *P. kuykendallii* NRRL B-59562 | SDX22616.1 | 84.16 | LIGGHNILVTNPRDLPGTVKEL |
| *P. putida* CBF10-2 | OAI93987.1 | 84.16 | LIGNHNILISNPRDLPAMVKEL |
| *P. resinovorans* | WP_028626413.1 | 84.34 | LTGNHNVLISNPRDLPAMTKEL |
| *P. mangiferae* DMKU BBB3-04 | TRX73262.1 | 85.23 | LTGNHNILITNPRDLPGMVKEL |
| *P. khazarica* | WP_134676293.1 | 84.52 | LIGGHNVLLTNPRDLPSVVKDL |
| *P. alcaligenes* AVO110 | MBC9251595.1 | 86.20 | LTGNHNILVTNPRDLPGTVKEL |
| *P. cavernae* K2W31S-8 | AYC33791.1 | 84.34 | LIGGHNILITNPRDLPGTVKEL |
| *P. otitidis* MrB4 | BCA27519.1 | 84.59 | LTGNHNILISNPRDLPAMVKEL |
| *P. stutzeri* | MBF6622309.1 | 82.92 | LIGNHNILITNPRDLPAMIKDL |
| *P. flexibilis* JCM 14085 | KHO66267.1 | 84.59 | LTGNHNVLITNPRDLPAMIKDL |
| *P. xionganensis* R-22-3 w-18 | MVW76420.1 | 84.34 | LIGGHNVLITNPRDLPAMTKDL |
| *P. frederiksbergensis* SI8 | KHK62161.1 | 83.27 | LIGNHNILISNPRDLSAMVKEL |
| *P. fluorescens* NT0133 | KJH87468.1 | 83.1 | LIGNHNILISNPRDLSAMVKEL |
| *P. tohonis* | WP_236207986.1 | 84.77 | LTGNHNVLISNPRDLPTMVKEL |

| | | | |
|---|---|---|---|
| *P. rhodesiae* DLA23 | QVN00525.1 | 83.99 | LIGNHNILISNPRDLPAMVKEL |
| *P. mediterranea* EDOX | MBL0841323.1 | 83.45 | LIGNHNILISNPRDLPAMVKEL |
| *P. thermotolerans* | WP_017938723.1 | 83.27 | LIGGHNILITNPRDLPAMVKEL |
| *P. guryensis* SR9 | MBB1520630.1 | 84.34 | LTGNHNILITNPRDLPGTVKEL |
| *P. trivialis* IHBB745 | AKS08436.1 | 83.45 | LIGNHNILISNPRDLPAMVKEL |
| *P. brassicacearum* Bi06 | CAH0323621.1 | 83.63 | LIGNHNILISNPRDLPAMVKEL |
| *P. taiwanensis* xwS3 | NWL75364.1 | 83.45 | LTGNHNILISNPRDLPAMTKEL |
| *P. laurylsulfatiphila* AP3_16 | PPK38601.1 | 83.45 | LIGNHNILISNPRDLPAMVKEL |
| *P. laurylsulfativorans* AP3_22 | POF39209.1 | 83.81 | LIGNHNILISNPRDLPAMVKEL |
| *P. costantinii* 21815972 | NVZ21787.1 | 83.45 | LIGNHNILISNPRDLPAMVKEL |
| *P. lalkuanensis* PE08 | QEY64321.1 | 83.27 | LTGNHNILISNPRDLPAMTREL |
| *P. campi* S1-A32-2 | QKE62586.1 | 84.34 | LTGNHNILITNPRDLPGTVKEL |
| *P. cavernicola* K1S02-6 | RJG14059.1 | 83.45 | LIGSHNILITNPRDLPAMIKEL |
| *P. chlororaphis* EA105 | KHA70093.1 | 83.27 | LIGNHNILISNPRDLPAMVKEL |
| *P. songnenensis* L103 | RYJ59864.1 | 82.74 | LSGNHNILISNPRDLPAMIKDL |
| *P. izuensis* | WP_160106207.1 | 83.45 | LIGNHNILISNPRDLPAMVKEL |
| *P. corrugate* RM1-1-4 | AOE64340.1 | 83.1 | LIGNHNILISNPRDLPAMVKEL |
| *P. ogarae* | WP_204128328.1 | 83.45 | LIGNHNILISNPRDLPAMVKEL |
| *P. thivervalensis* SC5 | AXA55291.1 | 83.45 | LIGNHNILISNPRDLSAMVKEL |
| *P. synxantha* R6-28-08 | AZE68831.1 | 83.27 | LIGNHNILISNPRDLPAMVKEL |
| *P. helmanticensis* BIGb0525 | TDV44006.1 | 83.27 | LIGNHNILISNPRDLPAMVKEL |

| | | | |
|---|---|---|---|
| *P. kilonensis* | WP_024616910.1 | 83.45 | LIGNHNILISNPRDLPAMVKEL |
| *P. moorei* CCUG 53114 | KAB0495447.1 | 83.45 | LIGNHNILISNPRDLPAMVKEL |
| *P. lurida* 114A4 | MBA1294871.1 | 83.63 | LIGNHNILISNPRDLPAMVKEL |
| *P. mohnii* BOE200 | MBH8614930.1 | 83.45 | LIGNHNILISNPRDLPAMVKEL |
| *P. viridiflava* | WP_122674469.1 | 83.63 | LIGNHNILISNPRDLPAMVKEL |
| *P. mucidolens* | WP_084381059.1 | 83.63 | LIGNHNILISNPRDLSAMVKEL |
| *P. fildesensis* KG01 | KMT53924.1 | 83.63 | LIGNHNILISNPRDLPAMVKEL |
| *P. yamanorum* B4002 | NVZ92296.1 | 83.45 | LIGNHNILISNPRDLSAMVKEL |
| *P. moraviensis* 36B3 | RON99995.1 | 83.27 | LIGNHNILISNPRDLPAMVKEL |
| *P. koreensis* AB36 | RYM44580.1 | 83.27 | LIGNHNILISNPRDLPAMVKEL |
| *P. simiae* MEB105 | KIQ12695.1 | 83.45 | LIGNHNILISNPRDLPAMVKEL |
| *P. granadensis* F6_4S_P_1C | MBN6774081.1 | 83.27 | LIGNHNILISNPRDLPAMVKEL |
| *P. huaxiensis* | WP_110968545.1 | 83.45 | LIGNHNILISNPRDLPAMVKEL |
| *P. arsenicoxydans* E3 | TPG79102.1 | 83.63 | LIGNHNILISNPRDLPAMVKEL |
| *P. indica* PIC105 | PAU64194.1 | 83.81 | LNGCHNVLITNPRDLPGMIRDL |
| *P. syringae* PA-1-3B | MCF5705526.1 | 83.27 | LIGNHNILISNPRDLPAMVKEL |
| *P. gessardii* PSB00014 | MBH3425107.1 | 83.45 | LIGNHNILISNPRDLPAMVKEL |
| *P. reactans* C5002 | NWA69384.1 | 83.45 | LIGNHNILISNPRDLPAMVKEL |
| *P. libanensis* DSM 17149 | KRP43192.1 | 82.92 | LIGNHNILISNPRDLPAMVKEL |
| *P. tolaasii* 2192T | ARB28676.1 | 83.1 | LIGNHNILISNPRDLPAMVKEL |
| *P. khorasanensis* SWRI153 | MBV4485496.1 | 83.1 | LIGNHNILISNPRDLPAMVKEL |

| | | | |
|---|---|---|---|
| *P. cremoris* WS 5096 | MBC2381977.1 | 83.27 | LIGNHNILISNPRDLPAMVKEL |
| *P. cedrina* BS2981 | SDT31708.1 | 83.27 | LIGNHNILISNPRDLPAMVKEL |
| *P. glycinae* PSB00018 | MBH3408202.1 | 83.1 | LIGNHNILISNPRDLPAMVKEL |
| *P. karstica* CCM 7891 | MTD22140.1 | 82.92 | LIGNHNILISNPRDLPAMVKEL |
| *P. bananamidigenes* | WP_065259335.1 | 83.27 | LIGNHNILISNPRDLPAMVKEL |
| *P. proteolytica* CCUG 51515T | KAA8705669.1 | 82.92 | LIGNHNILISNPRDLPAMVKEL |
| *P. khavaziana* SWRI124 | MBV4481306.1 | 82.92 | LIGNHNVLISNPRDLPAMVKEL |
| *P. viciae* 11K1 | QBZ88412.1 | 82.92 | LIGNHNILISNPRDLSAMVKEL |
| *P. ekonensis* COR58 | MBV4457085.1 | 82.92 | LIGNHNILISNPRDLPAMVKEL |
| *P. farris* SWRI79 | MBV4461951.1 | 83.45 | LIGNHNILISNPRDLTAMVKEL |
| *P. grimontii* DSM 17515 | TWR63689.1 | 83.27 | LIGNHNILISNPRDLPAMVKEL |
| *P. sputi* | WP_236177820.1 | 83.1 | LIGNHNILISNPRDLPAMVKEL |
| *P. azotoformans* | RMT64654.1 | 83.45 | LIGNHNILISNPRDLPAMVKEL |
| *P. ullengensis* UL070 | MBB2494899.1 | 84.23 | LTGNHNILVTNPRDLPGTVKEL |
| *P. vancouverensis* CCUG 49675 | KAB0500079.1 | 83.1 | LIGNHNILISNPRDLPAMVKEL |
| *P. anguilliseptica* MatS1 | MCE5362680.1 | 83.45 | LIGGHNVLITNPRDLPSMTKDL |
| *P. atagonensis* | WP_166219121.1 | 82.92 | LIGNHNILISNPRDLPAMVKEL |
| *P. mandelii* PD30 | KDD68846.1 | 83.45 | LIGNHNILISNPRDLTAMVKEL |
| *P. gozinkensis* | WP_192562140.1 | 82.92 | LIGNHNILISNPRDLPAMVKEL |
| *P. salmasensis* SWRI126 | QXH79967.1 | 83.27 | LIGNHNILISNPRDLPAMVKEL |

| | | | |
|---|---|---|---|
| *P. marginalis* pv. Marginalis ICMP 3555 | RMP05188.1 | 82.92 | LIGNHNILISNPRDLPAMVKEL |
| *P. botevensis* COW3 | MBV4476468.1 | 83.63 | LIGNHNILISNPRDLPAMVKEL |
| *P. lini* 48C10 | RON30104.1 | 83.45 | LIGNHNILISNPRDLTAMVKEL |
| *P. nabeulensis* E10B | TFY90391.1 | 83.45 | LIGNHNILISNPRDLPAMVKEL |
| *P. gregormendelii* LMG 28632 | MBN3965914.1 | 82.74 | LIGNHNILISNPRDLPAMVKEL |
| *P. protegens* | WP_047288658.1 | 82.56 | LMGNHNILISNPRDLPAMVKEL |
| *P. saponiphila* DSM 9751 | SEB59069.1 | 82.92 | LMGNHNILISNPRDLPAMVKEL |
| *P. carnis* UCD_MED3 | MBV2082169.1 | 83.45 | LIGNHNILISNPRDLPAMVKEL |
| *P. taeanensis* MS-3 | KFX69438.1 | 82.21 | LIGNHNVLITNPRDLPGMVKDL |
| *P. tehranensis* SWRI196 | MBC3347638.1 | 82.56 | LIGNHNILISNPRDLPAMVKEL |
| *P. reinekei* CCUG 53116 | KAB0487234.1 | 82.38 | LIGNHNILISNPRDLPAMVKEL |
| *P. jessenii* E2333 | RDL23893.1 | 82.74 | LIGNHNILISNPRDLPAMVKEL |
| *P. saudiphocaensis* AGROB56 | MBE7926993.1 | 83.15 | LIGNHNILITNPRDLPAMIKDL |
| *P. lactis* | MBA6041862.1 | 83.27 | LIGNHNILISNPRDLPAMVKEL |
| *P. extremaustralis* DSM 17835 | TWS01464.1 | 83.36 | LIGNHNILISNPRDLPSMVKEL |
| *P. dryadis* P6B | TBU91931.1 | 82.38 | LIGNHNVLITNPRDLPTMVKDL |
| *P. kairouanensis* KC12 | TFY89268.1 | 82.74 | LIGNHNILISNPRDLPAMVKEL |
| *P. orientalis* DSM 17489 | KRP64839.1 | 83.1 | LIGNHNILISNPRDLPAMVKEL |
| *P. gingeri* J4002 | NWE44669.1 | 82.38 | LIGNHNILISNPRDLSAMVKEL |
| *P. muyukensis* COW39 | QXH34539.1 | 83.45 | LIGNHNVLISNPRDLPAMVKEL |

| | | | |
|---|---|---|---|
| *P. panacis* UKR2 | KAA6170636.1 | 82.92 | LIGNHNILISNPRDLPAMVKEL |
| *P. kunmingensis* | WP_104096911.1 | 81.49 | LSGNHNILISNPRDLPAMIKDL |
| *P. pisciculturae* P115 | MBF6030105.1 | 82.74 | LIGNHNILISNPRDLPAMVKEL |
| *P. shirazensis* SWRI56 | MBV4503001.1 | 82.94 | LIGNHNVLISNPRDLPAMVKEL |
| *P. spelaei* CCM 7893 | MUF03042.1 | 82.21 | LIGNHNILISNPRDLPAMVKEL |
| *P. fuscovaginae* LMG 2158 | SEI07384.1 | 82.03 | LLGNHNILISNPRDLPAMVKEL |
| *P. alkylphenolica* Neo | QGW78809.1 | 81.67 | LIGNHNILISNPRDLPAMVKEL |
| *P. xanthosomae* COR54 | QXH45613.1 | 82.74 | LIGNHNVLISNPRDLPAMVKEL |
| *P. poae* 29G9 | ROM52574.1 | 82.56 | LIGNHNILISNPRDLPAMVKEL |
| *P. lopnurensis* AL-54 | MBE7376606.1 | 81.49 | LSGNHNILITNPRDLPAMIKDL |
| *P. parafulva* CRS01-1 | AIZ32078.1 | 82.56 | LIGNHNVLISNPRDLPAMVKEL |
| *P. mosselii* SC006 | TLP62988.1 | 82.74 | LIGNHNVLISNPRDLPAMVKEL |
| *P. fakonensis* COW40 | QXH50194.1 | 82.74 | LIGNHNVLISNPRDLPAMVKEL |
| *P. veronii* S4_EA_3b | MBI6555778.1 | 83.18 | LIGNHNVLISNPRDLPAMVKEL |
| *P. reidholzensis* CCOS 865 | SYX91533.1 | 82.92 | LIGNHNVLISNPRDLPAMVKEL |
| *P. vranovensis* 15D11 | ROL74871.1 | 81.49 | LIGSHNILISNPRDLPAMVKEL |
| *P. Antarctica* PAMC 27494 | ANF87948.1 | 82.74 | LIGNHNILISNPRDLPAMVKEL |
| *P. cichorii* | WP_200992706.1 | 82.21 | LIGNHNILISNPRDLPAMVKEL |
| *P. insulae* | WP_204915683.1 | 83.27 | LIGAHNILVTNPRDLPGTVKEL |
| *P. shirazica* | WP_238066809.1 | 82.38 | LIGNHNVLISNPRDLPAMVKEL |
| *P. fluvialis]* | WP_184681766.1 | 81.67 | LIGGHNVLISNPRDLPAMIKDL |

| | | | |
|---|---|---|---|
| *P. urmiensis* | WP_186557723.1 | 82.92 | LIGNHNVLISNPRDLPAMVKEL |
| *P. guineae* | WP_090241499.1 | 83.15 | LIGGHNVLITNPRDLPAMTKDL |
| *P. entomophila* | WP_248921020.1 | 82,92 | LIGNHNILISNPRDLPAMVKEL |
| *P. oryzicola* | WP_186676204.1 | 82.21 | LIGNHNVLISNPRDLPAMVKEL |
| *P. batumici* | WP_040070254.1 | 81.85 | LSGNHNILISNPRDLSSMVKEL |
| *P. monteilii* | WP_119369154.1 | 82.21 | LIGNHNVLISNPRDLPAMVKEL |
| *P. caspiana* | WP_087270693.1 | 81.49 | LIGNHNILISNPRDLPAMVKEL |
| *P. soli* | WP_023631539.1 | 82.21 | LIGNHNVLISNPRDLPAMVKEL |
| *P. oryziphila* | WP_125859601.1 | 82.21 | LIGNHNILISNPRDLPAMVKEL |
| *P. capeferrum* | WP_181128491.1 | 82.03 | LAGNHNVLISNPRDLPAMVKEL |
| *P. azotifigens* | WP_181069361.1 | 81.11 | LIGAHNVLITNPRDLPAFVKDL |
| *P. salomonii* | RMQ90046.1 | 81.85 | LIGNHNILISNPRDLPAMVKEL |
| *P. lemoignei* | SQF99988.1 | 80.96 | LIGSHNILISNPRDLPAMVKEL |
| *P. foliumensis* | WP_187522506.1 | 81.67 | LIGNHNILISNPRDLPAMVKEL |
| *P. laurentiana* | WP_163935429.1 | 81.32 | LIGSHNILISNPRDLPAMVKEL |
| *P. faucium* | WP_236237620.1 | 82.03 | LIGNHNVLISNPRDLPAMVKEL |
| *P. japonica* | WP_042126533.1 | 80.60 | LIGNHNILISNPRDLPAMVKEL |
| *P. lalucatii* | WP_213640783.1 | 82.62 | LIGNHNVLITNPRDLPAMVKDL |
| *P. flavescens* | WP_179537669.1 | 82.21 | NA |
| *P. bohemica* | WP_110947979.1 | 81.32 | LIGNHNILISNPRDLSAMVKEL |
| *P. asturiensis* | WP_073170226.1 | 80.78 | LIGNHNVLISNPRDLPAMVKEL |

| | | | |
|---|---|---|---|
| *P. graminis* | WP_074891667.1 | 81.14 | LSGNHNVLISNPRDLSAMVKEL |
| *P. tructae* | WP_130263645.1 | 80.96 | LIGSHNILISNPRDLPAMVKEL |
| *P. endophytica* | WP_055101100.1 | 80.96 | LLGNHNVLISNPRDLPAMVKEL |
| *P. punonensis* | WP_073265773.1 | 82.21 | NA |
| *P. saxonica* | WP_146426243.1 | 81.49 | LLGNHNILISNPRDLPAMVKEL |
| *P. edaphica* | WP_177033300.1 | 81.49 | LIGNHNILISNPRDLPAMVKEL |
| *P. daroniae* | WP_131193191.1 | 81.85 | NA |
| *P. bacterium* | MBH2036460.1 | 81.49 | LIGNHNLLVTNPRDLPGMIKDL |
| *P. lundensis* | WP_169880254.1 | 81.49 | LIGNHNVLISNPRDLPAMVKEL |
| *P. kirkiae* | WP_131183650.1 | 81.14 | NA |
| *P. balearica* | WP_061338326.1 | 81.72 | LIGAHNILITNPRDLPTMVKDL |
| *P. juntendi* | WP_182366232.1 | 81.85 | LIGNHNVLISNPRDLPAMVKEL |
| *P. floridensis* | WP_083184351.1 | 80.60 | LIGNHNVLISNPRDLPAMVKEL |
| *P. lutea* | WP_191944493.1 | 80.78 | LSGNHNVLISNPRDLSAMVKEL |
| *P. versuta* | WP_060694566.1 | 80.60 | LLGNHNVLISNPRDLPAMVKEL |
| *P. jilinensis* | WP_119701379.1 | 80.43 | NA |
| *P. helleri* | WP_153378413.1 | 80.78 | LLGNHNILISNPRDLPAMVKEL |
| *P. segetis* | WP_089359276.1 | 80.78 | LIGAHNILITNPRDLPAVVNDL |
| *P. taetrolens* | WP_048379042.1 | 80.96 | LIGNHNVLISNPRDLSAMVKEL |
| *P. fragi* | WP_095029855.1 | 8.43 | LLGNHNVLISNPRDLPAMVKEL |
| *P. straminea* | WP_093502797.1 | 80.96 | NA |

| | | | |
|---|---|---|---|
| *P. avellanae* | WP_024697501.1 | 80.60 | LIGNHNILISNPRDLPSMVKEL |
| *P. mangrovi* | WP_108106740.1 | 80.43 | NA |
| *P. alliivorans* | WP_184320742.1 | 80.25 | LIGNHNVLISNPRDLPAMVKEL |
| *P. fulva* | WP_182139659.1 | 81.49 | LIGNHNVLISNPRDLSAMVKEL |
| *P. urumqiensis* | WP_120993685.1 | 79.89 | LIGAHNVLITNPRDLAAFTKDL |
| *P. amygdali* | WP_054097109.1 | 79.18 | LLGNHNILISNPRDLPAMVKEL |
| *P. savastanoi* | WP_122218246.1 | 79.18 | LLGNHNILISNPRDLPAMVKEL |
| *P. congelans* | WP_096125679.1 | 79.00 | LLGNHNILISNPRDLPAMVKEL |
| *P. agarici* | WP_060782680.1 | 80.07 | LIGNHNILISNPRDLSAMVKEL |
| *P. nanhaiensis* | WP_223655554.1 | 78.07 | NA |
| *P. marincola* | WP_150548657.1 | 78.43 | NA |
| *P. luteola* | WP_125888235.1 | 78.14 | LLGNHNVLITNPRDLPTMIKDL |
| *P. sagittaria* | WP_092431855.1 | 77.78 | NA |
| *P. zeshuii* | WP_181123557.1 | 78.14 | LLGNHNVLITNPRDLPTMIKDL |
| *P. asuensis* | WP_188865694.1 | 77.60 | LLGNHNVLITNPRDLPTMIKDL |
| *P. duriflava* | WP_145140531.1 | 78.14 | LLGNHNVLITNPRDLPAMIKDL |
| *P. typographi* | WP_190425339.1 | 77.76 | LMGNHNVLITNPRDLPAMTKEL |
| *P. pohangensis* | WP_090194462.1 | 76.51 | NA |
| *P. profundi* | WP_150301230.1 | 75.27 | NA |
| *P. saliphila* | WP_150303759.1 | 75.27 | NA |
| *P. matsuisoli* | WP_188982677.1 | 72.86 | NA |

| | | | |
|---|---|---|---|
| *P. inefficax* | MCM8914657.1 | 82.55 | LIGNHNVLISNPRDLPAMVKEL |
| *P. guangdongensis* | WP_090213733.1 | 73.04 | NA |
| *P. oryzae* | WP_090349359.1 | 74.19 | NA |
| *P. psychrotolerans* | WP_058760915.1 | 72.81 | NA |
| *P. oryzihabitans* | WP_241809649.1 | 72.81 | NA |
| *P. cyclaminis* | WP_193901953.1 | 83.08 | NA |
| *P. farsensis* | WP_186536971.1 | 62.81 | NA |
| *P. defluvii* | WP_065760699.1 | 62.21 | NA |
| *P. maumuensis* | WP_217868799 | 61.14 | NA |
| *P. brassicae* | NER62307.1 | 61.68 | NA |
| *P. peli* | WP_090248416.1 | 61.85 | NA |
| *P. asiatica* | WP_137164607.1 | 61.32 | NA |
| *P. umsongensis* | WP_168757347.1 | 60.07 | NA |
| *P. bharatica* | WP_009405195.1 | 61.14 | NA |
| *P. uvaldensis* | WP_232776210.1 | 60.07 | NA |
| *P. vlassakiae* | WP_186603409.1 | 60.78 | NA |
| *P. borbori* | WP_090498438.1 | 61.68 | NA |
| *P. urethralis* | WP_176507878.1 | 60.61 | NA |
| *P. pharyngis* | WP_236190925.1 | 59.61 | NA |
| *P. monsensis* | WP_186746659.1 | 59.79 | NA |
| *P. tremae* | WP_236513726.1 | 60.07 | NA |

| | | | |
|---|---|---|---|
| *P. coronafaciens* | WP_122311533.1 | 60.25 | NA |
| *P. corrugata* | WP_175363759.1 | 59.18 | NA |
| *P. iridis* | WP_210703327.1 | 59.54 | NA |
| *P. lactucae* | WP_205491739.1 | 60.07 | NA |
| *P. zanjanensis* | WP_186709201.1 | 59.18 | NA |
| *P. baetica* | WP_221730730.1 | 59.00 | NA |
| *P. asplenii* | WP_102900200.1 | 59.25 | NA |
| *P. caricapapayae* | RMV94038.1 | 60.39 | NA |
| *P. rustica* | WP_212543876.1 | 59.25 | NA |
| *P. paraversuta* | WP_202210966.1 | 58.47 | NA |
| *P. migulae* | WP_084319057.1 | 59.00 | NA |
| *P. bubulae* | WP_130872039.1 | 58.82 | NA |
| *P. zeae* | WP_186625330.1 | 59.07 | NA |
| *P. saudimassiliensis* | WP_044501029.1 | 59.79 | NA |
| *P. cerasi* | WP_065350390.1 | 59.71 | NA |
| *P. iranensis* | WP_186567676.1 | 58.65 | NA |
| *P. petroselini* | WP_231807743.1 | 59.00 | NA |
| *P. bijieensis* | WP_249016042.1 | 58.82 | NA |
| *P. prosekii* | WP_121731231.1 | 58.47 | NA |
| *P. silesiensis* | WP_064679340.1 | 58.90 | NA |
| *P. meliae* | WP_044343430.1 | 59.22 | NA |

| | | | |
|---|---|---|---|
| *P. brenneri* | WP_090291800.1 | 58.82 | NA |
| *P. laoshanensis* | WP_149331581.1 | 60.11 | NA |
| *P. paralactis* | WP_057704236.1 | 58.82 | NA |
| *P. abyssi* | WP_096006403.1 | 58.90 | NA |
| *P. cannabina* | WP_055000596.1 | 58.65 | NA |
| *P. baltica* | WP_185793898.1 | 58.54 | NA |
| ***Halopseudomonas*** | | | |
| *H. pelagia* | WP_022964164.1 | 78.11 | NA |
| *H. salina* | WP_150278990.1 | 77.94 | NA |
| *H. oceani* | WP_104739548.1 | 77.78 | NA |
| *H. bauzanensis* | EZQ17079.1 | 77.68 | NA |
| *H. formosensis* | NLC00270.1 | 77.14 | NA |
| *H. pertucinogena* | WP_188635600.1 | 74.91 | NA |
| *H. aestusnigri* | WP_088275114.1 | 59.96 | NA |
| *H. sabulinigri* | WP_092283841.1 | 59.33 | NA |
| *H. gallaeciensis* | WP_118130125.1 | 58.90 | NA |
| *H. pachastrellae* | SFM16149.1 | 58.54 | NA |
| *H. xiamenensis* | WP_185266019.1 | 58.82 | NA |
| ***Azotobacter*** | | | |
| *A. salinestris* | WP_152386347.1 | 74.73 | LLGNHNLLITDPRDLPGMVREL |
| *A. chroococcum* | WP_165893083.1 | 73.12 | LLGNHNLLISDPRDLPGMVREL |

| | | | |
|---|---|---|---|
| *A. beijerinckii* | WP_090937998.1 | 73.84 | LTGNHNLLISDPRDLPAMVREL |
| *A. vinelandii* | WP_012701566.1 | 73.06 | LIGNHNLLITDPRDLPGMVRVL |
| ***Azomonas*** | | | |
| *A. macrocytogenes* | WP_183167613.1 | 71.71 | LIGAHNVLITNPRDLPAFIKDL |
| *A. agilis* | WP_144570445.1 | 67.21 | LIGSHSLLIVDPRDLSTLIRVL |
| ***Endozoicomonas*** | | | |
| *E. atrinae* | WP_066013753.1 | 65.89 | NA |
| *E. elysicola* | WP_026258079.1 | 65.89 | NA |
| *E. acropora* | WP_101746573.1 | 64.4 | NA |
| *E. arenosclerae* | WP_062268756.1 | 64.16 | NA |
| *E. ascidiicola* | WP_067519230.1 | 65.47 | NA |
| *E. montiporae* | WP_034876558.1 | 64.57 | NA |
| *E. numazuensis* | WP_034838388.1 | 63.39 | NA |
| ***Thalassolituus*** | | | |
| *T. marinus* | WP_225671693.1 | 64.34 | NA |
| *T. oleivorans* | WP_015486765.1 | 63.72 | NA |
| ***Marinobacter*** | | | |
| *M. salicampi* | WP_166259760.1 | 63.38 | NA |
| *M. goseongensis* | WP_248181088.1 | 63.20 | NA |
| *M. halotolerans* | WP_150913290.1 | 63.11 | NA |
| *M. segnicrescens* | WP_091850285.1 | 62.48 | NA |

| | | | |
|---|---|---|---|
| *M. confluentis* | WP_135801941.1 | 63.29 | NA |
| *M. daqiaonensis* | WP_092008766.1 | 63.38 | NA |
| *M. salarius* | WP_085680518.1 | 62.66 | NA |
| *M. changyiensis* | WP_152206729.1 | 63.44 | NA |
| *M. manganoxydans* | WP_008171819.1 | 63.18 | NA |
| *M. gudaonensis* | WP_091985593.1 | 63.44 | NA |
| *M. caseinilyticus* | WP_166268387.1 | 63.02 | NA |
| *M. algicola* | WP_007152576.1 | 62.66 | NA |
| *M. subterrani* | WP_048494758.1 | 62.72 | NA |
| *M. fuscus* | WP_106765131.1 | 63.36 | NA |
| *M. maroccanus* | WP_104321534.1 | 62.82 | NA |
| *M. nauticus* | WP_224848548.1 | 63.54 | NA |
| *M. adhaerens* | WP_041645242.1 | 62.82 | NA |
| *M. bryozoorum* | WP_248234290.1 | 62.12 | NA |
| *M. pelagius* | WP_091999381.1 | 62.23 | NA |
| *M. flavimaris* | WP_104271182.1 | 62.64 | NA |
| *M. salsuginis* | WP_136629430.1 | 62.64 | NA |
| *M. lipolyticus* | WP_213476103.1 | 62.39 | NA |
| *M. lutaoensis* | WP_076724125.1 | 61.51 | NA |
| *M. aromaticivorans* | WP_100688648.1 | 62.43 | NA |
| *M. zhejiangensis* | WP_092023867.1 | 62.21 | NA |

| | | | |
|---|---|---|---|
| *M. santoriniensis* | WP_008938484.1 | 62.30 | NA |
| *M. nitratireducens* | KEF32272.1 | 62.21 | NA |
| *M. nanhaiticus* | WP_004582476.1 | 62.30 | NA |
| *M. vulgaris* | WP_114612863.1 | 62.09 | NA |
| *M. daepoensis* | WP_029653343.1 | 62.72 | NA |
| *M. nitratireducens* | WP_036128814.1 | 62.21 | NA |
| *M. excellens HL-55* | KPQ30209.1 | 62.27 | NA |
| *M. halophilus* | WP_106673060.1 | 62.09 | NA |
| *M. salinexigens* | WP_149599508.1 | 61.78 | NA |
| *M. salexigens* | WP_100640529.1 | 62.21 | NA |
| *M. alexandrii* | WP_138441538.1 | 62.03 | NA |
| *M. zhanjiangensis* | WP_189577896.1 | 61.76 | NA |
| *M. perlucidum* | WP_114416312.1 | 64.94 | NA |
| *M. sediminum* | WP_203300677.1 | 62.03 | NA |
| *M. salinus* | WP_070966705.1 | 62.21 | NA |
| *M. oulmenensis* | WP_183700662.1 | 61.37 | NA |
| *M. litoralis* | WP_114333636.1 | 61.76 | NA |
| *M. antarcticus* | WP_072797248.1 | 61.84 | NA |
| *M. mobilis* | WP_091812100.1 | 61.12 | NA |
| *M. maritimus* | WP_144776680.1 | 61.48 | NA |
| *M. fonticola* | WP_148862619.1 | 61.40 | NA |

| | | | |
|---|---|---|---|
| *M. piscensis* | WP_144821906.1 | 60.00 | NA |
| *M. psychrophilus* | WP_048388949.1 | 60.94 | NA |
| *M. halodurans* | WP_131483374.1 | 61.12 | NA |
| *M. persicus* | WP_227663537.1 | 60.39 | NA |
| *M. gelidimuriae* | WP_026224292.1 | 61.12 | NA |
| ***Halomonas*** | | | |
| *H. utahensis* | WP_218668292.1 | 60.69 | NA |
| *H. heilongjiangensis* | WP_102629303.1 | 63.64 | NA |
| *H. ventosae* | WP_133635601.1 | 63.08 | NA |
| *H. korlensis* | WP_089792514.1 | 61.92 | NA |
| *H. nigrificans* | WP_096655061.1 | 63.74 | NA |
| *H. socia* | WP_163502500.1 | 62.96 | NA |
| *H. chromatireducens* | WP_066452348.1 | 61.62 | NA |
| *H. andesensis* | WP_126942923.1 | 62.31 | NA |
| *H. venusta* | WP_146945268.1 | 62.34 | NA |
| *H. humidisoli* | WP_095602254.1 | 62.34 | NA |
| *H. hydrothermalis* | WP_172420406.1 | 62.15 | NA |
| *H. pantelleriensis* | WP_089657799.1 | 61.85 | NA |
| *H. campisalis* | WP_238976403.1 | 61.71 | NA |
| *H. azerbaijanica* | TFH85557.1 | 61.11 | NA |
| *H. olivaria* | WP_249977216.1 | 62.80 | NA |

| | | | |
|---|---|---|---|
| *H. muralis* | WP_089724748.1 | 63.23 | NA |
| *H. beimenensis* | WP_097789621.1 | 62.52 | NA |
| *H. alkaliantarctica* | WP_160865853.1 | 62.29 | NA |
| *H. massiliensis* | WP_075879629.1 | 62.10 | NA |
| *H. zhangzhouensis* | WP_234273484.1 | 62.08 | NA |
| *H. halocynthiae* | WP_027967043.1 | 61.54 | NA |
| *H. campaniensis* | WP_038482926.1 | 62.34 | NA |
| *H. montanilacus* | WP_114479994.1 | 62.06 | NA |
| *H. pellis* | WP_149329101.1 | 61.87 | NA |
| *H. johnsoniae* | WP_193462082.1 | 61.60 | NA |
| *H. lysinitropha* | WP_151444583.1 | 61.48 | NA |
| *H. xianhensis* | WP_092844243.1 | 62.29 | NA |
| *H. stevensii* | WP_016914337.1 | 61.22 | NA |
| *H. aerodenitrificans* | WP_234254173.1 | 62.80 | NA |
| *H. glaciei* | WP_179917003.1 | 61.22 | NA |
| *H. neptunia* | WP_240718436.1 | 61.34 | NA |
| *H. stenophila* | WP_183381960.1 | 59.93 | NA |
| *H. meridiana* | WP_172416053.1 | 61.60 | NA |
| *H. kenyensis* | WP_181514132.1 | 62.62 | NA |
| *H. eurihalina* | WP_149320562.1 | 62.29 | NA |
| *H. titanicae* | WP_144815405.1 | 61.52 | NA |

| | | | |
|---|---|---|---|
| *H. lutescens* | WP_188638225.1 | 61.41 | NA |
| *H. ethanolica* | WP_234268252.1 | 62.24 | NA |
| *H. arcis* | WP_089708214.1 | 61.68 | NA |
| *H. zincidurans* | WP_031383005.1 | 62.10 | NA |
| *H. aquamarina* | WP_089675885.1 | 61.41 | NA |
| *H. piezotolerans* | WP_153842444.1 | 61.68 | NA |
| *H. lactosivorans* | WP_111412427.1 | 61.50 | NA |
| *H. xinjiangensis* | WP_043527140.1 | 61.54 | NA |
| *H. caseinilytica* | WP_064698711.1 | 61.91 | NA |
| *H. boliviensis* | WP_007113207.1 | 61.42 | NA |
| *H. cerina* | WP_183324041.1 | 60.19 | NA |
| *H. litopenaei* | WP_206937103.1 | 61.42 | NA |
| *H. saccharevitan* | WP_089850361.1 | 60.93 | NA |
| *H. qijiaojingensis* | WP_189466395.1 | 61.68 | NA |
| *H. sulfidoxydans* | WP_209538858.1 | 61.31 | NA |
| *H. subterranea* | WP_092830852.1 | 61.50 | NA |
| *H. pacifica* | WP_146804099.1 | 62.10 | NA |
| *H. elongata* | WP_013331874.1 | 61.73 | NA |
| *H. bachuensis* | WP_167118969.1 | 60.41 | NA |
| *H. antri* | WP_219792718.1 | 60.56 | NA |
| *H. anticariensis* | WP_016415842.1 | 62.24 | NA |

| | | | |
|---|---|---|---|
| *H. icarae* | NAW11434.1 | 60.26 | NA |
| *H. sulfidivorans* | WP_209477630.1 | 60.22 | NA |
| *H. smyrnensis* | WP_016856290.1 | 60.85 | NA |
| *H. ilicicola* | WP_072821794.1 | 61.16 | NA |
| *H. alimentaria* | WP_240894282.1 | 60.00 | NA |
| *H. fontilapidosi* | WP_183313285.1 | 60.48 | NA |
| *H. shengliensis* | SDO30167.1 | 59.89 | NA |
| *H. cupida* | WP_073433210.1 | 60.37 | NA |
| *H. sedimenti* | WP_180094573.1 | 60.49 | NA |
| *H. urmiana* | WP_138179835.1 | 61.05 | NA |
| *H. sulfidaeris* | WP_113270921.1 | 61.71 | NA |
| *H. halodenitrificans* | WP_245598342.1 | 59.85 | NA |
| *H. huangheensis* | WP_021820695.1 | 59.04 | NA |
| *H. rituensis* | WP_114487314.1 | 59.59 | NA |
| *H. organivoran* | WP_183386578.1 | 60.37 | NA |
| *H. halmophila* | WP_141317918.1 | 60.49 | NA |
| *H. zhuhanensis* | WP_160417975.1 | 59.41 | NA |
| *H. borealis* | WP_136247551.1 | 60.67 | NA |
| *H. niordiana* | WP_136253790.1 | 59.85 | NA |
| *H. malpeensis* | WP_227389229.1 | 60.00 | NA |
| ***Oceanospirillum*** | | | |

| | | | |
|---|---|---|---|
| *O. sanctuarii* | WP_086480429.1 | 60.50 | NA |
| *O. linum* | WP_078320405.1 | 59.07 | NA |
| *O. multiglobuliferum* | WP_078743680.1 | 60.32 | NA |
| ***Marinospirillum*** | | | |
| *M. celere* | WP_091958177.1 | 63.45 | NA |
| *M. alkaliphilum* | WP_072324599.1 | 64.87 | NA |
| *M. minutulum* | WP_051168020.1 | 61.71 | NA |
| *M. insulare* | WP_051610219.1 | 62.27 | NA |
| ***Reinekea*** | | | |
| *R. forsetii* | WP_215999193.1 | 61.62 | NA |
| *R. blandensis* | WP_008043928.1 | 59.46 | NA |
| *R. marinisedimentorum* | WP_132700829.1 | 58.00 | NA |
| ***Chromohalobacter*** | | | |
| *C. sarecensis* | WP_246969246.1 | 62.83 | NA |
| *C. nigrandesensis* | WP_246896569.1 | 62.08 | NA |
| *C. canadensis* | WP_246921457.1 | 61.52 | NA |
| *C. japonicus* | WP_246895188.1 | 61.68 | NA |
| *C. moromii* | WP_247639484.1 | 61.50 | NA |
| *C. marismortui* | TDU24908.1 | 60.97 | NA |
| ***Microbulbifer*** | | | |
| *M. rhizosphaerae* | WP_183458563.1 | 61.94 | NA |

| | | | |
|---|---|---|---|
| *M. aggregans* | WP_226667445.1 | 61.99 | NA |
| *M. agarilyticus* | WP_010132834.1 | 61.61 | NA |
| *M. variabilis* | WP_020414791.1 | 62.36 | NA |
| *M. elongatus* | WP_231757053.1 | 61.42 | NA |
| *M. hydrolyticus* | WP_161858158.1 | 61.64 | NA |
| *M. harenosus* | WP_138234782.1 | 61.80 | NA |
| *M. donghaiensis* | WP_234995036.1 | 61.61 | NA |
| *M. hainanensis* | WP_193165358.1 | 61.80 | NA |
| *M. yueqingensis* | WP_091512943.1 | 61.05 | NA |
| ***Aidingimonas*** | | | |
| *A. halophila* | WP_092571037.1 | 62.27 | NA |
| *A. lacisalsi* | WP_148254314.1 | 60.79 | NA |
| ***Marinomonas*** | | | |
| *M. atlantica* | WP_067098163.1 | 60.73 | NA |
| *M. aquimarina* | SBS28527.1 | 60.52 | NA |
| *M. fungiae* | WP_055463257.1 | 60.15 | NA |
| *M. gallaica* | WP_067036621.1 | 60.18 | NA |
| *M. ostreistagni* | WP_204147095.1 | 59.48 | NA |
| *M. rhizomae* | RBP85725.1 | 59.37 | NA |
| *M. shanghaiensis* | WP_111639654.1 | 58.70 | NA |
| *M. polaris* | WP_072837890.1 | 57.83 | NA |

| | | | |
|---|---|---|---|
| ***Salinicola*** | | | |
| *S. acroporae* | WP_110717964.1 | 61.05 | NA |
| *S. peritrichatus* | WP_110650383.1 | 60.11 | NA |
| *S. tamaricis* | WP_106420441.1 | 60.04 | NA |
| *S. corii* | WP_149437876.1 | 58.49 | NA |
| ***Cobetia*** | | | |
| *C. amphilecti* | WP_225347545.1 | 59.07 | NA |
| *C. crustatorum* | WP_144726529.1 | 59.63 | NA |
| ***Sinobacterium*** | | | |
| *S. norvegicum* | WP_237444816.1 | 57.57 | NA |
| *S. caligoides* | WP_123713848.1 | 56.84 | NA |
| **Others** | | | |
| *Atopomonas hussainii* | WP_074867227.1 | 78.65 | NA |
| *Denitrificimonas caeni* | WP_022964979.1 | 76.52 | NA |
| *Thiopseudomonas alkaliphila* | WP_053106538.1 | 74.51 | NA |
| *Aestuariirhabdus litorea* | WP_125015272.1 | 69.89 | NA |
| *Ventosimonas gracilis* | WP_068389457.1 | 69.00 | NA |
| *Zooshikella ganghwensis* | WP_027707937.1 | 66.73 | NA |
| *Spartinivicinus ruber* | WP_163833456.1 | 66.31 | NA |
| *Venatorbacter cucullus* | WP_228344254.1 | 64.70 | NA |
| *Oleiphilus messinensis* | WP_087464521.1 | 64.63 | NA |

| | | | |
|---|---|---|---|
| *Bacterioplanes sanyensis* | WP_094060881.1 | 64.52 | NA |
| *Hydrocarboniclastica marina* | WP_136548965.1 | 64.49 | NA |
| *Kistimonas asteriae* | WP_211824234.1 | 64.17 | LNTLFVGLMKNPDFLQLDFKSL |
| *Perlucidibaca piscinae* | WP_022955677.1 | 63.65 | NA |
| *Entomomonas moraniae* | WP_127162468.1 | 63.48 | NA |
| *Terasakiispira papahanaumokuakeensis* | WP_068997478.1 | 62.79 | NA |
| *Halospina denitrificans* | WP_133735935.1 | 62.68 | NA |
| *Simiduia agarivorans* | WP_015045484.1 | 62.22 | NA |
| *Hahella chejuensis* | WP_011396504.1 | 62.03 | NA |
| *Umboniibacter marinipuniceus* | WP_121877514.1 | 61.83 | NA |
| *Sansalvadorimonas verongulae* | WP_155159165.1 | 61.65 | NA |
| *Halovibrio salipaludis* | PAU80975.1 | 61.41 | NA |
| *Euryarchaeota archaeon* | MBE02978.1 | 61.15 | NA |
| *Parendozoicomonas haliclonae* | WP_087108240.1 | 61.11 | NA |
| *Acinetobacter baumannii* | SSU26913.1 | 60.96 | NA |
| *Lipotes vexillifer* | XP_007447258.1 | 60.78 | NA |
| *Natronospirillum operosum* | WP_135481310.1 | 60.73 | NA |
| *Gynuella sunshinyii* | WP_044618238.1 | 60.29 | NA |
| *Saccharospirillum impatiens* | WP_028669882.1 | 60.00 | NA |
| *Fluviicoccus keumensis* | WP_130412325.1 | 59.71 | NA |

| | | | |
|---|---|---|---|
| *Motiliproteus coralliicola* | RDE19752.1 | 59.59 | NA |
| *Maribrevibacterium harenarium* | WP_140588686.1 | 59.52 | NA |
| *Balneatrix alpica* | WP_027312460.1 | 58.55 | NA |
| *Agitococcus lubricus* | WP_107866136.1 | 57.07 | NA |

**Table S5.** Bacterial strains and plasmids used in this study

| Strain or plasmid | Phenotype and/or characteristic(s)[a] | Source or reference |
|---|---|---|
| ***P. aeruginosa*** | | |
| PAO1 | Wild-type strain of *P. aeruginosa* | Laboratory collection |
| Δ*fadD1* | Deletion mutant derived from PAO1 with *fadD1* being deleted | This study |
| Δ*fadD1*(*fadD1*) | Mutant Δ*fadD1* harboring the expression construct pBBRI-5-*fadD1* | This study |
| Δ*fadD1*(*fadD1$^{LM}$*) | Mutant Δ*fadD1* harboring the expression construct pBBRI-5-*fadD1$^{LM}$* (FadD1$^{LM}$: L275A L282A L289A L296A) | This study |
| Δ*fadD1*(*fadD2*) | Mutant Δ*fadD1* harboring the expression construct pBBRI-2-*fadD2* | This study |
| Δ*fadD1*(*fadD2$_{LZ}$*) | Mutant Δ*fadD1* harboring the expression construct pBBRI-2-*fadD2$_{LZ}$* (*fadD2* containing leucine zipper structure) | This study |
| Δ*dspI* | Deletion mutant derived from PAO1 with *dspI* being deleted | This study |
| Δ*dspI*(*dspI*) | Mutant Δ*dspI* harboring the expression construct pLAFR3-*dspI* | This study |
| Δ*dspI*(*fadD1*) | Mutant Δ*dspI* harboring the expression construct pBBRI-5-*fadD1* | This study |
| Δ*dspI*Δ*fadD1* | Deletion mutant derived from PAO1 with *dspI* and *fadD1* being deleted | This study |
| Δ*dspI*Δ*fadD1*(*fadD1*) | Mutant Δ*dspI*Δ*fadD1* harboring the expression construct pBBRI-5-*fadD1* | This study |
| Δ*dspI*Δ*fadD1*(*fadD2$_{LZ}$*) | Mutant Δ*dspI*Δ*fadD1* harboring the expression construct pBBRI-2-*fadD2$_{LZ}$* | This study |
| Δ*dspI*Δ*fadD1*(*fadD1$^{R454A}$*) | Mutant Δ*dspI*Δ*fadD1* harboring the expression construct pBBRI-2-*fadD1$^{R454A}$* | This study |
| Δ*dspI*Δ*fadD1*(*fadD1$^{G488A}$*) | Mutant Δ*dspI*Δ*fadD1* harboring the expression construct pLAFR3-*fadD1$^{G488A}$* | This study |

| | | |
|---|---|---|
| Δ*dspI*Δ*fadD1*(*fadD1*$^{R552A}$) | Mutant Δ*dspI*Δ*fadD1* harboring the expression construct pLAFR3-*fadD1*$^{R552A}$ | This study |
| PAO1(pBBRI-2-*egfp*) | PAO1 harboring the expression construct pBBRI-2-*egfp* | This study |
| Δ*fadD1*(pBBRI-2-*egfp*) | Mutant Δ*fadD1* harboring the expression construct pBBRI-2-*egfp* | This study |
| Δ*fadD1*(pBBRI-2-*fadD1-egfp*) | Mutant Δ*fadD1* harboring the expression construct pBBRI-2-*fadD1-egfp* | This study |
| PAO1(P*lasR-lacZ*) | PAO1 harboring the reporter construct P*lasR-lacZ* | This study |
| Δ*fadD1*(P*lasR-lacZ*) | Δ*fadD1* harboring the reporter construct P*lasR-lacZ* | This study |
| PAO1(P*lasI-lacZ*) | PAO1 harboring the reporter construct P*lasI-lacZ* | This study |
| Δ*fadD1*(P*lasI-lacZ*) | Δ*fadD1* harboring the reporter construct P*lasI-lacZ* | This study |
| PAO1(P*rhlR-lacZ*) | PAO1 harboring the reporter construct P*rhlR-lacZ* | This study |
| Δ*fadD1*(P*rhlR-lacZ*) | Δ*fadD1* harboring the reporter construct P*rhlR-lacZ* | This study |
| PAO1(P*rhlI-lacZ*) | PAO1 harboring the reporter construct P*rhlI-lacZ* | This study |
| Δ*fadD1*(P*rhlI-lacZ*) | Δ*fadD1* harboring the reporter construct P*rhlI-lacZ* | This study |
| PAO1(P*mvfR-lacZ*) | PAO1 harboring the reporter construct P*mvfR-lacZ* | This study |
| Δ*fadD1*(P*mvfR-lacZ*) | Δ*fadD1* harboring the reporter construct P*mvfR-lacZ* | This study |
| PAO1(P*pqsA-lacZ*) | PAO1 harboring the reporter construct P*pqsA-lacZ* | This study |
| Δ*fadD1*(P*pqsA-lacZ*) | Δ*fadD1* harboring the reporter construct P*pqsA-lacZ* | This study |
| PAO1(P*PA3298-lacZ*) | PAO1 harboring the reporter construct P*PA3298-lacZ* | This study |

| | | |
|---|---|---|
| Δ*fadD1*(P*PA3298-lacZ*) | Δ*fadD1* harboring the reporter construct P*PA3298-lacZ* | This study |
| PAO1(P*fadD2-lacZ*) | PAO1 harboring the reporter construct P*fadD2-lacZ* | This study |
| Δ*fadD1*(P*fadD2-lacZ*) | Δ*fadD1* harboring the reporter construct P*fadD2-lacZ* | This study |
| Δ*dspI*(P*lasR-lacZ*) | Δ*dspI* harboring the reporter construct P*lasR-lacZ* | This study |
| Δ*dspI*(P*lasI-lacZ*) | Δ*dspI* harboring the reporter construct P*lasI-lacZ* | This study |
| Δ*dspI*(P*rhlR-lacZ*) | Δ*dspI* harboring the reporter construct P*rhlR-lacZ* | This study |
| Δ*dspI*(P*rhlI-lacZ*) | Δ*dspI* harboring the reporter construct P*rhlI-lacZ* | This study |
| Δ*dspI*(P*mvfR-lacZ*) | Δ*dspI* harboring the reporter construct P*mvfR-lacZ* | This study |
| Δ*dspI*(P*pqsA-lacZ*) | Δ*dspI* harboring the reporter construct P*pqsA-lacZ* | This study |
| ***B. cenocepacia*** | | |
| H111 | Wild type strain, Genomovars III of the *B. cepacia* complex | Laboratory collection |
| Δ$rpfF_{Bc}$ | BDSF-minus mutant derived from H111 with $rpfF_{Bc}$ being deleted | Laboratory collection |
| H111(*fadD1*) | H111 harboring the expression construct pBBRI-5-*fadD1* | This study |
| H111(pBBRI-5-*mCherry*) | H111 harboring the expression construct pBBRI-5-*mCherry* | This study |
| ***Escherichia coli*** | | |
| DH5α | *supE44 lacU169(80lacZM15) hsdR17 recA1 endA1 gyrA96 thi-1 relA1 pir* | Laboratory collection |
| BL21 | *F-ompT hsdS (rB-mB-) dcm+ Tetr gal (DE3) endA* | Laboratory collection |
| **Plasmid** | | |
| pK18 | pK18, *sacB*$^+$; gene replacement vector, Kan$^r$ | Laboratory collection |

| | | |
|---|---|---|
| pK18-*fadD1* | pK18 containing fragments flanking *fadD1* | This study |
| pK18-*dspI* | pK18 containing fragments flanking *dspI* | This study |
| pBBRI-5 | Broad host range cloning vector, $Gm^r$ | Laboratory collection |
| pBBRI-5-*fadD1* | pBBRI-5 containing *fadD1* | This study |
| pBBRI-5-*fadD1*$^{LM}$ | pBBRI-5 containing *fadD1*$^{LM}$ ($FadD1^{LM}$: L275A L282A L289A L296A) | This study |
| pBBRI-5-*mCherry* | pBBRI-5 containing *mCherry* | This study |
| pBBRI-2 | Broad host range cloning vector, $Kan^r$ | Laboratory collection |
| pBBRI-2-*egfp* | pBBRI-2 containing *egfp* | This study |
| pBBRI-2-*fadD1-egfp* | pBBRI-2 containing *fadD1* and *egfp* | This study |
| pBBRI-2-*fadD2* | pBBRI-2 containing *fadD2* | This study |
| pBBRI-2-*fadD2*$_{LZ}$ | pBBRI-2 containing *fadD2*$_{LZ}$ (*fadD2* containing leucine zipper structure) | This study |
| pBBRI-2-*fadD1*$^{R454A}$ | pBBRI-2 containing *fadD1*$^{R454A}$ | This study |
| pLAFR3 | Broad host range cloning vector, $Tet^r$ | Laboratory collection |
| pLAFR3-*dspI* | pLAFR3 containing *dspI* | This study |
| pLAFR3-*fadD1*$^{G488A}$ | pLAFR3 containing *fadD1*$^{G488A}$ | This study |
| pLAFR3-*fadD1*$^{R552A}$ | pLAFR3 containing *fadD1*$^{R552A}$ | This study |
| pET21a | Expression vector, $Apm^r$ | Laboratory collection |
| pDBHT2 | Expression vector, $Kan^r$ | Laboratory collection |
| pET21a-*fadD1* | pET21a containing *fadD1* | This study |
| pET21a-*fadD1*$^{LM}$ | pET21a containing *fadD1*$^{LM}$ ($FadD1^{LM}$: L275A L282A L289A L296A) | This study |
| pET21a-AMP-B(*fadD1*) | pET21a containing AMP-binding enzyme domain of FadD1 | This study |

| | | |
|---|---|---|
| pET21a-AMP-B_C(*fadD1*) | pET21a containing AMP-binding enzyme C-terminal domain of the FadD1 | This study |
| pET21a-*fadD1*$^{W434A}$ | pET21a containing *fadD1*$^{W434A}$ | This study |
| pET21a-*fadD1*$^{T437A}$ | pET21a containing *fadD1*$^{T437A}$ | This study |
| pET21a-*fadD1*$^{R454A}$ | pET21a containing *fadD1*$^{R454A}$ | This study |
| pDBHT2-*fadD1*$^{G488A}$ | pDBHT2 containing *fadD1*$^{G488A}$ | This study |
| pDBHT2-*fadD1*$^{R552A}$ | pDBHT2 containing *fadD1*$^{R552A}$ | This study |
| pET21a-*fadD2* | pET21a containing *fadD2* | This study |
| pET21a-*fadD2*$_{LZ}$ | pET21a containing *fadD2*$_{LZ}$ (*fadD2* containing leucine zipper structure) | This study |
| pET28a | Expression vector, Kan$^{r}$ | Laboratory collection |
| pET28a-*fadD1*$_{Pf}$ | pET28a containing *fadD1*$_{Pf}$ (fatty acid degradation enzyme in *P. fluorescens* Migula ATCC17518) | This study |
| pME2-*lacZ* | Broad-host-range cloning vector, Tet$^{r}$ | Laboratory collection |
| P*lasR*-*lacZ* | pME2-*lacZ* containing the promoter of *lasR* | This study |
| P*lasI*-*lacZ* | pME2-*lacZ* containing the promoter of *lasI* | This study |
| P*rhlR*-*lacZ* | pME2-*lacZ* containing the promoter of *rhlR* | This study |
| P*rhlI*-*lacZ* | pME2-*lacZ* containing the promoter of *rhlI* | This study |
| P*mvfR*-*lacZ* | pME2-*lacZ* containing the promoter of *mvfR* | This study |
| P*pqsA*-*lacZ* | pME2-*lacZ* containing the promoter of *pqsA* | This study |

| | | |
|---|---|---|
| P*fadD2*-*lacZ* | pME2-*lacZ* containing the promoter of *fadD2* | This study |
| P*PA3298*-*lacZ* | pME2-*lacZ* containing the promoter of *PA3298* | This study |

[a] $Kan^r$, $Tet^r$, and $Gm^r$ indicate resistance to kanamycin, tetracycline, and gentamicin, respectively.

**Table S6.** PCR primers used in this study

| Primer | Sequence (5'-3') |
|---|---|
| For deletion | |
| *fadD1*L-F | CCGGAATTCCAAGGACGTCGATTTCTCCAAC |
| *fadD1*L-R | TTCCTGGATGATGAAGCCCACTCCTAAGCAACAG |
| *fadD1*R-F | GGAGTGGGCTTCATCATCCAGGAAGACGGCTAC |
| *fadD1*R-R | CCCAAGCTTAAAAATATTCGGAATTGTCGCA |
| *dspI*L-F | CCGGGATCCTTCGTCTATGCCATGAAAGG |
| *dspI*L-R | TGCGCCACTTGACGGCAGTGTTCATGAAGT |
| *dspI*R-F | CACTGCCGTCAAGTGGCGCAACTGCTGAG |
| *dspI*R-R | CCCAAGCTTGTCTGGGTACAGGCCGATGC |
| For *in trans* expression | |
| *fadD1*-F | CCCAAGCTTATGATCGAAAACTTCTGGAAGG |
| *fadD1*-R | CCGGAATTCGCAACGGCGGACTTACTTC |
| *fadD1*$^{LM}$L-R | CGTCGCGCGGGTTGGTGAT**TGC**GATGTTGTGGT TGCCGGTG**TGC**ATCAT |
| *fadD1*$^{LM}$R-F | AATCACCAACCCGCGCGAC**GCA**CCGTCGATGCT CAAGGAC**GCA**GGCCAG |
| *FadD2*-F | CCCAAGCTTATGCAACCTGAATTCTGGAA |
| *FadD2*-R | CCGGAATTCCAGGCGATTTCGCGCAGCT |
| *FadD2*$_{LZ}$L-R | GGTCGCGCGGGTTGGTGATCAGGATGTTGTGGT TGCCGGTGAGGGTGAAGGCATAGATG |
| *FadD2*$_{LZ}$R-F | GATCACCAACCCGCGCGACCTGCCGTCGATGCT CAAGGACCTCGCGAACTGCATGTGCA |
| *dspI*-F | TATGACCATGATTACGAATTCATGAACACTGCCG TCGAACC |
| *dspI*-R | ACGACGGCCAGTGCCAAGCTTTCAGCAGTTGCG CCACTTG |
| For protein expression | |
| FadD1-His-F | CCGGGATCCATGATCGAAAACTTCTGGAAGGAC |
| FadD1-His-R | CCCAAGCTTTTACTTCTGGCCCGCTTTC |
| FadD1$^{LM}$L-His-R | CGTCGCGCGGGTTGGTGAT**TGC**GATGTTGTGGT TGCCGGTG**TGC**ATCAT |

| | |
|---|---|
| FadD1$^{LM}$R-His-F | AATCACCAACCCGCGCGAC**GCA**CCGTCGATGCTCAAGGAC**GCA**GGCCAG |
| *fadD1*$^{W434A}$L-R | GGTCTTCAG**TGC**ACCATCGG |
| *fadD1*$^{W434A}$R-F | CCGATGGT**GCA**CTGAAGACC |
| *fadD1*$^{T437A}$L-R | GATATCGCC**TGC**CTTCAGCC |
| *fadD1*$^{T437A}$R-F | GGCTGAAG**GCA**GGCGATATC |
| *fadD1*$^{R454A}$L-R | GTCCTTCTT**TGC**GTCGACGA |
| *fadD1*$^{R454A}$R-F | TCGTCGAC**GCA**AAGAAGGAC |
| *fadD1*$^{G488A}$L-R | GTCGGGGAT**TGC**GATCGCGG |
| *fadD1*$^{G488A}$R-F | CCGCGATC**GCA**ATCCCCGAC |
| pDBHT2-*fadD1*$^{R552A}$-His-R | CTCGAGTGCGGCCGCAAGCTTTTACTTCTGGCCCGCTTTCTTCAGCTCTTCGTC**TGC**CA |
| pDBHT2-*fadD1*-His-F | TTTCAGGGCCATATGGGATCCATGATCGAAAACTTCTG |
| pDBHT2-*fadD1*-His-R | CTCGAGTGCGGCCGCAAGCTTTTACTTCTGGCCCGCTT |
| AMP-B(*fadD1*)-His-R | CCCAAGCTTTTCGTTCGGATACACATTGA |
| AMP-B_C(*fadD1*)-His-F | CCGGAATTCTCCGGCTTCAATGTGTATCC |
| FadD2-His-F | CCGGGATCCATGCAACCTGAATTCTGGAA |
| FadD2-His-R | CCCAAGCTTTCAGGCGATTTCGCGCAGCT |
| FadD2$_{LZ}$L-His-R | GGTCGCGCGGGTTGGTGATCAGGATGTTGTGGTTGCCGGTGAGGGTGAAGGCATAGATG |
| FadD2$_{LZ}$R-His-F | GATCACCAACCCGCGCGACCTGCCGTCGATGCTCAAGGACCTCGCGAACTGCATGTGCA |
| FadD1$_{Pf}$-His-F | CCGGGATCCATGAACGAAGACTTTTGGAA |
| FadD1$_{Pf}$-His-R | CCCAAGCTTTTACTTTTTCAGGCCCAGCT |
| For EMSA assay | |
| EMSA-*lasR*-F | CTGGGTATTCAGTTCGCATA |
| EMSA-*lasR*-R | CAGCTCAAGAAAACCGTCAA |
| EMSA-mutant *lasR*L-R | AGTACGATACAGAGGGTAAGCAAACGTTTA |
| EMSA-mutant *lasR*R-F | CTTACCCTCTGTATCGTACTAGGTGCATCA |
| EMSA-*lasI*-F | GCAGATATATAGGGAAGGGC |
| EMSA-*lasI*-R | CCGACCAATTTGTACGATCA |
| EMSA-*rhlI*-F | ATCATCCTGAGCATCTCCGA |

| | |
|---|---|
| EMSA-*rhlI*-R | TTCCAGCGATTCAGAGAGCA |
| EMSA-*rhlR*-F | GTCGGCGTTTCATGGAATTG |
| EMSA-*rhlR*-R | CCACAGCAAAAAGCCTCCGT |
| EMSA-*mvfR*-F | CTGAAACAGAGCCGTCATCC |
| EMSA-*mvfR*-R | GTTCACGTGATTCAGGTTAT |
| EMSA-*pqsA*-F | TGTAACGGTTTTTGTCTGGC |
| EMSA-*pqsA*-R | GGTCAGGTTGGCCAATGTGG |
| EMSA-*PA0692*-F | ATCTCATGCAGATGCCGGAG |
| EMSA-*PA0692*-R | GTGCGCACAATTGAACAGCG |
| EMSA-mutant *PA0692*L-R | AAAACGGCGCTATGAAATCTCGTTCTTTTC |
| EMSA-mutant *PA0692*R-F | AGATTTCATAGCGCCGTTTTTAACAGGTTG |
| EMSA-*flgM*-F | AGAAGCTGCGCCGACCGGT |
| EMSA-*flgM*-R | ATTCAGCCGGTTGAAGTCGA |
| EMSA-mutant *flgM*L-R | TCGAAGGTATTTGCCTAGCTTTTCGGCGAA |
| EMSA-mutant *flgM*R-F | AGCTAGGCAAATACCTTCGAGGTTTACAAC |
| EMSA-*katA*-F | CGTAGAAGCTGCCGAATAAG |
| EMSA-*katA*-R | GGCAGTGGTCAGGCGGGTCT |
| EMSA-mutant *katA*L-R | TAAGGATGAACCTACAATCCAATTTATTAA |
| EMSA-mutant *katA*R-F | GGATTGTAGGTTCATCCTTAACCTGCTTTT |
| EMSA-*lasR*$_{Pf}$-F | TATGACTCAAGCTGACTGGA |
| EMSA-*lasR*$_{Pf}$-R | TTCCAGCTCAAGAAAACCGT |
| For fluorescence | |
| EGFP-F | CCGCTCGAGATGGTGAGCAAGGGCGAGGAG |
| EGFP-R | CCCAAGCTTTCAAAGATCTACCATGTACAGCTCG |
| mCherry-F | CCGCTCGAGGTGAGCAAGGGCGAGGAGGA |
| mCherry-R | CCCAAGCTTCTTGTACAGCTCGTCCATGC |
| For RT-qPCR | |
| q16s-F | GCGCAACCCTTGTCCTTAGTT |
| q16s-R | TGTCACCGGCAGTCTCCTTAG |
| q*lasR*-F | GCCTTCATCGTCGGCAACTAC |
| q*lasR*-R | GCGCACCACTGCAACACTTC |
| q*lasI*-F | CCGTTTCGCCATCAACTCTG |
| q*lasI*-R | GATCATCATCTTCTCCACGCCTA |
| q*rhlR*-F | CTCCTCGGAAATGGTGGTCTGG |

| | |
|---|---|
| q*rhlR*-R | CGGAAAGCACGCTGAGCAAAT |
| q*rhlI*-F | CCATCCGCAAACCCGCTACA |
| q*rhlI*-R | TCACCGCCACCACCGAACTG |
| q*mvfR*-F | CCTGCGGGTGCTGCTGGATA |
| q*mvfR*-R | CGACGACGAACGCCTTGGTGTAG |
| q*pqsA*-F | CAAGGTGAATGGCCGCTGGGTG |
| q*pqsA*-R | CGGAAGGTTGTCGTGGTAGAGGGTGT |
| q*PA3298*-F | TTCGTTGCAGGCAGTCGG |
| q*PA3298*-R | AGTCGCTTGAGCATCTCGG |
| q*fadD2*-F | TATGGCCTCACCGAATGCTC |
| q*fadD2*-R | GACATTGAAGCCGGAGACCA |
| For reporter | |
| P*lasR*-F | CCCAAGCTTACCTATGCGCCGCCGTTG |
| P*lasR*-R | CCGGAATTCCAGCTCAAGAAAACCGTCAA |
| P*lasI*-F | CCCAAGCTTTTCGAACATCCGGTCAGCAA |
| P*lasI*-R | CCGGAATTCCCGACCAATTTGTACGATCA |
| P*rhlR*-F | CCCAAGCTTGCGCGCGTGGGATCTTCCTC |
| P*rhlR*-R | CCGGAATTCCCACAGCAAAAAGCCTCCGT |
| P*rhlI*-F | CCCAAGCTTTGGAACGAGGCTCGCGATTGG |
| P*rhlI*-R | CCGGAATTCTTCCAGCGATTCAGAGAGCA |
| P*pqsA*-F | CCCAAGCTTGTGTCCCGACCGGCGATTCC |
| P*pqsA*-R | CCGGAATTCGGTCAGGTTGGCCAATGTGG |
| P*mvfR*-F | CCCAAGCTTGATTCTAACCGCATAGGTCG |
| P*mvfR*-R | CCGCTCGAGGTTCACGTGATTCAGGTTAT |
| P*fadD2*-F | CCCAAGCTTGGAGATGTAGCTGCCCATGC |
| P*fadD2*-R | CCGCTCGAGGCGTTTGTCGTTCCAGAATT |
| P*PA3298*-F | CCCAAGCTTGAACATCCAGGTCGGCACCA |
| P*PA3298*-R | CCGCTCGAGCAACCGACTGCCTGCAACGA |
